# Engineering ERα degraders with pleiotropic ubiquitin ligase ligands maximizes therapeutic efficacy by co-opting distinct effector ligases

**DOI:** 10.1101/2024.06.09.595178

**Authors:** Anna Shemorry, Willem den Besten, Melinda M. Mulvihill, Curt J. Essenburg, Nicole Blaquiere, Tracy Kleinheinz, Elisia Villemure, Frank Peale, Gauri Deshmukh, Danilo Maddalo, Elizabeth Levy, Kebing Yu, Elizabeth A. Tovar, Emily Wolfrum, Robert A. Blake, Karthik Nagapudi, William F. Forrest, Steven T. Staben, Carrie R. Graveel, Wayne J. Fairbrother, Ingrid E. Wertz

**Author notes:** Current address: Lyterian Therapeutics, South San Francisco, CA 94080, USA. Current address: Amgen, Thousand Oaks, CA 91320, USA. Current address: Fuhong Biopharma, Inc., Shanghai 201206, China (Kebing Yu). Current address: Lycia Therapeutics, South San Francisco, CA 94080, USA. These authors contributed equally (Anna Shemorry, Willem den Besten).

## Abstract

Proximity-inducing compounds that modulate target protein homeostasis are an emerging therapeutic strategy [1]. While the inherent complexity of these bifunctional compounds poses challenges for rational design and bioavailability, their composition also provides opportunities to co-opt specific cellular proteins to maximize therapeutic impact. Here, we systematically evaluate the cellular efficacy, biophysical mechanisms, and therapeutic benefits of a series of bifunctional degrader compounds, that are all engineered with the Estrogen Receptor-alpha (ERα)-inhibitor endoxifen linked to different bioactive ubiquitin ligase ligands. Bifunctional ERα degraders that incorporate CRL4-CRBN-binding ligands promoted the most potent ERα degradation, whereas those incorporating either CRL2-VHL- or IAP-binding ligands maximized the depth of ERα degradation. Notably, ERα degraders containing pan-IAP antagonist ligands significantly decreased the proliferation of ERα-dependent cells relative to clinical-stage ERα-degraders, including the SERDs fulvestrant and GDC-9545 and the bifunctional degrader ARV-471. Mechanistic studies revealed that pan-IAP antagonist-based ERα degraders uniquely promote TNFα-dependent cell death, unlike the clinical-stage comparators. Remarkably, the pan-IAP antagonist-ERα-degraders co-opt distinct effector ligases to achieve dual therapeutic effects: they harness XIAP within tumor cells to promote ERα degradation, and activate cIAP1/2 within tumor and immune cells to induce TNFα that drives tumor cell death. Our studies demonstrate a broader concept that co-opting the discrete functions of a selected set of cellular effectors, while simultaneously modulating therapeutic target protein homeostasis, are dual strategies that can be leveraged to maximize the efficacy of induced proximity therapeutics.

## Introduction

Co-opting ubiquitin system machinery for therapeutic benefit is a promising strategy, as exemplified by the clinical success of retrospectively-discovered degrader compounds such as thalidomide analogs [2–4] and fulvestrant [5] as well as the progression of rationally engineered degrader compounds through clinical trials [6, 7]. The ubiquitin system is a primary conduit for regulated degradation of cellular proteins. Ubiquitination describes the covalent modification of protein substrates with ubiquitin, a 76-amino acid protein that, when assembled into polyubiquitin chains, binds the proteasome and directs substrate degradation. Ubiquitin ligases are the arbiters of ubiquitination, because they recruit both the target protein and cooperating enzymes to facilitate the reaction [8].

Ubiquitin ligases may be recruited to therapeutic targets via monovalent molecular glues or heterobifunctional degrader compounds; the latter comprises both target-binding and ubiquitin ligase-binding ligands that are joined by a chemical linker. This design facilitates target protein ubiquitination and subsequent proteasomal degradation via induced proximity with the ubiquitin ligase [9]. Advantages of degrader compounds relative to small-molecule inhibitors include sub-stoichiometric efficacy, catalytic activity, and the potential to degrade targets that are otherwise intractable to small-molecule inhibition. Indeed, these attributes are translating to promising clinical efficacy [1]. Since the original publications describing the design and cellular activity of heterobifunctional degraders [10, 11], additional cellular systems and enzymes have been harnessed to degrade or modify target proteins for therapeutic benefit, including the autophagy system [12, 13], cell surface proteins [14, 15], kinases [16], phosphatases [17], and other cellular effectors that regulate protein homeostasis in a proximity-dependent manner [18].

The therapeutic promise of induced proximity therapeutics could be enhanced by resolving some of the challenges inherent to this strategy. For example, most bifunctional small-molecule degraders fall outside the “Rule of Five” properties that describe the likelihood of oral bioavailability [19]. Furthermore, the design of bifunctional compounds is largely empirical, such that the target ligand, ligase ligand, linker, and attachment sites must all be optimized to identify the most potent, selective, and bioavailable compounds. As such, considerable effort has been invested in data curation and correlative analyses in order to identify ideal degrader compound properties, with the goal of streamlining the design of efficacious compounds [20]. Delivery methodologies [21, 22] and design strategies [23] have also been employed to improve the properties and overall efficacy of this promising class of compounds.

Another strategy that can be employed to maximize the therapeutic benefit of bifunctional degraders is to harness the biological activities of both the target ligand and the ubiquitin ligase ligand, in order to complement the therapeutic effect of target protein degradation. In this study, we systematically compare the cellular effects of bifunctional degrader compounds, which are all engineered with the same target-binding ligand and different biologically-active ligase ligands. We report a specific bioactive ligase ligand that can enhance the cellular efficacy of the bifunctional degrader compound beyond that driven by target degradation alone. Importantly, we perform cellular co-culture and in vivo studies to reveal that the therapeutic impact of this bifunctional degrader extends beyond tumor cells, by harnessing the activity of a different set of ubiquitin ligases within immune cells to drive tumor cell death.

## Results

### Bioactive ligase ligands differentially contribute to bifunctional ERα degrader potency

To systematically profile the cellular effects of different ubiquitin ligase ligands that are incorporated into heterobifunctional degraders, we selected ligands that are reported to regulate specific cellular pathways and targets. These can be categorized into compounds that bind multi-subunit or single-protein ubiquitin ligases (Fig. 1A). For the Cullin/RING ubiquitin ligase (CRL) complexes, we chose ligands that bind the substrate-binding subunits of CRL2-VHL or CRL4-CRBN ligases. VHL regulates HIFα transcription factors [24] and the VHL-binding ligand VH-032 antagonizes binding of endogenous CRL2-VHL substrates, the hydroxylated HIFα transcription factors. Thus VH-032 blocks HIFα ubiquitination and subsequent degradation, and thereby upregulates HIF-target genes [25–27] (Fig. 1A). The clinically-approved immunomodulatory drug (IMiD) pomalidomide (Fig. 1A) binds CRBN and functions as a molecular glue to recruit neosubstrates. The ubiquitination and subsequent degradation of these neosubstrates, which include IKZF1, IKZF3, and CK1α, compromise tumor cell viability [3, 4, 28, 29]. Importantly, co-degradation of neosubstrates along with the targeted substrate has been reported when IMiDs are incorporated in bifunctional degraders [30].

**Figure 1.**
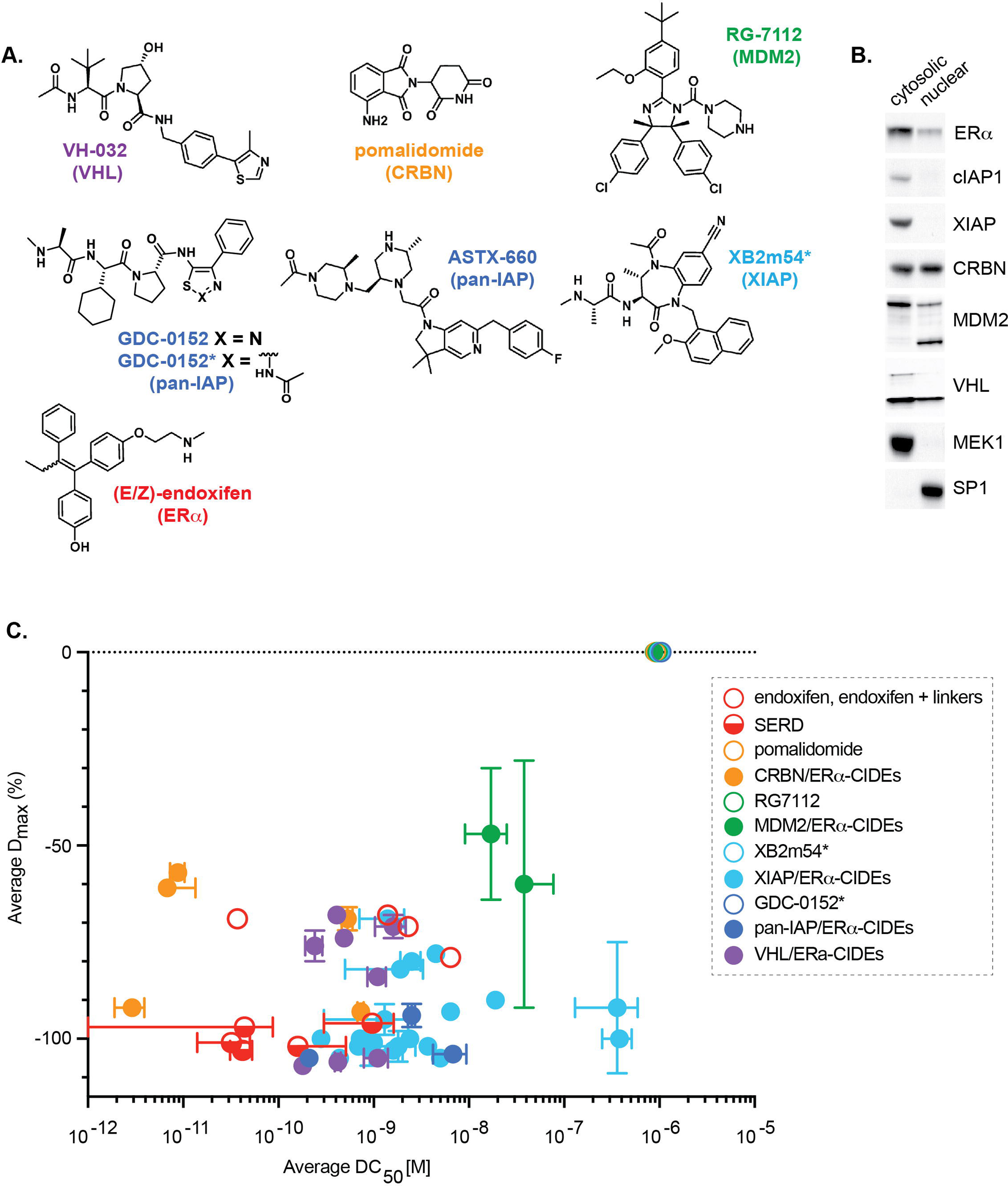
Bioactive ligase ligands differentially contribute to bifunctional ERα degrader potency. 1a. Chemical structures of the key E3 ligase ligands and ERL ligand used as standalone or as part of CIDEs in this study. Color code: VHL ligand = purple, CRBN ligand = orange, MDM2 ligand = green, pan-IAP ligands = dark blue, XIAP ligand = light blue, ERL ligand (E/Z-endoxifen) = red. The corresponding CIDEs match the color of their associated E3 ligase. 1b. E3 ligases co-opted by heterobifunctional degraders differ in their subcellular distribution. Lysates from MCF7 cells were separated into nuclear and cytosolic fractions and western blotted as indicated for E3 ligases or the relevant subunits. 1c. Graph comparing ERL degradation parameters DC_50_ (half maximal degradation concentration) and D_max_ (asymptotic maximum attainable degradation response) measured for various classes of ERL CIDEs, SERMs and SERDs. Changes in ERL protein levels in MCF7 cells were measured after 4 hour treatment with a range of concentrations of each compound and endogenous ERL protein levels measured using high content fluorescence imaging. Data represent mean DC_50_ and D_max_ with error bars representing standard deviation.

To co-opt single-protein ubiquitin ligases, we selected the MDM2 antagonist RG7112 ([31] and three different antagonists of inhibitor of apoptosis (IAP) ligases, GDC-0152 [32], an ASTX-660 analog [33], and XB2m54 and analogs [34, 35] (Fig. 1A). RG7112 binds MDM2 at the substrate binding site and thereby blocks recruitment, ubiquitination, and subsequent degradation of the p53 tumor suppressor. Elevated p53 levels promote cell cycle arrest, cell death, and tumor growth inhibition [36, 37]. GDC-0152 and ASTX-660 are “pan” IAP antagonists, because they bind the Baculoviral IAP Repeat-3 (BIR3) domain of multiple mammalian IAP family members, including X chromosome-linked IAP (XIAP), cellular IAP-1 (cIAP1), and cellular IAP-2 (cIAP2) that in turn promotes Tumor Necrosis Factor-α (TNFα)-dependent tumor cell death [38, 39]. XB2m54 preferentially binds the XIAP BIR2 domain, and thus inhibits nucleotide-binding oligomerization domain-2 (NOD2)-mediated inflammatory signaling [35]. RG7112, GDC-0152, and ASTX-660 have entered clinical trials given their promising anti-proliferative effects in tumor cells, and ASTX-660 (tolinapant) has recently been granted orphan drug designation for T-cell lymphomas [39–41].

We selected Estrogen Receptor-α (ERα) as our therapeutic target of interest, a nuclear hormone receptor that drives tumor cell proliferation in over 80% of breast cancer patients [42]. In addition to this clinical significance, our rationale for targeting ERα includes the following considerations: 1) high-affinity, therapeutically validated ERα antagonists are available for use as degrader ligands, 2) these ERα antagonists can be used as clinically-relevant controls to compare the cellular effects of ERα inhibition vs. ERα degradation, and 3) numerous selective ERα degraders (SERDs), including fulvestrant, GDC-810, GDC-927, and GDC-9545 are available as clinically-relevant comparators for ERα degradation [43–45]. We used the ERα inhibitor endoxifen (4-hydroxy-N-desmethyltamoxifen) as the target-binding ligand for our studies, given its enhanced pre-clinical efficacy relative to tamoxifen, and promising clinical trial data [46, 47] (Fig. 1A).

We first characterized the sub-cellular localization of ERα and the ubiquitin ligases, or the relevant ligase subunits, that are targeted by the above-mentioned small-molecule ligands (Fig. 1B). Western blot analysis of the ERα-dependent MCF-7 cell line revealed that certain ubiquitin ligases and subunits are localized to the cytosolic and/or nuclear fractions, whereas ERα is found in both compartments. The ligases cIAP2 and ML-IAP were poorly detected by western blot analysis in either cellular compartment in MCF-7 cells, likely due to lower expression. Thus, cIAP1, XIAP, CRBN, MDM2, and VHL are expressed and co-localize with ERα in MCF-7 cells, and thus are available to cooperate in bifunctional degrader-induced target ubiquitination.

Next, we synthesized endoxifen-based heterobifunctional degraders, here designated as chemical inducers of degradation (CIDEs), which incorporated the various ligase ligands described above (Fig S1A) and evaluated their effect on endogenous ERα protein levels by immunofluorescence detection. We also included SERDs, ERα antagonists, endoxifen conjugated with linker moieties, and free ligase ligands as controls (Table 1). ERα-CIDEs from our compound library that co-opt the IAP family of single-protein ligases or the CRL2-VHL ubiquitin ligase complex promoted the most complete degradation of endogenous ERα. Indeed, some ERα-CIDEs promoted comparable maximal ERα degradation (D_max_) as SERD clinical candidates, albeit with lower potency (DC_50_) (Fig. 1C). Representative ERα-CIDEs, SERDs, and control ligands were also evaluated by western blot analysis, which corroborated the immunofluorescence data (Fig. S1B-F, Table 1).

### IAP-based ERα degraders promote high-affinity ternary complexes

Ternary complex assembly between a target protein, degrader compounds, and the recruited ligase enables productive target ubiquitination for degradation, and can be quantitated by metrics such as the half-life (T_1/2_) and affinity (K_d_) measurements [30, 48, 49]. Varying the linker designs, ligase-binding ligand, target-binding ligand and linker attachment sites can either enhance or reduce ternary complex cooperativity. Systematic evaluation of ternary complex metrics across bifunctional degraders that recruit both single protein ligases and multi-subunit ligases to the same target protein has not been reported, nor have such data been correlated with degrader compound potency (DC_50_) and maximal target degradation (D_max_) measurements using untagged, endogenous target protein immunofluorescence. We therefore established ERα SPR-based binary and ternary complex binding assays to evaluate the relationships between CIDE-induced ERα DC_50_ and D_max_ values, with T_1/2_ and K_d_ ternary complex metrics.

To enable direct comparison between each ERα:CIDE:ligase ternary complex the common element, ERα, was immobilized onto an SPR sensorchip followed by the injection of each degrader compound to saturation (Fig. 2A). After optimizing the assay for each ligase, the ligase binding to the ERα:CIDE:ligase ternary complex was measured in dose response. VHL was co-expressed and purified with Elongin-B and -C to ensure optimal stability and a relevant conformational state; this complex has been reported to bind VH-032 with nM affinity [27]. We confirmed that XB2m54 binds XIAP-BIR2, and that ASTX660 and GNE-0152 bind XIAP-BIR3, all with nM binding affinities (Fig. S2A). Next, we selected XIAP-, pan-IAP, and VHL-based ERα-CIDEs having a range of maximal ERα degradation (D_max_) and potency (DC_50_) values, as evaluated by endogenous ERα immunofluorescence, for SPR analysis (Fig 2B, Fig. S2B, Fig 1C, Table 2).

**Figure 2.**
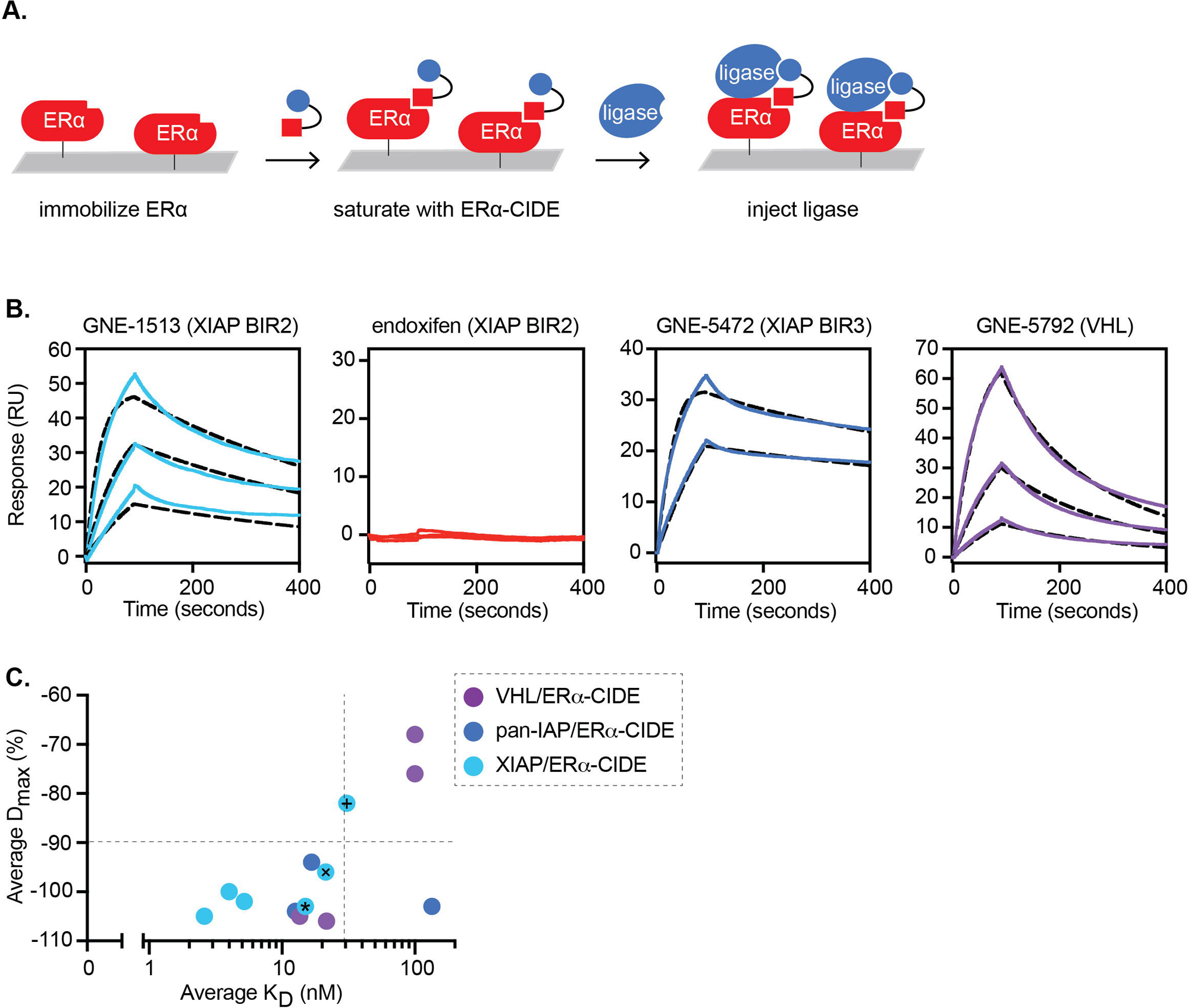
IAP-based ERα degraders promote high-affinity ternary complexes. 2a. Pictorial depiction of SPR ternary complex assessments involving CIDEs reported in this work. For all studies, ERL was immobilized to the SPR sensor chip followed by CIDE saturation. Ligase was then measured for binding to the ERL:CIDE complex in dose response. 2b. Representative SPR ternary complex assessments for ERL:CIDE binding to XIAP BIR2 (light blue), XIAP BIR3 (dark blue), VHL (purple), or negative control ligand endoxifen with XIAP BIR2 (red). Kinetic 1:1 fits shown in black dashed lines. 2c. Correlation of average ternary complex equilibrium binding constant (K_D_) measured by SPR with average maximum cellular degradation (D_max_) measured by ERL immunofluorescence for a set of ERL CIDEs paired to VHL (purple), XIAP BIR3 (dark blue), or XIAP BIR2 (light blue) is shown. Compounds marked with +, x. * belong to a series that contain the same ERL ligand (endoxifen) and linker and differ only in their XIAP BIR2 ligand. The R-squared K_D_ / D_max_ value for the 3 compound series is 0.994 and the R-squared K_D_ / cellular degradation potency (DC_50_) value for the 3 compounds is 0.960.

Our studies with VHL-, XIAP-BIR2-, and pan-IAP-BIR3-ligand-based ERα-CIDEs revealed that maximal ERα degradation correlates with higher affinity ternary complexes, and modestly correlates with more sustained ternary complexes (Fig. 2C, S2C, Table 2). More specifically, compounds that facilitate productive ERα degradation with a D_max_ of −90% or more form more stable complexes with measured binding affinities < 30 nM, and have half-lives > 30 seconds (Fig. 2C, S2C). ERα degradation potency (DC_50_) did not correlate with either ternary complex metric (data not shown). Interestingly, in this collection of compounds XIAP-BIR2 and pan-IAP-BIR3-based ERα degraders form more stable ternary complexes than VHL-based ERα degraders.

Because ternary complex half-life and affinity are two of many factors that determine cellular target degradation, deviations from the correlation may be due to compound properties including poor cellular permeability. To investigate this idea more thoroughly, we designed a series of XIAP-BIR2-based ERα-CIDE analogs that share the same endoxifen/linker component, but have variations of the XIAP-BIR2 ligand, that span a range of XIAP-BIR2 binding affinities. Importantly, these ERα-CIDE analogs have comparable cell permeability, as suggested by the similar cell potency difference by nanoBRET in digitonin-permeabilized versus intact cells (Fig S2D). These ERα-CIDE analogs demonstrate strong correlations between maximal ERα degradation, ternary complex affinity, and ternary complex half-life (Fig 2C, S2C). The ERα degradation potency (DC_50_) afforded by these CIDEs also strongly correlates with ternary complex affinity and ternary complex half-life (Table 2). These collective findings are corroborated by western blot analysis (Fig S2D). Thus, bifunctional ERα-CIDEs that promote longer-lived ternary complexes and have higher ternary complex affinity promote more complete and more potent cellular ERα degradation. These ternary complex data are consistent with previous reports on CRL2-VHL- and CRL4-CRBN-based BET and SMARCA heterobifunctional degraders [48, 50, 51] and extend the continuity of these principles from multi-subunit ubiquitin ligases to single-protein ligases, as correlated with the degradation of an endogenous, untagged protein target. Collectively, these data inform design principles to streamline development of potent and efficacious ERα degrader compounds.

### Pan-IAP antagonist-based ERα-CIDEs uniquely promote tumor cell death

Having identified and characterized potent ERα-CIDEs, our next goal was to compare the cellular effects of incorporating different biologically-active ligase-binding ligands within the bifunctional ERα degraders. We selected the most potent bifunctional ERα degraders based on the pan-IAP antagonist GDC-0152 (GNE-5472, designated as the pan-IAP/ERα-CIDE), the XIAP-selective antagonist XB2m54 (GNE-1513 or GNE-1567, designated as XIAP/ERα-CIDE), and the VHL-binding ligand VH-032 (GNE-5792, designated VHL/ERα-CIDE) (Fig. 1C, Fig. S1B, Table 1). We evaluated cell proliferation kinetics by monitoring NucLightRed-expressing MCF-7 cells via the Incucyte system, and included the ERα antagonist tamoxifen, the SERDs fulvestrant, GDC-810, GDC-927, and GDC-9545, and CRBN ligand-based ERα bifunctional degrader ARV-471 as clinical control compounds. The pan-IAP/ERα-CIDE was clearly the most potent inhibitor of cellular proliferation (Fig. 3A, S3A). Light microscopy revealed a cell death phenotype in cells treated with either GDC-0152 or the corresponding pan-IAP/ERα-CIDE. In contrast, tamoxifen, fulvestrant, XB2m54, and XIAP/ERα-CIDE clearly reduced cellular proliferation relative to DMSO-treated cells, but did not promote a cell death phenotype (Fig. 3B). Each of these bifunctional ERα degraders reduced ERα levels comparable to fulvestrant (Sup. Fig S3B). Thus the pan-IAP/ERα-CIDE has uniquely potent anti-proliferative activity in MCF7 cells, even compared to the clinical compounds GDC-0152, tamoxifen, fulvestrant, GDC-810, GDC-927, GDC-9545, and ARV-471.

**Figure 3.**
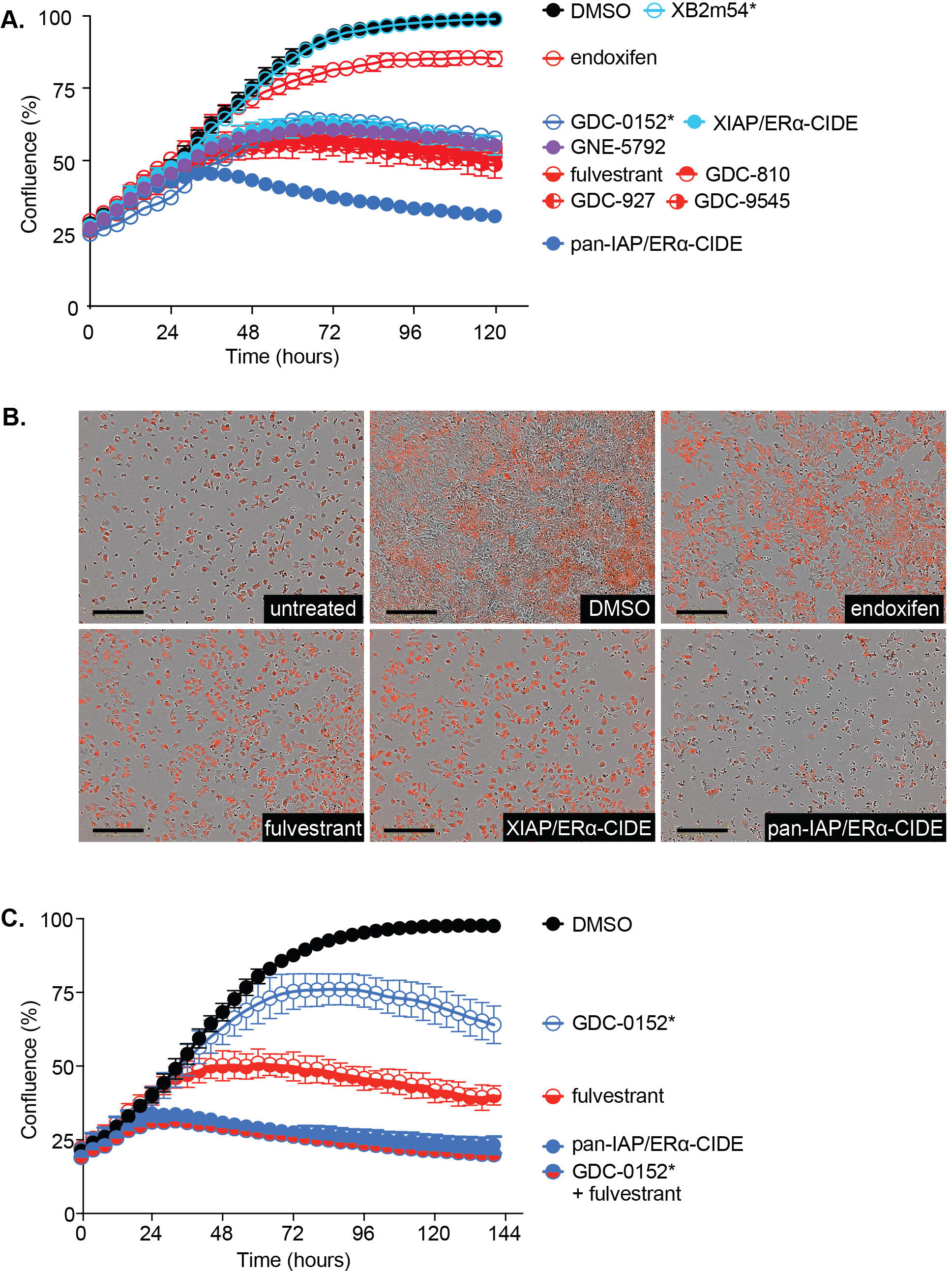
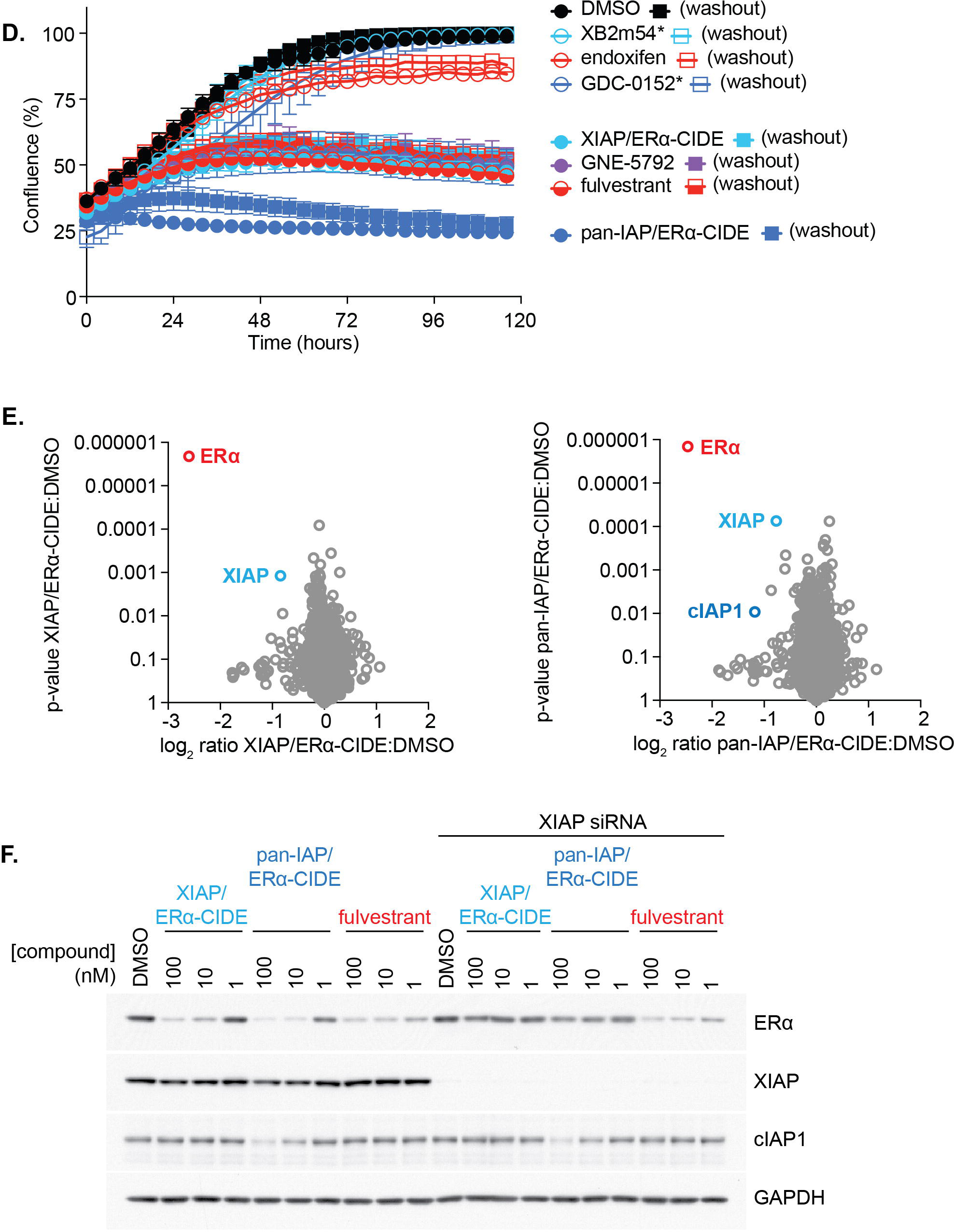
Pan-IAP antagonist-based ERα-CIDEs uniquely promote tumor cell death. 3a. Proliferation of MCF7 cells was monitored by confluence in an Incucyte instrument. Cells were treated as indicated with 250 nM of each compound. 3b. Images were captured from the Incucyte at the end of the experiment. 3c. The proliferation of MCF7 cells was monitored as in a. to demonstrate that the treatment with pan-IAP/ERα-CIDE is as effective as the combination of ERa inhibition with IAP inhibition. 3d. MCF7 cells were treated overnight with the indicated compounds and washed with media three times. Cells either continued to be treated as indicated or were grown in regular media. pan-IAP/ERα-CIDE demonstrated a robust inhibition of proliferation even after compound washout. 3e. Volcano plots of global proteomic profiling in the MCF7 cell line treated with either XIAP/ERα-CIDE or pan-IAP/ERα-CIDE compared to DMSO for 4 hours. The x axis corresponds to a log2 fold change and the y axis indicates p value. Highlighted in red is the statistically significant fold change of ERα compared to XIAP against the globally profiled proteome. 3f. Knockdown of XIAP rescues degradation of ERa induced by XIAP/ERα-CIDE and pan-IAP/ERα-CIDE. MCF7 cells were transfected with siRNA against XIAP and treated with compounds as indicated.

Culturing cells with a caspase-activated fluorescent dye revealed that both GDC-0152 and the pan-IAP/ERα-CIDE promoted caspase activation, consistent with the observed cell death phenotype and the reported mechanism of pan-IAP antagonists (Sup. Fig. S3C, S3D, [52, 53]. Interestingly, even though treatment with GDC-0152 alone or co-treatment of GDC-0152 with endoxifen ultimately activated caspases similarly to the pan-IAP/ERα-CIDE (Sup. Fig. S3E), both of these treatments were less efficacious than the pan-IAP/ERα-CIDE in limiting cellular confluency (Sup. Fig. S3F). This observation suggests that simultaneous ERα degradation and pan-IAP antagonism contribute to the enhanced efficacy of the pan-IAP/ERα-CIDE. To test this hypothesis, we co-treated cells with fulvestrant to promote ERα degradation and GDC-0152 to antagonize IAPs. This combination was as efficacious as the pan-IAP/ERα-CIDE, corroborating the idea that the enhanced efficacy of this engineered degrader is achieved via the dual effects of ERα degradation and pan-IAP antagonism (Fig. 3C).

Degrader compounds are reported to have enhanced activity relative to occupancy-driven inhibitors, owing to their catalytic mechanism of action [9]. We therefore performed washout studies to evaluate the benefits of sustained catalytic ERα degradation relative to occupancy-driven target inhibition. The pan-IAP/ERα-CIDE remained efficacious in proliferation studies after washout and also promoted sustained caspase activation after washout, in contrast to GDC-0152 (Fig. 3D, Sup Fig S3G). Our next goal was therefore to characterize the mechanistic basis of the enhanced potency of the pan-IAP/ERα-CIDE.

We chose the pan-IAP/ERα-CIDE and the XIAP/ERα-CIDE as comparators for additional studies: while both ERα-CIDEs engage IAP ligases, they have distinct effects on cell death induction, perhaps due to the specific IAP ligases recruited. Both ERα-CIDEs promoted comparable dose- and time-dependent ERα degradation (Fig S3H, S3I). The activity of both compounds was also dependent on the ubiquitin/proteasome system, as ERα degradation was blocked by the proteasome inhibitor bortezomib and the UAE1 inhibitor MLN7243. The NAE1 inhibitor had no effect on ERα degradation, as is expected, given that the IAP ligases are not neddylation-dependent (Fig S3J). The efficacy of both ERα-CIDEs is also dependent on the ERα binding function of endoxifen and their respective ligase ligands, as ligand competition studies blocked ERα degradation (Fig S3K, S3L). Notably, these mechanistic studies all revealed that cIAP1 levels are also depleted by treatment with the pan-IAP/ERα-CIDE, but not the XIAP/ERα-CIDE. This finding, along with the distinct cell viability effects that are induced by each ERα degrader, are consistent with the respective GDC-0152 and XB2m54 mechanisms of action [35, 54].

More specifically, the pan-IAP/ERα-CIDE induces cIAP1 degradation, caspase activation, and promotes cell death. As such, the pan-IAP/ERα-CIDE promotes ERα degradation while maintaining the pan-IAP antagonist function of GDC-0152. XIAP largely escapes autoubiquitination and degradation induced by both IAP-based ERα degraders, as revealed by western blot analysis (Fig S3H, S3I, S3K, S3L) and corroborating global proteomics analysis (Fig. 3E). Thus, XIAP is likely the primary ligase responsible for promoting sustained ERα ubiquitination and subsequent degradation in cells treated with the pan-IAP/ERα-CIDE or the XIAP/ERα-CIDE. Indeed, siRNA-mediated XIAP knockdown attenuated ERα degradation induced by both ERα-CIDEs, but not by fulvestrant, which is not XIAP-dependent (Fig. 3F). The XIAP dependence of ERα degradation corroborates a previous study describing an ERα bifunctional degrader, comprised of an LCL161 IAP antagonist derivative conjugated to 4-hydroxytamoxifen [55]. In sum, our collective data indicate that incorporating the pan-IAP antagonist GDC-0152 into an endoxifen-based ERα-CIDE decreases cellular proliferation beyond that achieved by ERα degradation or by pan-IAP antagonism alone. As such, the pan-IAP/ERα-CIDE more significantly and durably reduces cell viability in MCF-7 cells relative to the clinical compounds GDC-0152, tamoxifen, fulvestrant, GDC-9545, and ARV-471.

### The pan-IAP/ERα-CIDE drives tumor cell death via cIAP activation-induced autocrine or paracrine TNFα expression

Our next goal was to evaluate the efficacy of the pan-IAP/ERα-CIDE and the XIAP/ERα-CIDE in a broader panel of ERα-dependent breast cancer cell lines. Consistent with the MCF-7 data, efficient ERα depletion was induced by both ERα-CIDEs in the expanded panel of cell lines (Fig. 4A). Interestingly, the efficiency of ERα degradation did not always translate to potent inhibition of cellular proliferation: in the CAMA-1, T47D, HCC1428, and BT-474-M1 cell lines, the cellular proliferation effects of the pan-IAP/ERα-CIDE were no more potent than fulvestrant or the XIAP/ERα-CIDE (Fig. 4B). This proliferation effect is not attributable to a lack of XIAP or cIAP protein expression levels (Fig. S4A).

**Figure 4.**
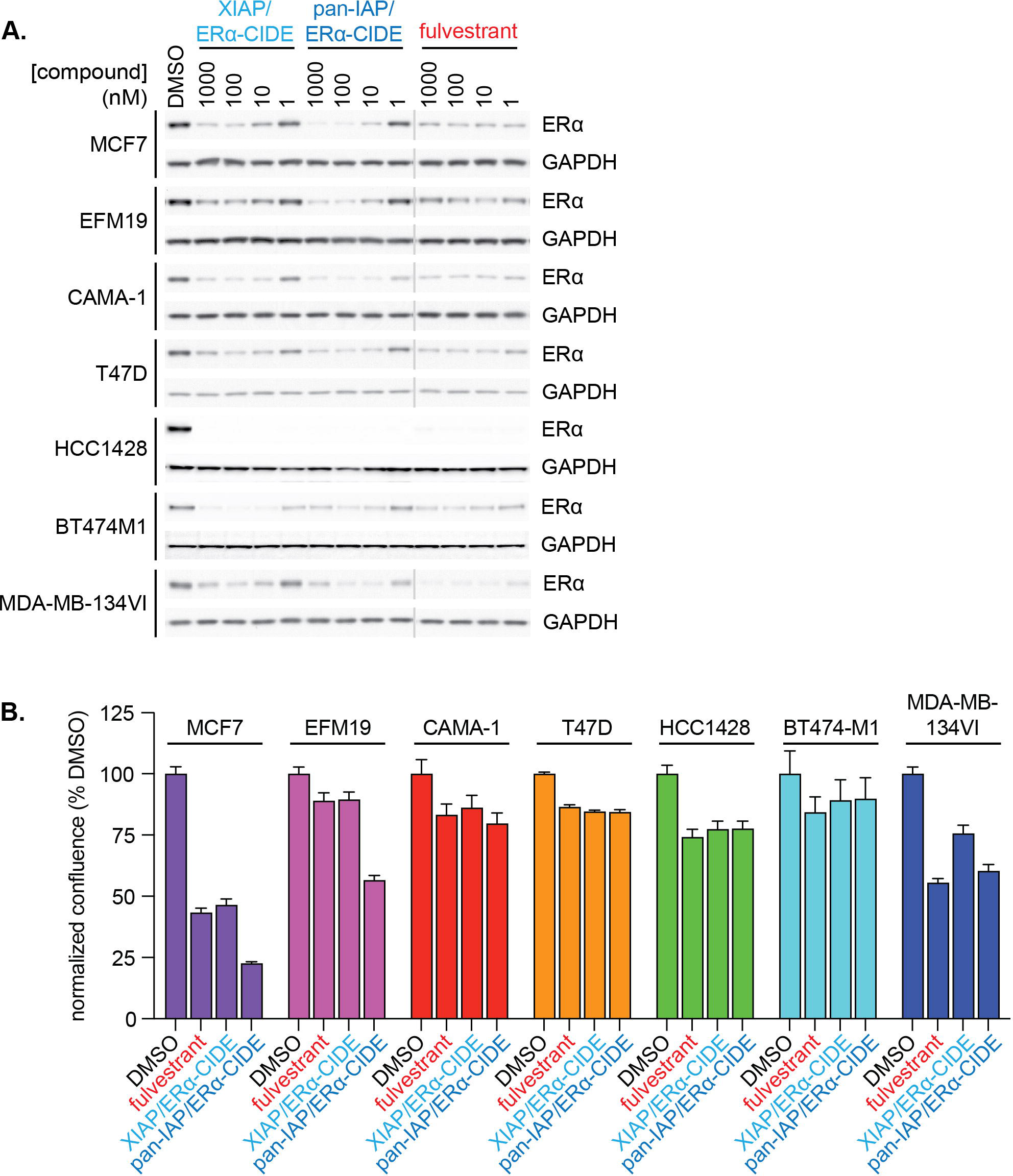
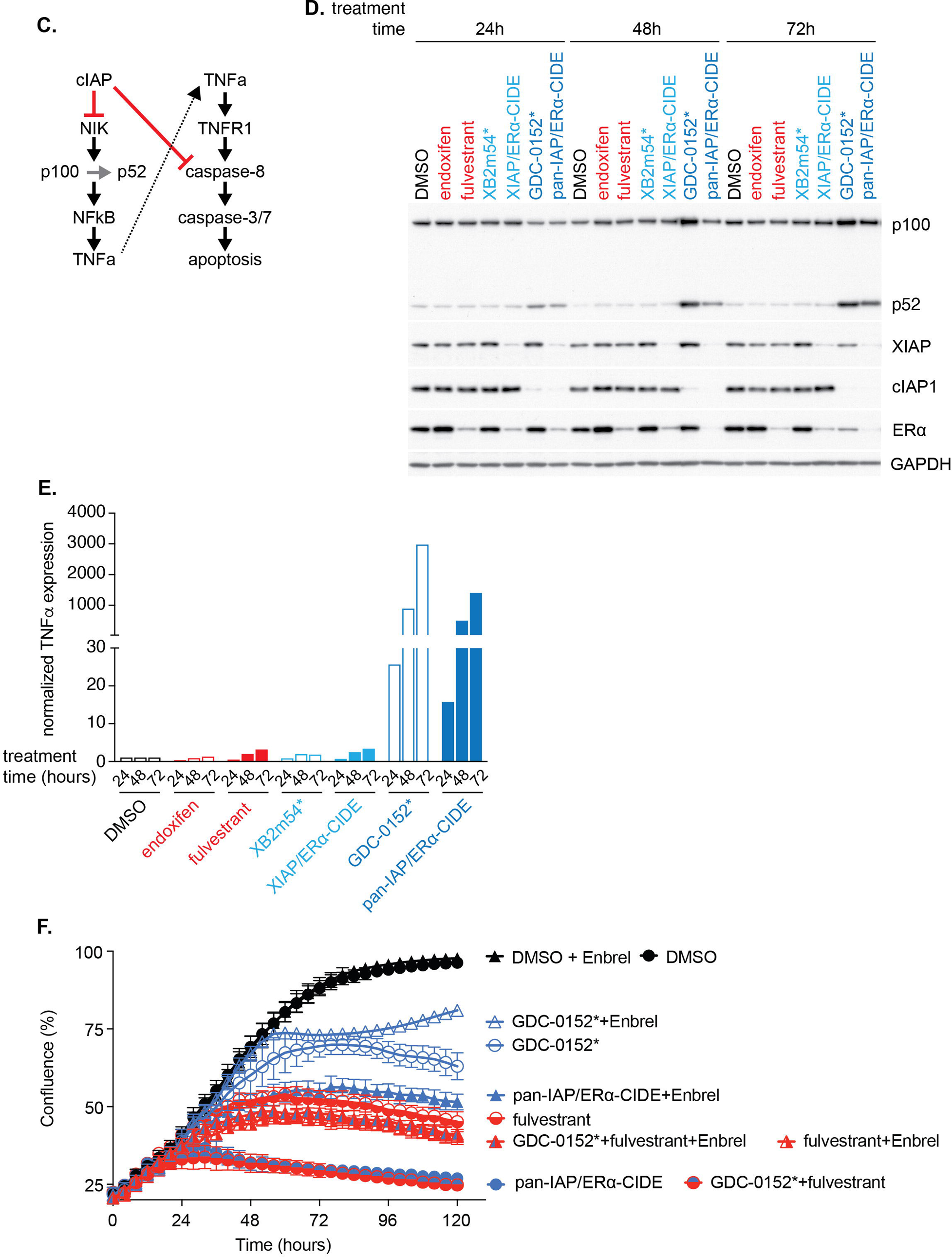
The pan-IAP/ERα-CIDE drives tumor cell death via cIAP activation-induced autocrine or paracrine TNFα expression. 4a. ER positive cell lines vary in ER degradation efficiency in response to XIAP/ERα-CIDE and pan-IAP/ERα-CIDE. 4b. Cell proliferation in response to different treatments was monitored in an Incucyte and the confluence normalized to DMSO after 144 hours was graphed. 4c. A simplified diagram of how cIAP influences both the non-canonical NFkB pathway and TNF production and TNF-TNFR1 signaling and apoptosis activation. 4d, e. GDC-0152 and pan-IAP/ERα-CIDE activate noncanonical NFkB demonstrated by the processing of p100 to p52 and activation of TNFa expression. MCF7 cells were treated with the indicated compounds for 24, 48, or 72 hours. Samples were either western blotted with the indicated antibodies or analyzed by qPCR for expression of TNFa. 4f. The addition of Enbrel to cells treated with pan-IAP/ERα-CIDE partially rescues cell proliferation defects. MCF7 cells were treated with 250 nM of the indicated compound with and without 10 ug/mL Enbrel. Cell proliferation was monitored using an Incucyte instrument.

It was therefore of interest to understand the physiological basis of the significant cell viability decrease induced by the pan-IAP/ERα-CIDE in only certain ERα-dependent breast cancer cell lines. We used the MCF-7 cell line as an initial model system for these studies, given the robust caspase activation response to the pan-IAP/ERα-CIDE (Fig S3C-S3E). Previous studies reported that pan-IAP antagonists activate non-canonical NFκB signaling [52, 53]. Mechanistically, monovalent pan-IAP antagonists activate cIAP1/2 autoubiquitination and rapid degradation. Loss of cIAP1/2 stabilizes the NIK kinase, and NIK-induced NFκB2 p100 phosphorylation promotes p100 proteasomal processing to p52. The p52 cleavage product heterodimerizes with RelB to activate non-canonical NFκB transcription, including enhanced TNFα expression (Fig. 4C). The induced expression of extracellular TNFα then activates TNF receptor-driven signaling in an auto- or paracrine fashion to promote caspase activation and apoptosis (Fig. 4C). Neither monovalent pan-IAP antagonists nor XIAP-selective ligands such as XB2m54 promote XIAP degradation or activation of cell death signaling. Indeed, both GDC-0152 and the pan-IAP/ERα-CIDE, but not XB2m54 or the XIAP/ERα-CIDE, promote p100 processing to p52 (Fig. 4D) and markedly enhance TNFα mRNA levels (Fig. 4E). Next, we co-treated MCF-7 cells with Enbrel, a fusion protein that neutralizes TNFα [56], in order to evaluate the role of TNFα in promoting caspase activation in this cellular system. While Enbrel co-treatment only modestly affected the proliferation of cells treated with DMSO or fulvestrant, Enbrel co-treatment clearly increased the confluence of cells treated with any GDC-0152-containing treatment regimen including GDC-0152 alone, GDC-0152 plus fulvestrant, or the pan-IAP/ERα-CIDE (Fig. 4F) and attenuated caspase activation (Fig. S4B). Importantly, in cells treated with the pan-IAP/ERα-CIDE or GDC-0152 plus fulvestrant, Enbrel co-treatment increased confluence to a similar level as cells treated with fulvestrant alone (Fig. 4F). Thus, in MCF-7 cells, the additional cellular proliferation decrease that is achieved by incorporating the pan-IAP antagonist ligand GDC-0152 within the ERα-CIDE is TNFα-driven and caspase-dependent.

A primary goal of our studies is to maximize the therapeutic benefit of heterobifunctional ERα degraders by incorporating biologically active ligase ligands that enhance cellular efficacy. As noted, the cell death effect of the pan-IAP/ERα-CIDE was not increased above ERα degradation alone in every evaluated ERα-dependent cell line (Fig. 4B). Because the cell death effect afforded by the pan-IAP/ERα-CIDE is TNFα-dependent in MCF-7 cells, we investigated the physiological effects of this ERα-CIDE on the expression of TNFα and of other cytokines and chemokines in the expanded panel of ERα-dependent breast cancer cell lines. We used the XIAP/ERα-CIDE that does not activate caspases (Fig S3C, S3D) or enhance TNFα expression (Fig 4E) in MCF-7 cells as a control. By quantitating cytokines and chemokines in a multiplexed immunoassay, we found that TNFα, IL6, and CXCL10 are significantly upregulated in both MCF-7 and EFM-19 cells in response to treatment with the pan-IAP/ERα-CIDE but not the XIAP/ERα-CIDE (Fig S5A, Extended Data Fig.1). Increased expression of TNFα, IL6, and CXLC10 are also correlated with decreased cell confluence in MCF-7 and EFM-19 cells in response to treatment with the pan-IAP/ERα-CIDE (Fig S5B). In parallel, we investigated cytokine and chemokine expression levels in peripheral blood mononuclear cells (PBMCs) after treatment with either ERα-CIDE, given that IAP antagonists are reported to regulate immune cell function and cytokine production [57]. The pan-IAP/ERα-CIDE, but not the XIAP/ERα-CIDE, significantly increased expression of six cytokines and chemokines, including TNFα, in PBMC cultures (Fig S5c).

Since we showed that cell death and caspase activation in MCF-7 cells treated with the pan-IAP/ERα-CIDE is TNFα-dependent, we reasoned that supplementing exogenous TNFα may provide the complementary milieu to foster the cell death phenotype in tumor cell lines that are deficient in TNFα production. To test this idea, we chose BT-474-M1 as the model cell line for our studies given the modest effects of the pan-IAP/ERα-CIDE on TNFα production (Fig S5A, Extended Data Fig.1) and on cell viability (Fig. 4B), despite robust ERα degradation (Fig. 4A). While TNFα co-treatment with DMSO or fulvestrant had modest effects on the rate of cell proliferation, treatment with TNFα in combination with GDC-0152 or the pan-IAP/ERα-CIDE markedly decreased cell confluence of BT-474-M1 cells (Fig. 5A). These data were corroborated by monitoring the kinetics of caspase activation, where co-treatment with TNFα with GDC-0152 or the pan-IAP/ERα-CIDE most profoundly antagonized BT-474-M1 proliferation or promoted caspase activation (Fig 5B). Next, we evaluated whether PBMCs, that are proficient in TNFα production in response to treatment with the pan-IAP/ERα-CIDE (Fig. S5C), can drive cell death via a paracrine mechanism in co-cultured BT-474-M1 cells. When BT-474-M1 cells were co-cultured with PBMCs, the pan-IAP/ERα-CIDE potently decreased cell confluence, an effect that was reversed with Enbrel co-treatment (Fig. 5C). Importantly, a control study revealed that Enbrel had little effect on the confluence of BT-474-M1 cells treated with DMSO, fulvestrant, XIAP/ERα-CIDE, or the pan-IAP/ERα-CIDE (Fig. S5E). Thus, efficacy of the pan-IAP/ERα-CIDE in ERα-dependent cell lines relies on both ERα depletion and the activation of TNFα-driven cell death pathways. Efficacy of the pan-IAP/ERα-CIDE is most profound in cell lines that produce autologous TNFα, such as MCF-7. Exogenous TNFα, either supplied as a recombinant cytokine or via co-culture with TNFα-producing immune cells, enables full efficacy in pan-IAP/ERα-CIDE-treated cell lines that do not produce sufficient TNFα to drive cell death.

**Figure 5.**
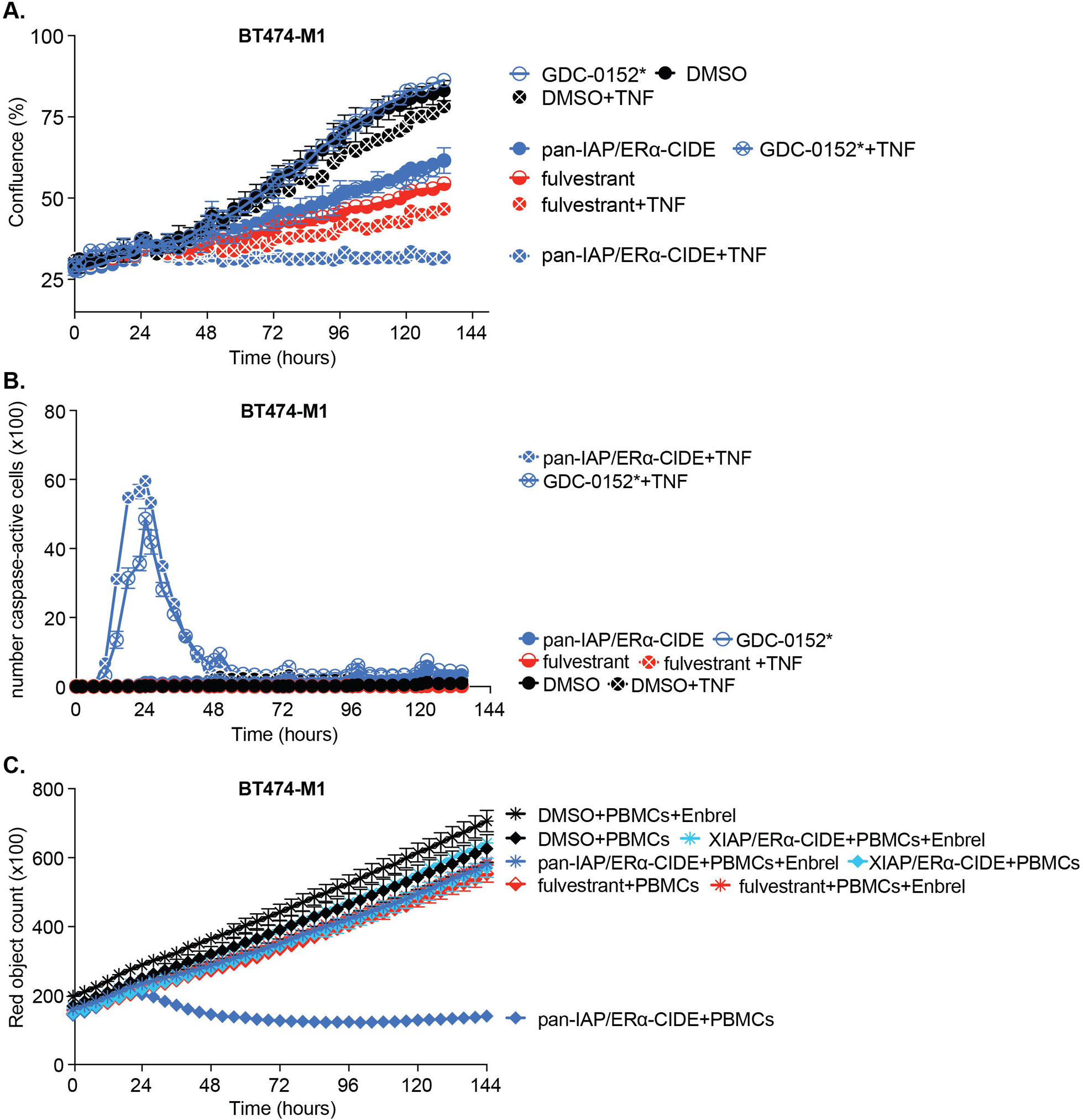
The pan-IAP/ERα-CIDE insensitive cell line, BT4741-M1, becomes sensitive to treatment in the presence of exogenous TNF alpha (TNF). 5a. BT4741-M1 cell proliferation was monitored after treatment with different compounds with and without 10 ng/mL amount of TNF. 5b. Caspase 3/7 was activated in BT4741-M1 cells treated with the combination of exogenous TNF-alpha and either GDC-0152 or pan-IAP/ERα-CIDE. BT4741-M1 cells were treated as in a. and a caspase activated fluorescent dye was monitored in an Incucyte instrument. 5c. Co-culturing BT4741-M1 with stimulated PBMCs attenuated cell proliferation in the presence of pan-IAP/ERα-CIDE. BT4741-M1 cells were grown in the presence or absence of stimulated PBMCs and treated with the indicated compounds. Cell proliferation was monitored using an Incucyte instrument that measured confluence.

### The pan-IAP/ERα-CIDE achieves tumor growth inhibition in a TNFα-dependent manner

Our findings raise the possibility of effecting maximal in vivo efficacy of the pan-IAP/ERα-CIDE by co-opting local tumor cell XIAP ligases to direct ERα degradation, and by activating systemic immune cell cIAP ligases to enhance TNFα expression and drive tumor cell death. We prepared four formulations of the pan-IAP/ERα-CIDE and evaluated the bioavailability in female C57BL/6 mice over 24 hours. The polymer-nanoparticle (PNP) formulation enabled a C_max_ and sustained serum concentration of the pan-IAP/ERα-CIDE above 1µM to afford the highest overall exposure relative to the other formulations (Fig S6A). This formulation also maintained an in vitro DC_50_ of 0.51nM (Fig S6A, inset) and was thus selected for further study. Next, we performed tumor growth inhibition studies in female NSG mice xenografted with MCF7 tumors (Fig S6B). The PNP formulation of the pan-IAP/ERα-CIDE was generally well-tolerated based on body weight measurements (Fig S6C), however only modest to insignificant tumor growth inhibition was achieved with any dosing regimen in this experimental model (Fig S6B).

Because we established that efficacy of the pan-IAP/ERα-CIDE relies on both ERα depletion and the activation of TNFα-driven cell death pathways within tumor cells, we reasoned that the immunocompromised state of NSG mice might preclude a full TNFα-driven response, thus contributing to the modest tumor growth inhibition. To investigate the impact of the pan-IAP/ERα-CIDE in an immunocompetent *in vivo* model of ERα-dependent breast cancer, we utilized an *Nf1*-deficient CRISPR rat model in which 100% of female rats develop aggressive and heterogeneous mammary adenocarcinomas within 14 weeks (Fig S6D) [58]. Indeed, several studies have demonstrated that the tumor suppressor *NF1* is a genetic driver of sporadic breast cancer, endocrine resistance, and inherited breast cancer [59–64]. The role of NF1 in negatively regulating RAS GTPase activity is established, and recent studies have also revealed that the NF1 protein, neurofibromin, interacts with ERα and modulates ERα transcription [58, 65]. *NF1* mutations are one of the most common mutations accrued in endocrine-resistant tumors, suggesting a key role of neurofibromin in modulating response to endocrine treatments [61, 62]. Resistance mechanisms in breast cancer due to *NF1* mutation likely occur through altered direct and indirect interactions with ERα [36, 58, 65–67]. Exons 21-22 within the cysteine-serine rich domain are targeted in the *Nf1*-deficient rat model, a modification that impacts both RAS and ERα interactions [68, 69]. Additionally, the strong estrogen dependence of *Nf1*-deficient tumors was demonstrated by the rapid tumor decrease that occurs after ovariectomy [58]. This unique *Nf1* rat model is invaluable for interrogating the role of estrogen-dependent breast cancer in an immunocompetent environment that demonstrates underlying endocrine-resistance signatures [58].

Furthermore, the rat mammary gland structure and immune system in the *Nf1*-deficient rat model more closely resemble humans than mouse mammary models and is the preferred toxicology model to use in preclinical studies [70, 71]. To evaluate the effect of the pan-IAP/ERα-CIDE on regulating rat tumor ERα expression levels, we developed an immunohistochemical H-scoring protocol and noted a dose-dependent reduction in endogenous ERα staining (Fig 6A). The 7.5 mg/kg dose was chosen for tumor growth inhibition studies given the significant decrease in ERα levels within tumors (Fig 6A, 6B), tolerability, and sustained serum exposure of the pan-IAP/ERα-CIDE (Fig S6E) that enabled a once-daily dosing protocol. The pan-IAP/ERα-CIDE is >99.9% bound to the plasma proteins, and the free drug concentration, assuming plasma and serum binding are similar, was 0.187 nM at 7.5 mg/kg, that is comparable to the *in vitro* DC_50_ (Fig 1C, S6A).

**Figure 6.**
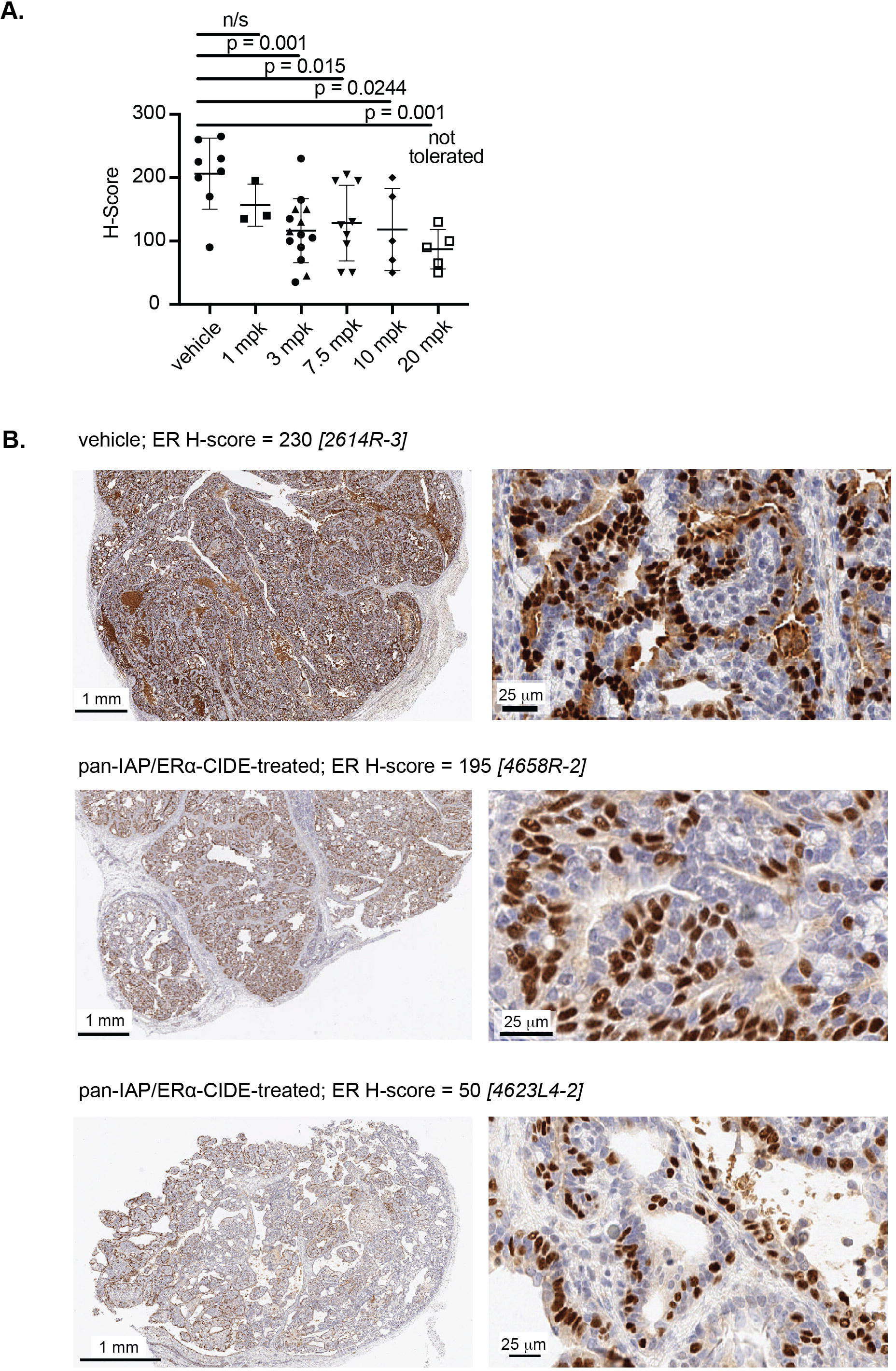
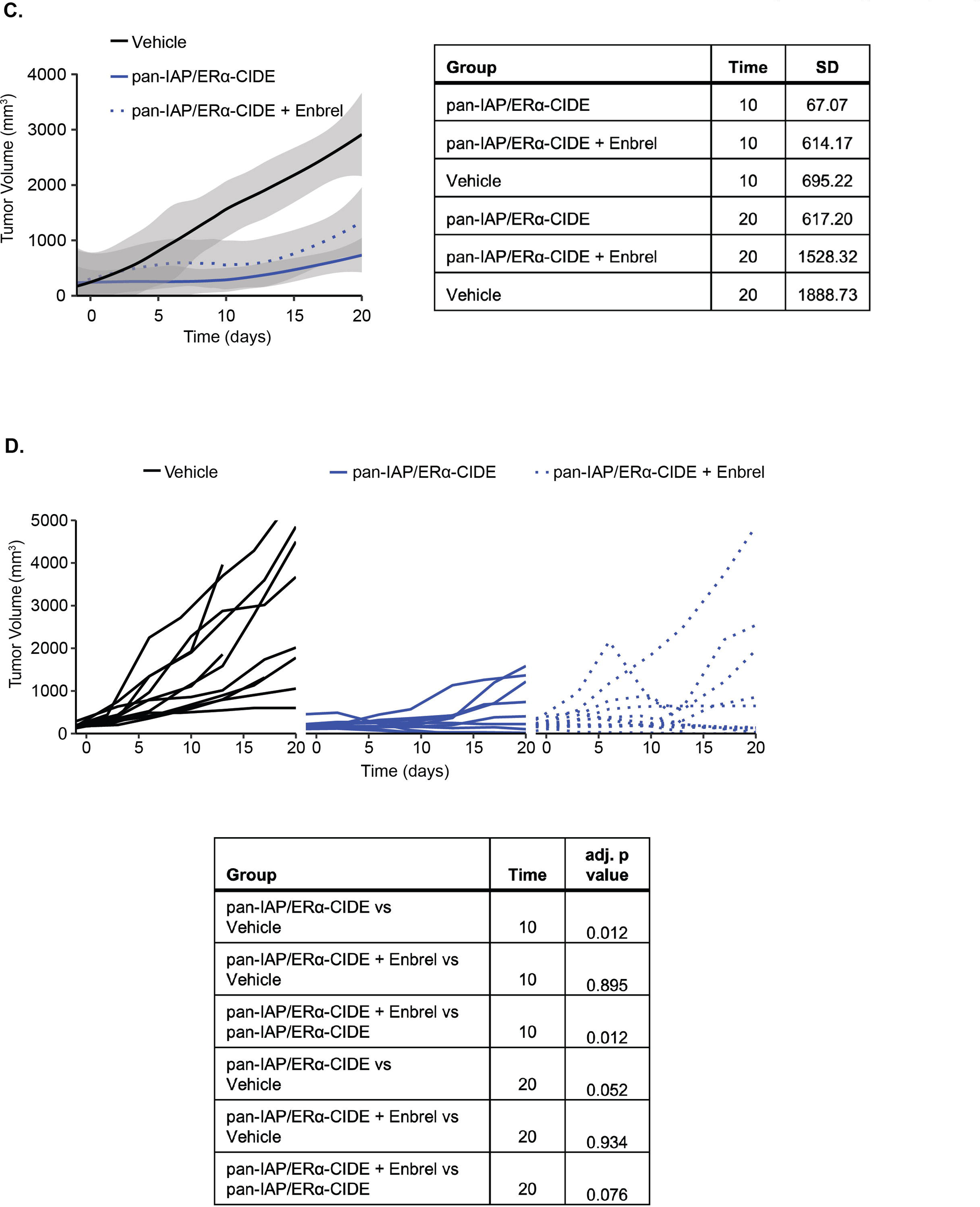
The pan-IAP/ERα-CIDE achieves tumor growth inhibition in a TNFα-dependent manner. 6a. Semi-quantitative evaluation of ER expression in tumor epithelial cells shows reduced signal relative to control tumors in animals treated with pan-IAP/ERα-CIDE at ≥ 3 mg/kg. 6b i. Immunostaining of tumors from vehicle-treated animals shows moderate to strong ER signal in nuclei of most glandular epithelial cells; ii., iii. Tumors from animals treated with 7.5 mg/kg pan-IAP/ERα-CIDE show reduced intensity and frequency of ER signal in glandular epithelial cell nuclei. 6 c, Treatment of mammary tumor bearing NF1 deficient rats with 7.5 mg/kg of pan-IAP/ERα-CIDE decreased tumor volume as compared to the vehicle. Co-treatment with 10 mg/kg rodent-like Enbrel partially reverted the tumor growth phenotype. The graph depicts the average tumor volume within each treatment group over time. 6 d. Graphs of individual tumor volumes demonstrate the amount of variability within both the 7.5 mg/kg pan-IAP/ERα-CIDE + Enbrel and vehicle treatment groups.

The pan-IAP/ERα-CIDE significantly inhibited tumor growth in the *NF1-*deficient ERα-dependent mammary tumor model relative to the vehicle control (Fig 6C). Analysis of serum and tumor samples confirmed exposure of the pan-IAP/ERα-CIDE in treated animals (Fig S6E), and kinetic body weight analysis indicated good tolerability (Fig S6F). To evaluate whether tumor growth inhibition in this model was TNFα-dependent, we dosed a cohort of pan-IAP/ERα-CIDE-treated animals with rodent-like Enbrel. Antagonizing TNFα attenuated tumor growth inhibition (Fig 6C) and increased the variability of response (Fig 6D). Collectively, these in vivo data corroborate our in vitro studies, and demonstrate that tumor growth inhibition by the pan-IAP/ERα-CIDE is driven by ERα depletion and the pan-IAP antagonist-induced activation of TNFα-activated cell death pathways.

## Discussion

Our primary research aim was to investigate whether we could maximize the therapeutic benefit of bifunctional degrader compounds by harnessing the biological activities of both the target ligand and the ubiquitin ligase ligand. We therefore systematically evaluated the efficacy of ERα-CIDEs synthesized with bioactive ligase ligands that co-opt either CRL2-VHL, CRL4-CRBN, MDM2, XIAP, or pan-IAP (cIAP1/2 and XIAP) ubiquitin ligase enzymes. A bifunctional degrader comprised of the pan-IAP antagonist GDC-0152, linked to the ERα inhibitor endoxifen, achieved the most potent growth inhibition of ERα-dependent tumor cells evaluated in our studies.

Our comprehensive mechanistic studies defined three key features comprising the basis of the pan-IAP/ERα-CIDE efficacy. First, the pan-IAP/ERα-CIDE dramatically decreased endogenous ERα protein levels: the depth of ERα degradation (D_max_) was comparable to or greater than clinical SERDs and ARV-471, while achieving picomolar degradation potency (DC_50_) in MCF7 cell culture studies. Biophysical analyses revealed that the high affinity of the ERα:pan-IAP/ERα-CIDE:IAP ligase ternary complex drives this ERα degradation efficacy. Second, the pan-IAP/ERα-CIDE uniquely activates tumor cell death pathways. This therapeutic effect complements the physiological consequences of ERα degradation. Notably, neither ARV-471 nor the clinical SERDs evaluated in our studies activate tumor cell death. Third, and most importantly, the pan-IAP/ERα-CIDE drives tumor growth inhibition via systemic effects that extend beyond tumor cells: the incorporated GDC-0152 ligand also promotes TNFα production by immune cells. TNFα then functions in an autocrine and paracrine fashion to activate tumor cell death pathways that, in cooperation with ERα depletion, activate the tumor cell death response. This finding raises the possibility that the efficacy of clinical ERα degrader compounds might also be enhanced by direct activation of tumor cell death pathways.

The therapeutic mechanisms of the pan-IAP/ERα-CIDE are related to, but distinct from, those achieved by certain clinical-stage degrader compounds including the thalidomide analog lenalidomide and the bifunctional degrader NX-2127. More specifically, lenalidomide and related analogs act as molecular glues that co-opt the CRL4-CRBN ligase to promote the degradation of neosubstrates such as IKZF1, IKZF3, and CK1α, that are essential for tumor cell viability. Lenalidomide and analogs also have immunomodulatory effects beyond tumor cells, including T-cell co-stimulation via induced IL-2 expression and IKZF1/3 degradation in T-cells, that activate a tumor immune response [72]. The bifunctional degrader NX-2127 is comprised of a BTK inhibitor linked with a thalidomide analog. NX-2127 harnesses CRL4-CRBN to degrade BTK via engineered induced proximity, while simultaneously promoting IKZF1/3 degradation in T-cells via the thalidomide analog to boost tumor immunity [73]. Thus, lenalidomide and NX-2127 promote the degradation of essential neosubstrates within tumor cells and simultaneously promote IKZF1/3 degradation in T-cells–distinct and complementary therapeutic effects that are both dependent on the CRL4-CRBN ligase. With these mechanisms in mind, it will be interesting to investigate whether ARV-471, and other bifunctional degraders that co-opt the CRL4-CRBN ligase for tumor-dependent target degradation, also boost tumor immunity via T-cell modulation.

Like the thalidomide analogs and NX-2127, the pan-IAP/ERα-CIDE also promotes tumor-localized and immune cell target degradation to achieve full therapeutic efficacy. However, the distinct cellular responses induced by the pan-IAP/ERα-CIDE are the result of harnessing different ubiquitin ligase enzymes. The tumor-centric efficacy of the pan-IAP/ERα-CIDE is due to XIAP-mediated ERα degradation, whereas the autocrine and paracrine TNFα-induced tumor cell death is dependent on cIAP1/2 activation, autoubiquitination, and subsequent degradation. The identification of other classes of cellular effectors that regulate protein homeostasis in a proximity-dependent manner [18] raises the possibility that the development of pleiotropic effector ligands, that are specifically engineered to harness a focused set of cellular effector proteins, may enable the rational design of induced proximity therapeutics with maximal therapeutic efficacy.

These collective features point to chemical design considerations for enhancing the efficacy of bifunctional degrader compounds. Indeed, co-opting the function of a selected set of cellular effectors, while simultaneously modulating therapeutic target protein homeostasis, are dual strategies that can be intentionally harnessed to maximize the efficacy of all induced proximity therapeutics.

## Supporting information

Synthetic procedures

---

**Supplementary Figure 1.**
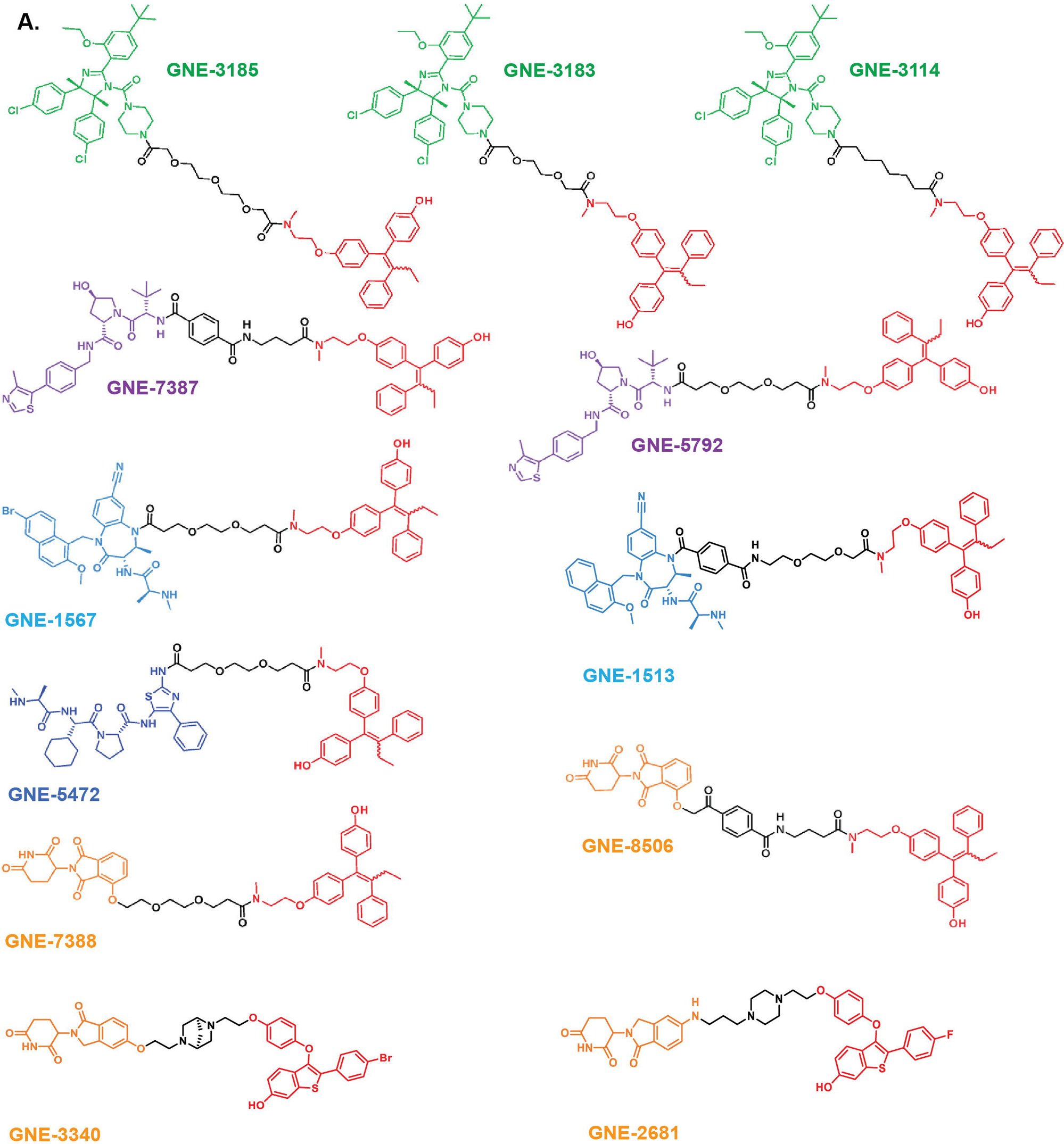
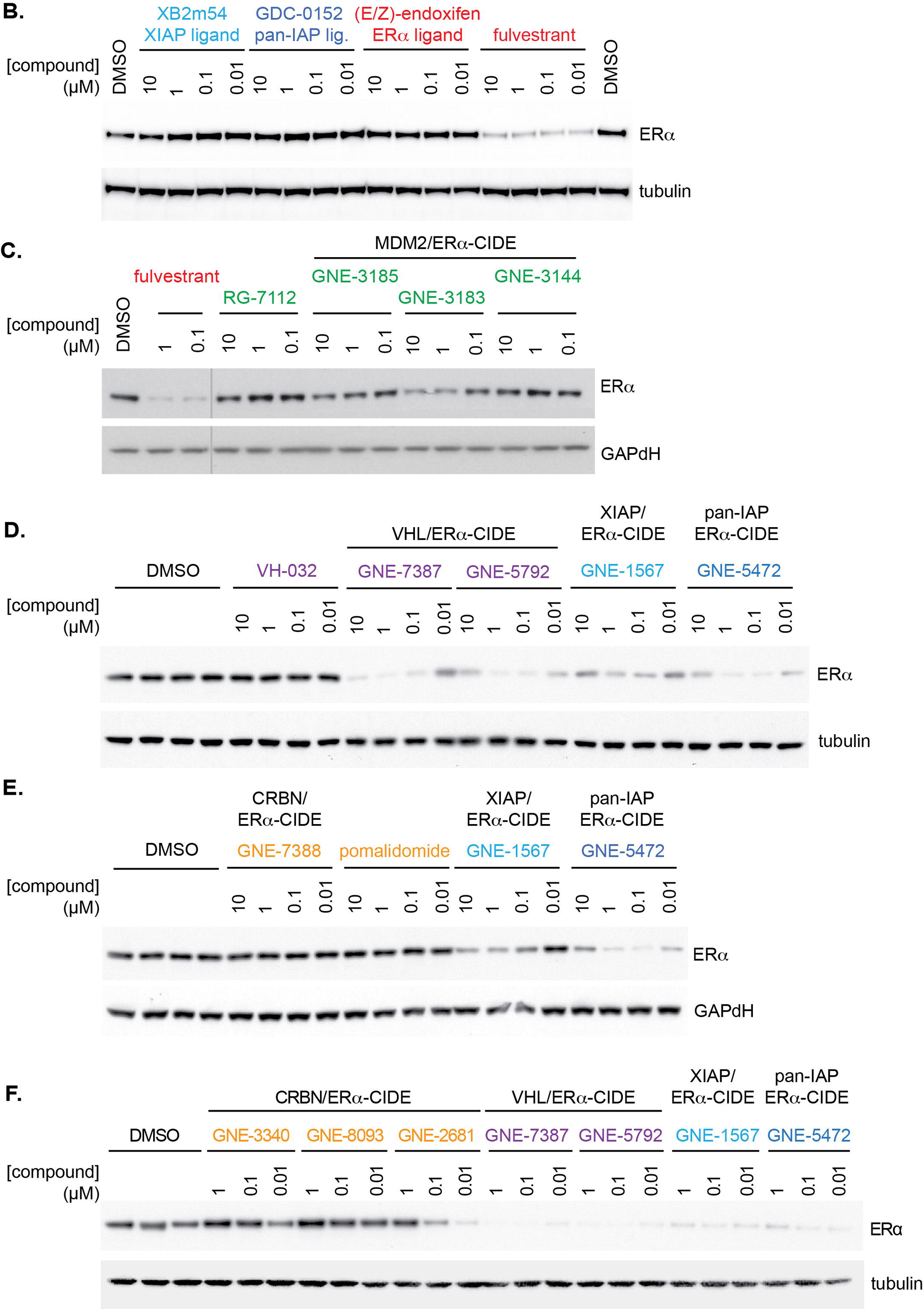
The degradation of ER**□** induced by endoxifen-based heterobifunctional degraders varies depending on the co-opted E3 ligase. S1a. Chemical structures of ERL CIDEs used in this study with the ERL ligand (E/Z-endoxifen) colored red, the linker black and the E3 ligase ligand matching the color of Fig. 1a. S1b-f. Comparison of ERL degradation mediated by various ERL-CIDEs. MCF7 cells were treated for 4 hours with the indicated concentrations of ERL-CIDEs. Cell lysates were analyzed by western blot with the indicated antibodies.

**Supplementary Figure 2.**
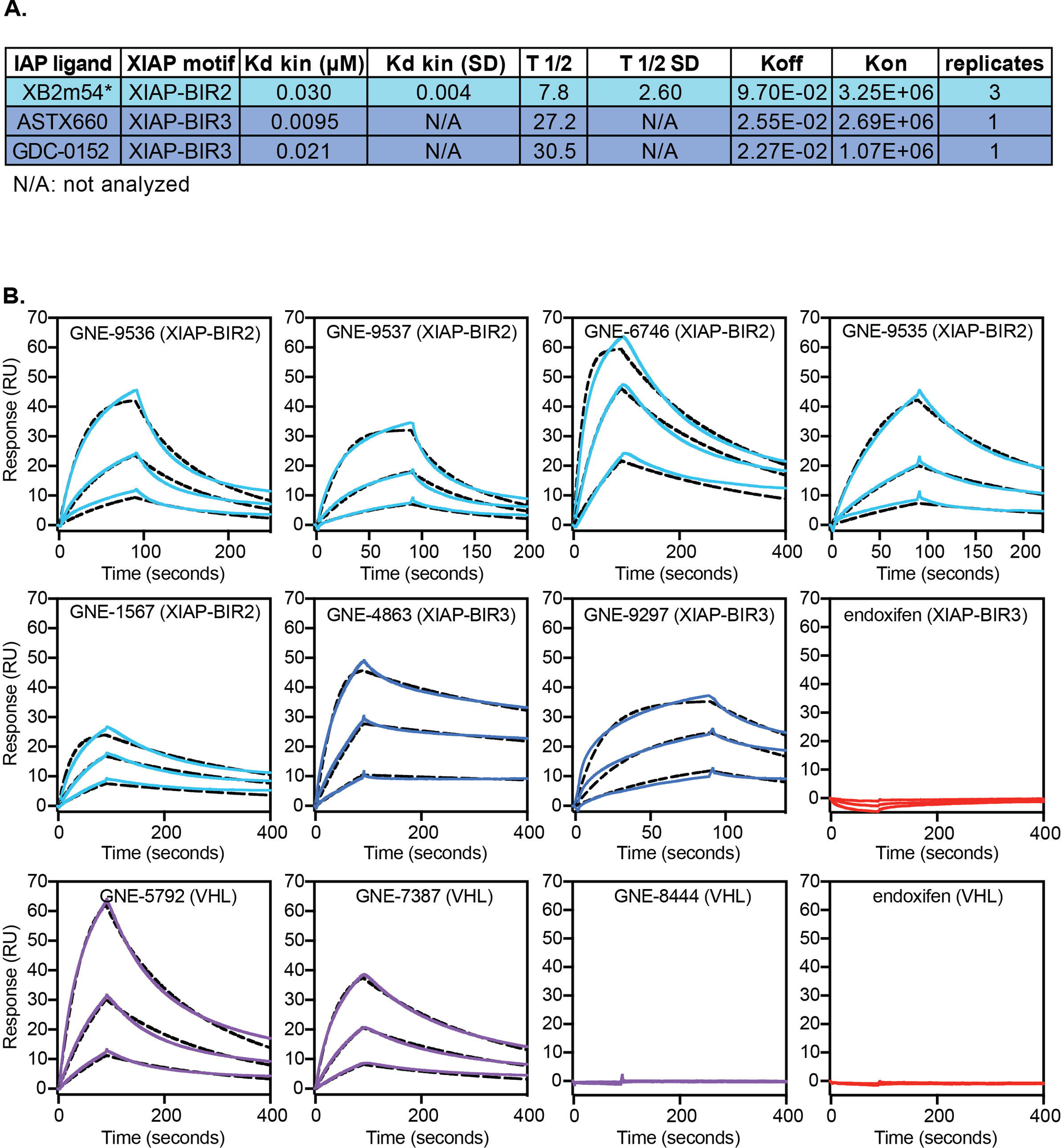
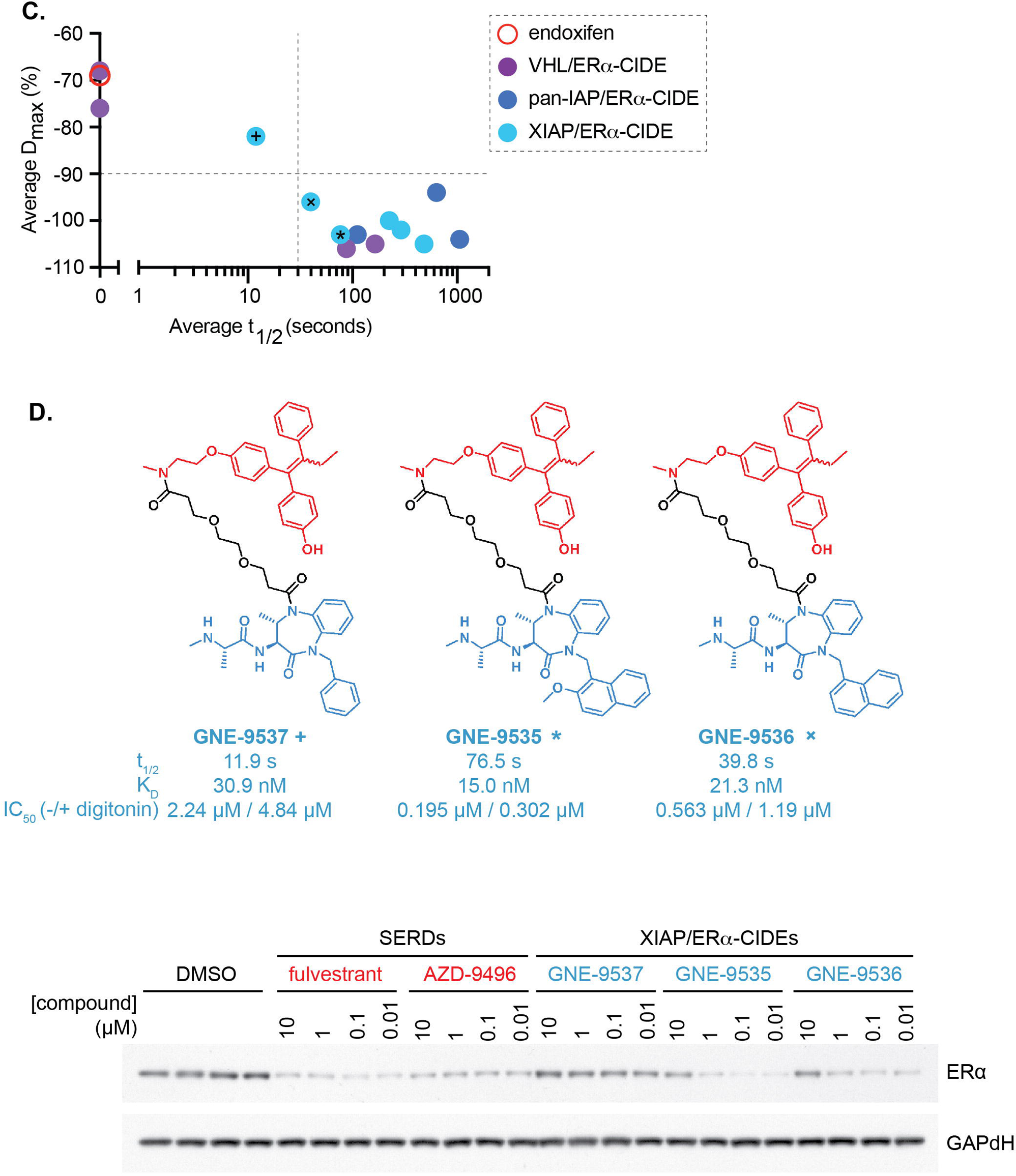
IAP-based ER**α** degraders promote high-affinity ternary complexes. S2a. SPR binding affinity and kinetic parameters for IAP ligands binding to XIAP-BIR2 or XIAP-BIR3. S2b. SPR ternary complex assessments for ERL:CIDE binding to XIAP BIR2 (light blue), XIAP BIR3 (dark blue), VHL (purple), or negative control ligand endoxifen (red). Kinetic 1:1 fits shown in black dashed lines. One of three replicates for each condition is shown. S2c. Correlation of average ternary complex half-life (t_1/2_) measured by SPR with average maximum cellular degradation (D_max_) measured by ERL immunofluorescence for a set of ERL CIDEs paired to VHL (purple), XIAP BIR3 (dark blue), or XIAP BIR2 (light blue) is shown. Compounds marked with +, x. * belong to a series that contain the same ERL ligand (endoxifen) and linker and differ only in their XIAP BIR2 ligand. The R-squared t_1/2_ / D_max_ value for the 3 compound series is 0.930 and the R-squared t_1/2_ / cellular degradation potency (DC_50_) value for the 3 compounds is 0.999. S2d. Comparison of ERL degradation mediated by XIAP/ERα-CIDE that have different XIAP binding affinity. Chemical structures of three XIAP/ERα-CIDE that contain the same ERL ligand (endoxifen, red) and linker (black) and differ only in their XIAP BIR2 ligand (light blue). Ternary complex SPR half-life (t_1/2_) and equilibrium binding constants (K_D_) are shown, as well as XIAP BIR2 potency measured by cellular nanoBRET in live versus permeabilized cells using digitonin. MCF7 cells were treated for 4 hrs with the indicated concentrations of each compound. Cell lysates were analyzed by western blot with the indicated antibodies.

**Supplementary Figure 3.**
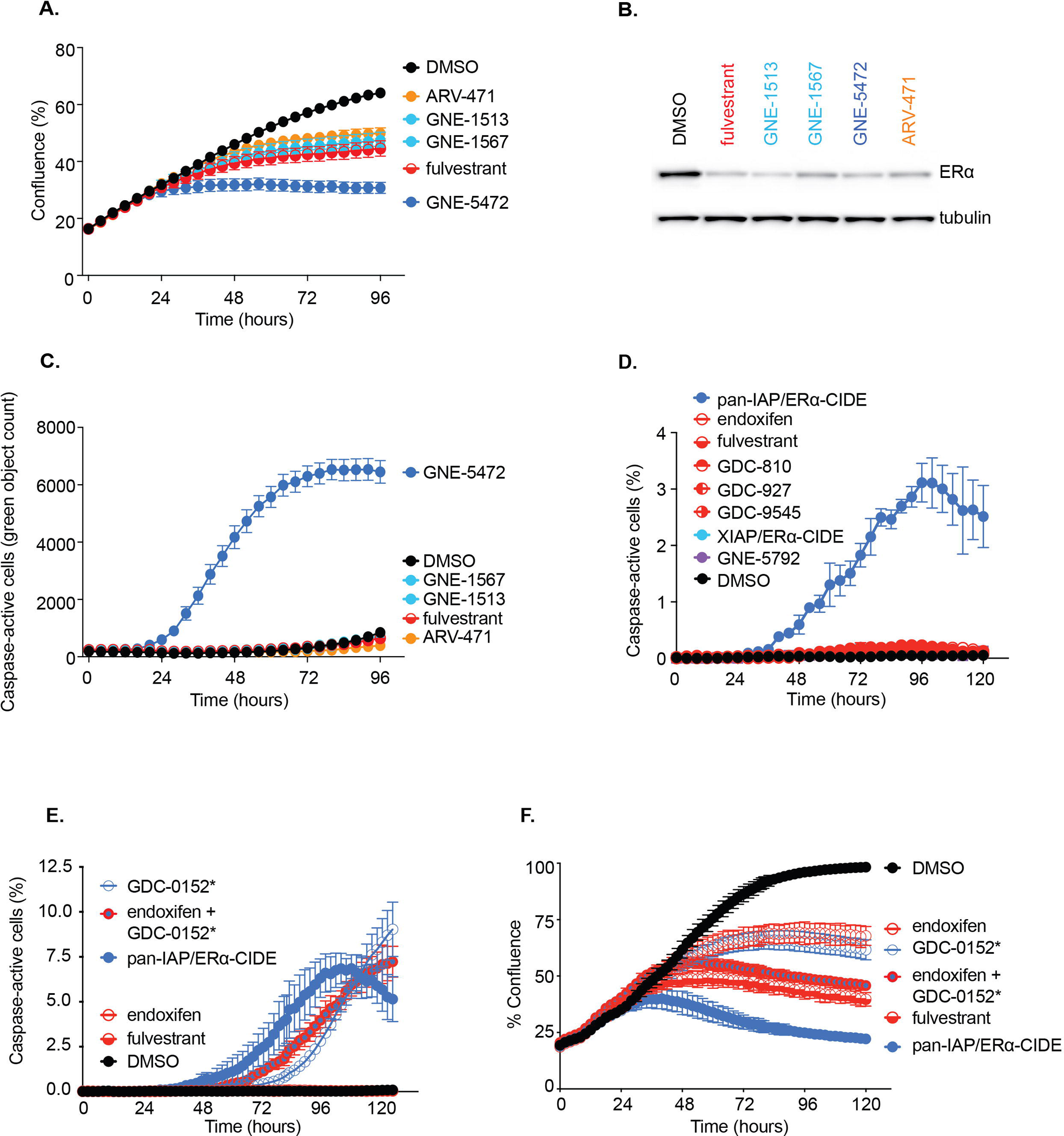
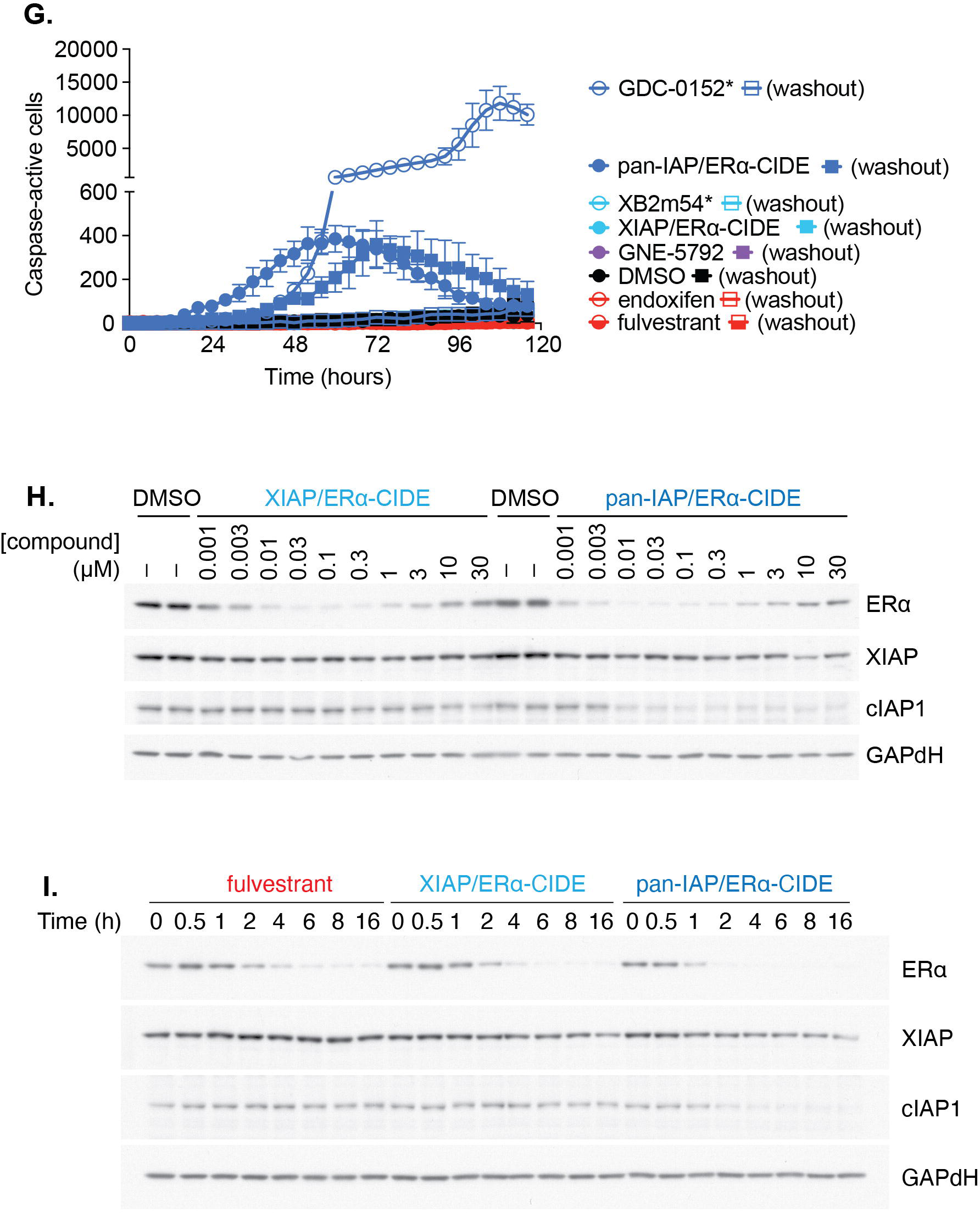
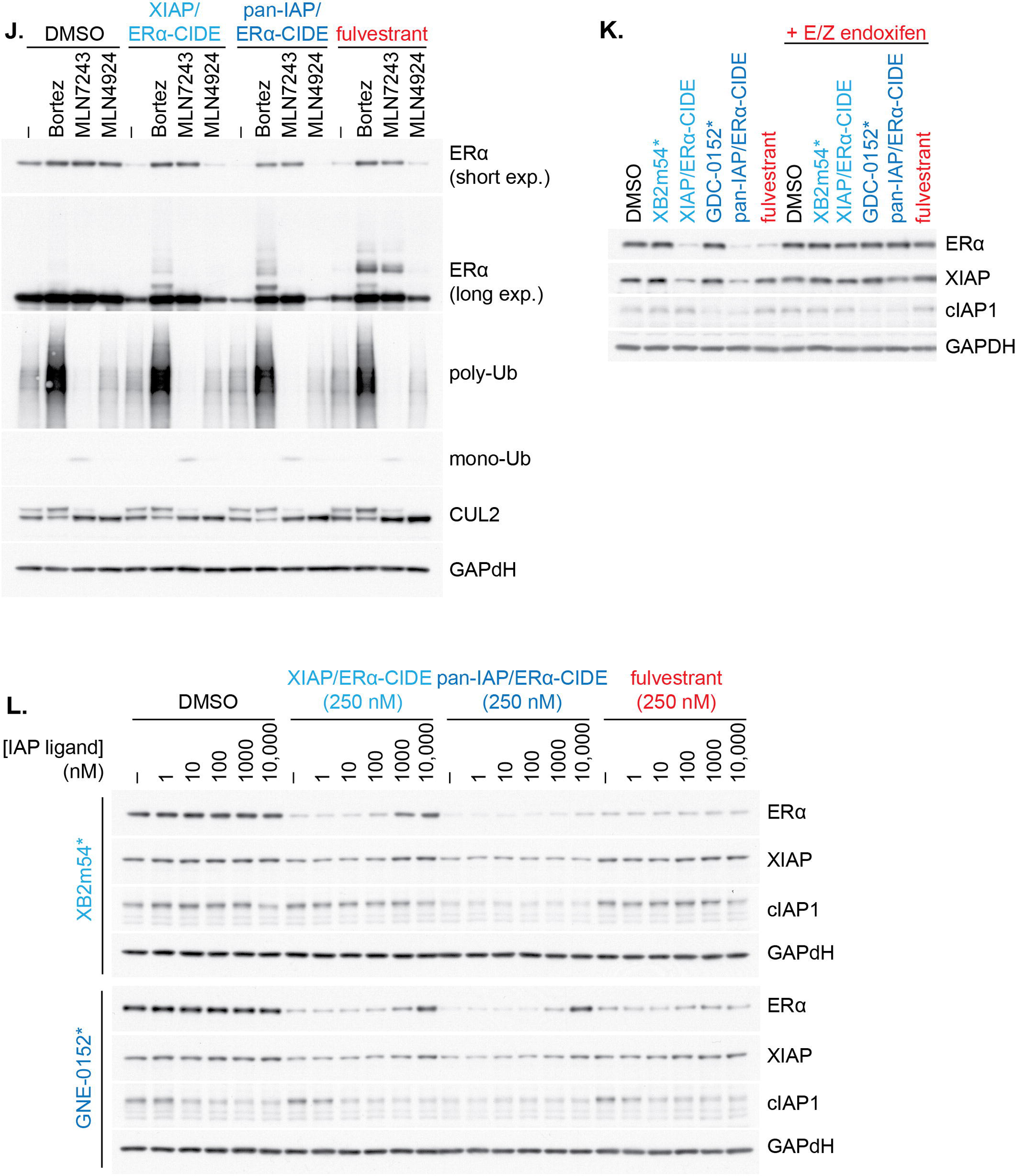
Pan-IAP antagonist-based ER**α**-CIDEs uniquely promote tumor cell death. S3a. pan-IAP/ERα-CIDE is more efficacious than the CRBN-based ERa degrader ARV-471. Side by side comparison of all three GNE compounds, fulvestrant, and ARV-471 in an Incucyte assay demonstrates that pan-IAP/ERα-CIDE is the best at inhibiting cell proliferation and is the only compound to induce caspase activation. Western blot analysis demonstrates that all compounds decrease ERa levels to approximately the same level after 4 hours. S3b. MCF7 cells were treated as indicated with 250 nM of each compound and Caspase 3/7 activation was measured over a period of 120 hours using a Caspase 3/7 activated fluorescent dye in an Incucyte instrument. S3c. MCF7 cells were treated as indicated with 250 nM of each compound and confluence as well as Caspase 3/7 activation was measured using an Incycyte instrument. S3d. MCF7 cells were treated as indicated with 250 nM of each compound. Cells were treated either continuously (solid symbols) or for 24 hrs followed by 3 media changes (open symbols) and grown in regular media. Caspase 3/7 activation was measured as in (b). S3e. MCF7 cells were treated for 4 hours with the indicated concentrations of compounds and cell lysates were analyzed by western blot with the indicated antibodies. S3f. MCF7 cells were treated with 250 nM of compound for the indicated times and cell lysates were analyzed by western blot with the indicated antibodies. S3g. MCF7 cells were pre-treated for 30 minutes with 1 uM of Bortezomib, MLN7243 or MLN4924 followed by 4 hours of treatment with DMSO or the indicated ERa degraders and cell lysates were analyzed by western blot with the indicated antibodies. S3h. MCF7 cells were treated for 4 hrs with 250 nM of the indicated ERa degraders either in the presence or absence of 10 uM endoxifen and cell lysates were analyzed by western blot with the indicated antibodies. S3i. MCF7 cells were treated for 4 hours with 100 nM of the indicated ERa degrader compounds in the presence of increasing concentrations of an IAP competitor ligand and cell lysates were analyzed by western blot with the indicated antibodies.

**Supplementary Figure 4.**
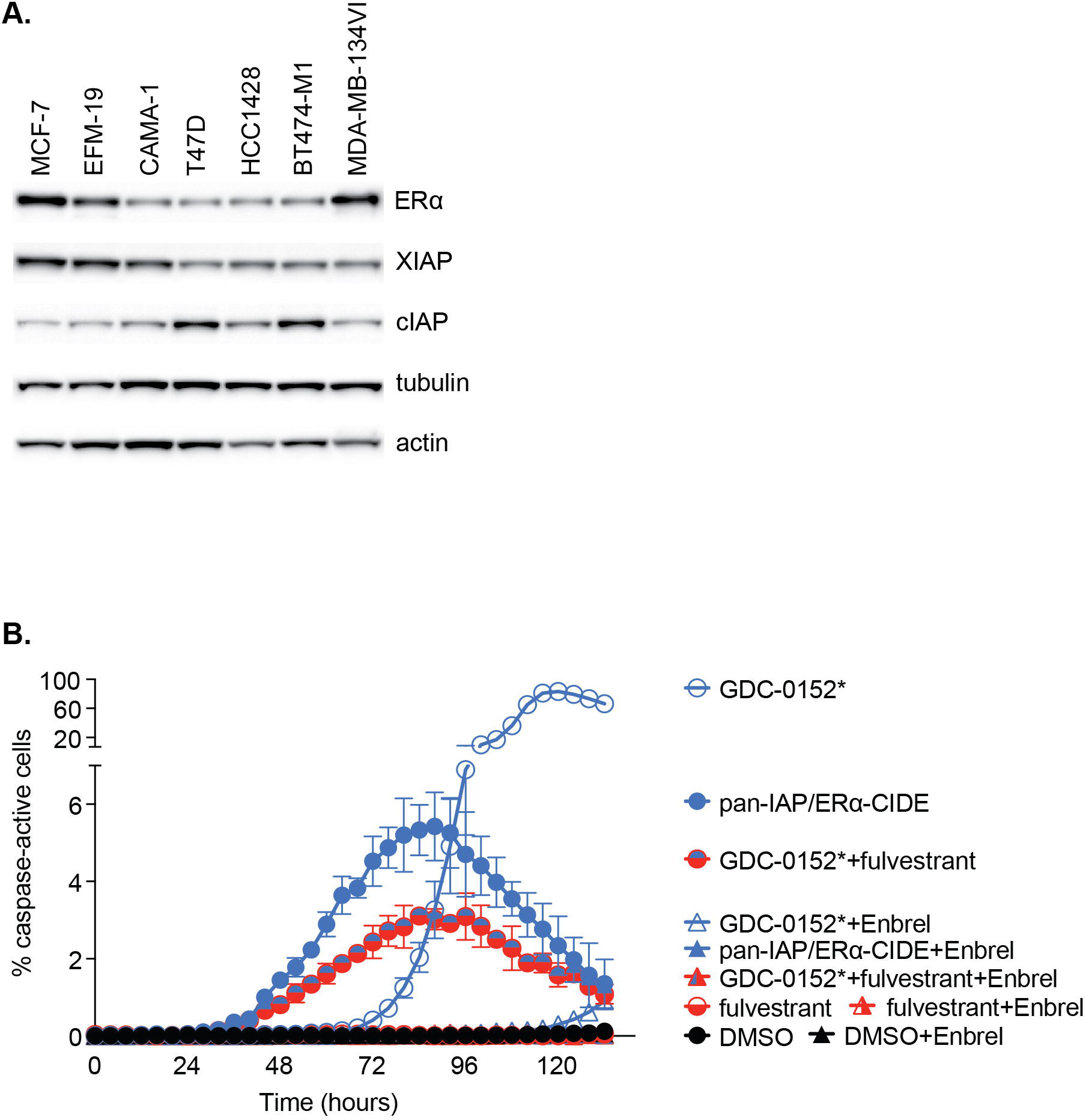
The proliferation effect induced by pan-IAP/ER**α**-CIDE is TNF**α**-driven and caspase-dependent and not attributable to a lack of XIAP or cIAP protein expression levels. S4a. A comparison of ERa, XIAP, and cIAP levels across the different ER positive breast cancer cell lines from most to least sensitive to pan-IAP/ERα-CIDE. Equal amounts of protein from the indicated cell lines were western blotted for ERa, XIAP, or cIAP and actin or tubulin for a loading control. S4b. MCF7 cells were treated with 250 nM of the indicated ERa degraders either in the absence or presence of 10 ug/mL Enbrel. Caspase 3/7 activation was measured over a period of 120 hours using an Incucyte instrument.

**Supplementary Figure 5.**
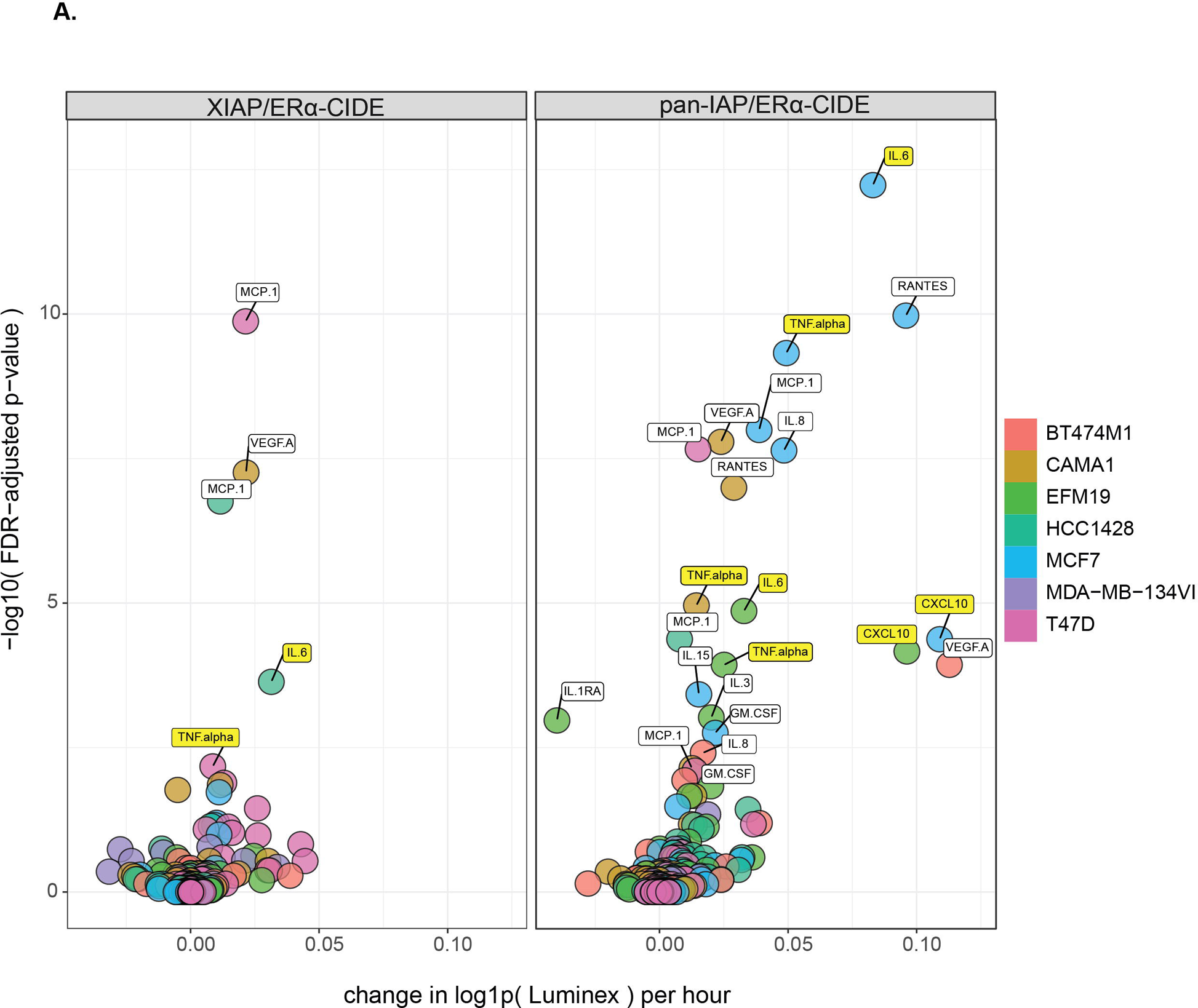
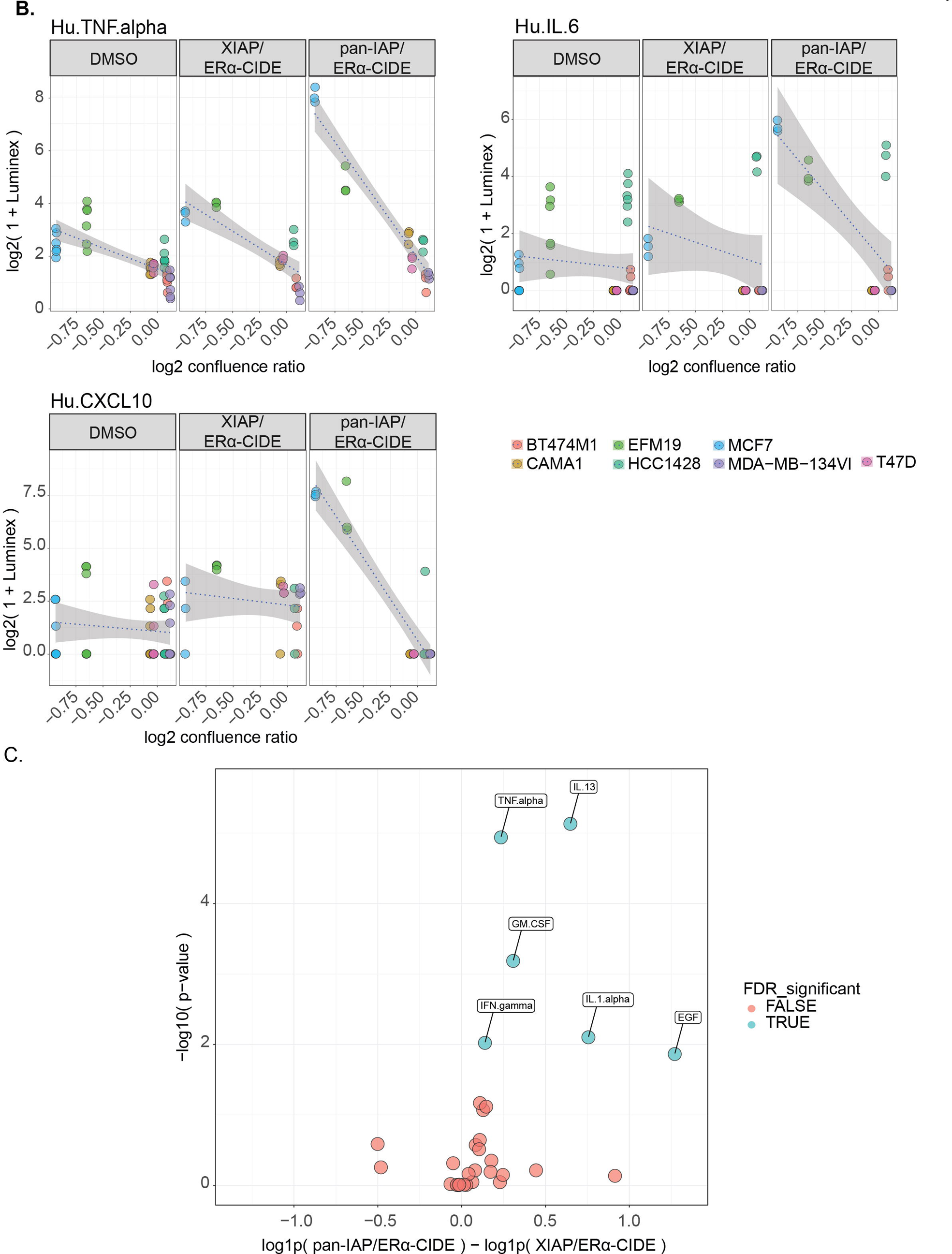
Quantification of cytokines and chemokines produced by cells in response to pan-IAP/ER**α**-CIDE treatment identified increased TNF-alpha, IL6, and CXCL10 levels as correlative to sensitivity. S5a. Volcano plots showing per-hour log fold change in Luminex-quantified cytokine scores measured at 24, 48, and 72 hours following XIAP/ERα-CIDE or pan-IAP/ERα-CIDE treatment of seven different cell lines. Notably, pan-IAP/ERα-CIDE (right panel) shows a number of cytokines with marked changes in MCF7 that are absent under XIAP/ERα-CIDE treatment (left panel). S5b. Cell line specific confluence ratios (horizontal axis) and their association with Luminex cytokine scores for TNF-alpha, IL6, and CXCL10. The MCF7 and EFM19 cells had log2 confluence ratios well below zero and also showed elevated Luminex scores at 72 hours after treatment with pan-IAP/ERα-CIDE at 72 hours compared to treatment with XIAP/ERα-CIDE or DMSO. S5c. Volcano plots showing the log fold change between effects on PBMCs of pan-IAP/ERα-CIDE compared to XIAP/ERα-CIDE in a parallel line ANCOVA of Luminex-quantified cytokines with observations at 24, 48, and 72 hours. It was observed that TNF-alpha, IL13, GM-CSF, IFN-gamma, IL1-alpha, and EGF showed significantly higher responses from pan-IAP/ERα-CIDE than from XIAP/ERα-CIDE, consistently across the three time points.

**Supplementary Figure 6.**
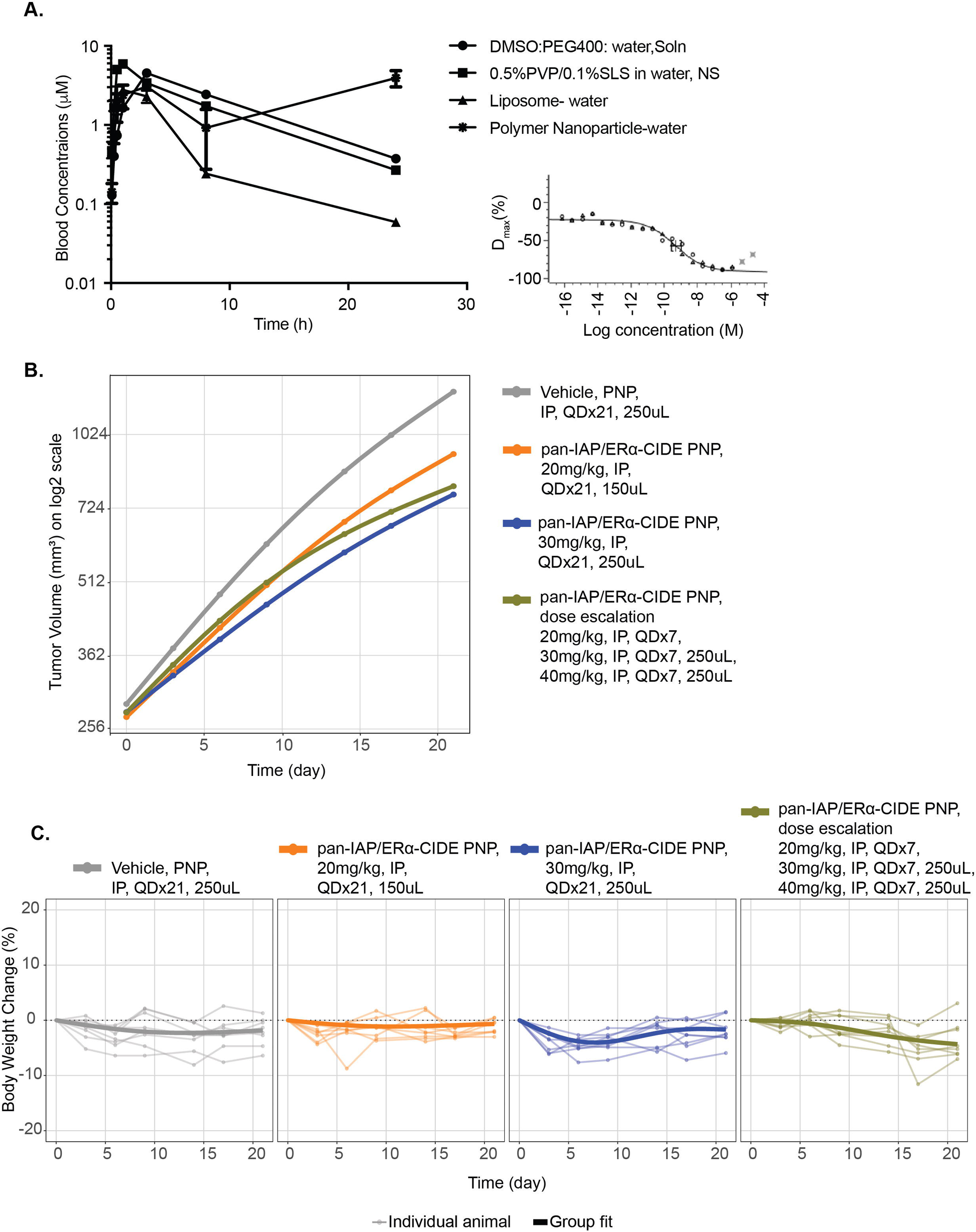
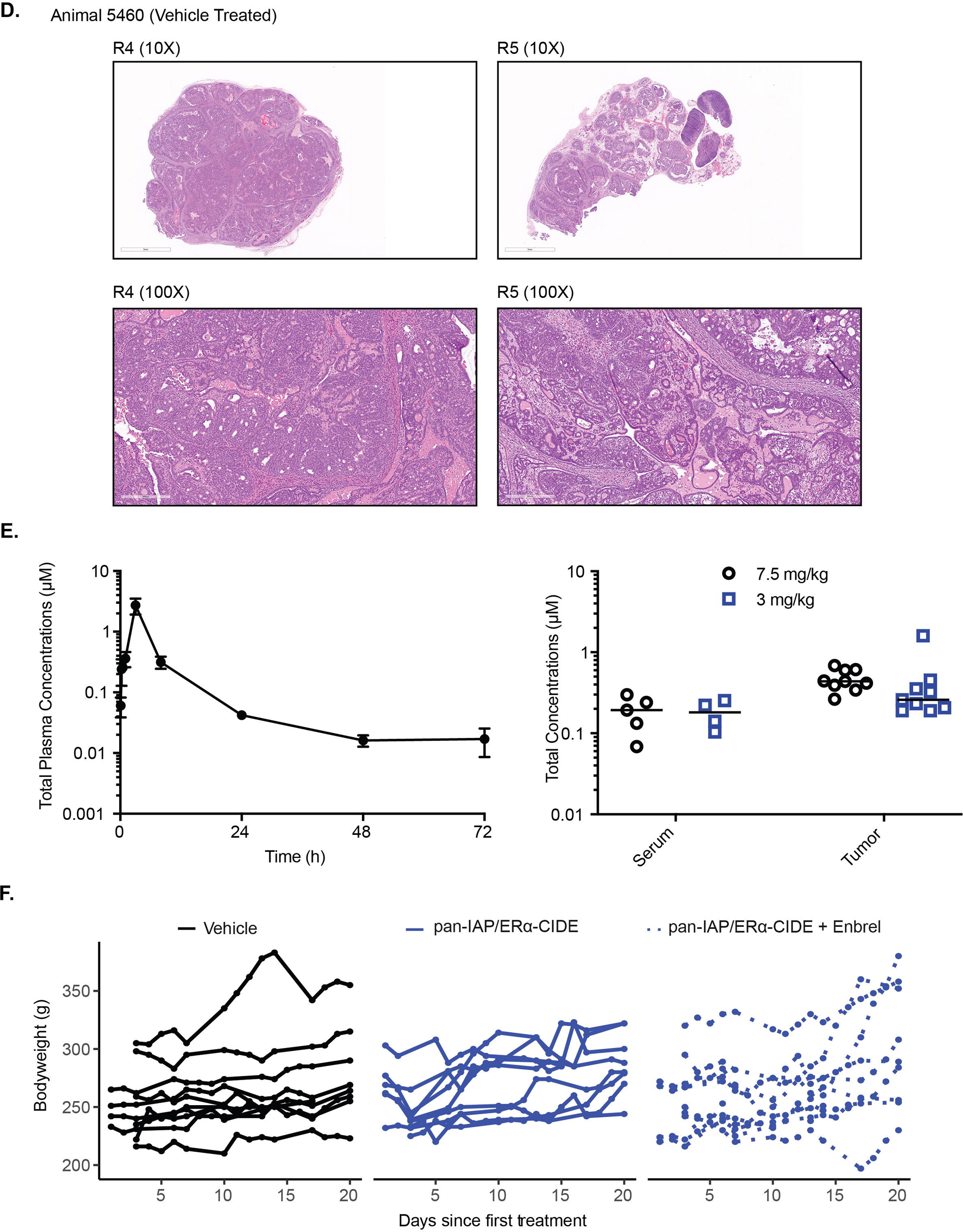
In vivo testing of pan-IAP/ER**α**-CIDE in mice and rats demonstrated the importance of an intact immune system for significant efficacy. S6a. Total blood concentration of pan-IAP/ERα-CIDE following IP administration at dose levels 5 and 20 mg/kg to female C57Bl-6 mice. n=3 mice per group were dosed with either 10% DMSO, 35% PEG400 55% water, adjusted to pH4 with NaOH; 0.5%PVP/0.1%SLS in water as nanosuspension; liposomes; and polymer nanoparticle (PNP) at a dose volume of 10mL/kg. The inset graph is from an in vitro experiment with MCF7 cells treated with PNP formulated pan-IAP/ERα-CIDE. Changes in ERL protein levels in MCF7 cells were measured after 4 hour treatment with a range of concentrations and endogenous ERL protein levels measured using high content fluorescence imaging. S6b. Modest to insignificant tumor growth inhibition was observed with PNP-formulated pan-IAP/ERα-CIDE treatment in a MCF7 xenograft model. MCF7 cells were transplanted on C57Bl-6 mice and after tumor formation, 8 mice per group were treated as indicated. The graph depicts the average tumor volume versus time after treatment. S6c. The body weights of mice from b. did not significantly change over treatment time. Individual body weights and the group fit of the body weight as percent change from the beginning of treatment were graphed over time. S6d. Histological staining of vehicle treated tumors demonstrates the variability of the NF1 deficient tumor models, where R4 represents a more homogenous tumor and R5 a more heterogeneous tumor. S6e. Total serum and tumor concentrations of pan-IAP/ERα-CIDE following IP administration at dose level 3 mg/kg and 7.5 mg/kg to female Sprague-Dalwey rats. n=9 per group were either dosed with polymer nanoparticle formulation at a dose volume of 0.1mL S6f. Treatment of NF1 deficient rats with pan-IAP/ERα-CIDE or pan-IAP/ERα-CIDE + Enbrel did not alter bodyweight over the treatment period. The body weights of individual rats were monitored and graphed with respect to time post treatment.

**Extended Data Heat Map.**
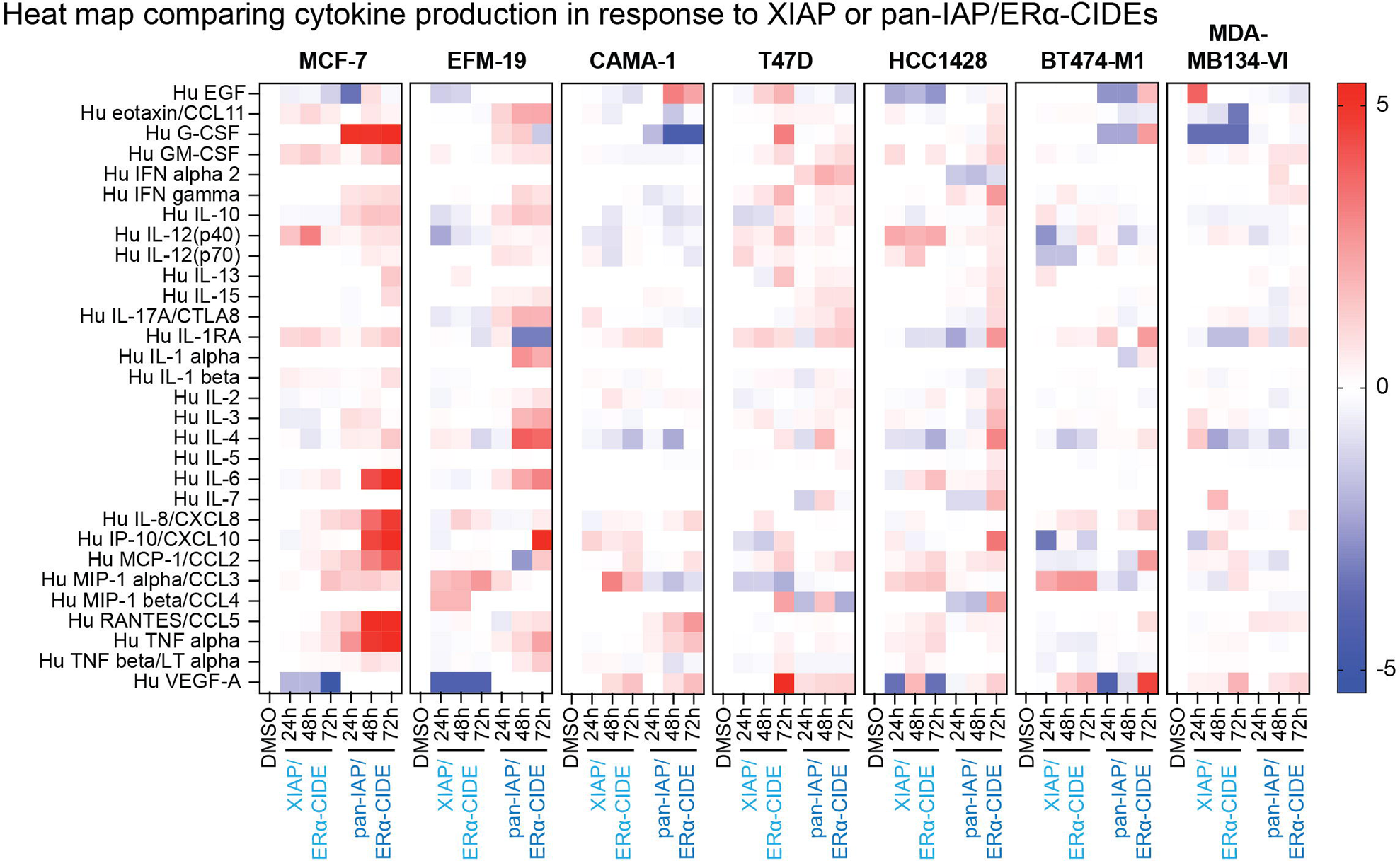
Heat map comparing cytokine production in response to either XIAP or pan-IAP/ERα-CIDEs. The indicated cell lines were treated with DMSO or 250 nM of either XIAP/ERα-CIDE or pan-IAP/ERα-CIDE for 24, 48, or 72 hours. Following treatment, the media from the treated cells was sent for analysis by Luminex to measure the amount of cytokines present. Data is graphed as a log2 fold change of cytokine detected from treatment compared to DMSO.

## Methods

### Chemical reagents

Fulvestrant (CAS# 129453-61-8), endoxifen (CAS# 112093-28-4), brilanestrant (GDC-0810, CAS# 1365888-06-7), GDC-0927 (CAS# 1642297-01-5) and giredestrant (GDC-9545, CAS# 1953133-47-5) were acquired from commercial sources. XB2m54 analog was made according to the reported method: Kester, R. F. et al J. Med. Chem. 56, 7788-7803 (2013). Synthesis of degrader GNE-1513 was made according to the method reported in patent WO2017201449A1. See supplementary information for synthetic procedure of included compounds.

### Western blot analysis

Cells were lysed in 6 M urea buffer (20 mM Tris pH 7.5, 6 M Urea, 0.1% Triton X-100, 12.5 mM NaCl and 2.5 mM MgCl2) supplemented with Roche protease inhibitor cocktail, centrifuged at 14000 rpm and supernatant collected. Cell lysates were estimated for total protein content using Pierce™ BCA Protein Assay Kit (23225). Cell lysates were prepared in LDS sample buffer and sample reducing agent (Invitrogen). Equal amounts of proteins were resolved on SDS-PAGE gels (NuPAGE 4-12% bis-tris) and transferred to nitrocellulose membranes. Following blocking (5% non-fat dry milk in TBS with 0.1% tween 20), membranes were incubated with the indicated antibody in blocking buffer (1% non-fat dry milk in TBS with 0.1% tween 20) overnight at 4oC. Membranes were washed four times with TBST buffer and incubated with the appropriate secondary antibody-HRP (1:10,000) in blocking buffer for 1hr at RT. Following membrane wash four times with TBST, chemiluminescence was developed.

ERa (sc-8002, Santa Cruz), cIAP1 (7065S, Cell Signaling Technology), XIAP (14334S, Cell Signaling Technology), MDM2 (OP46, Calbiochem), CRBN (11435-1-AP, Proteintech), VHL (68547S, Cell Signaling Technology), MEK1 (8727S, Cell Signaling Technology), SP1 (5931S, sc-59), tubulin (5346S, Cell Signaling Technology), Gapdh (3683S, Cell Signaling Technology), Cul2 (511800 Invitrogen), Ub (sc-8017, Santa Cruz), p100/p52 (4882S, Cell Signaling Technology), actin (12620S, Cell Signaling Technology)

### Global cellular expression analysis

RNA was isolated from cells using the RNeasy mini kit (Qiagen). cDNA was synthesized from 1 ug of RNA using SuperScript® III First-Strand Synthesis SuperMix. Gene expression was measured from cDNA using TaqMan™ Universal PCR Master Mix with TaqMan probes for TNFa (Hs00174128_m1) and GAPDH (Hs02758991_g1). The delta delta Ct method was used to calculate relative gene expression.

### Gene silencing studies

MCF-7 cells were transfected with Silencer Select siRNA (Thermo Fisher Scientific) against XIAP (s1454) or a non-targeting control (4390843, Negative Control No. 1) using RNAiMAX (Thermo Fisher Scientific) according to the manufacturer’s protocol.

### Subcellular fractionation

Nuclear and cytoplasmic fractions of MCF-7 were isolated using the NE-PER Nuclear and Cytoplasmic Extraction Kit (Thermo Fisher Scientific) following manufacturer’s instructions. Protein concentration was determined using the BCA Protein Assay Kit (Pierce) and equal amounts of protein were analyzed by western blot.

### Cell growth studies

Equal number of cells were plated in 50 ul of media in black 96 well clear bottom plates and allowed to adhere overnight. 50 ul of media containing 2x the amount of compound treatment was added to the wells. Cell growth was monitored using the Incucyte instrument (Sartorius) by analyzing confluence over time. Apoptosis was measured by using the green fluorescent caspase 3/7 dye (Sartorius) and determining the number of green fluorescent objects per well over time. 10 ng/ml TNFa

### Binary SPR method

SPR measuring binding of small molecule ligands to XIAP Bir2 or Bir3 proteins was performed as previously described (*J Am Chem Soc*. 2021 Jul 21;143(28):10571-10575). **E**xperiments were performed on a Biacore T200 or S200 instrument. XIAP BIR2 or Bir3 was biotinylated through an N-terminal Avi-tag and immobilized on a neutravidin (Thermo, 31000) coated C1 sensor chip (Biacore). The running buffer contained 50mM HEPES, pH 7.2, 150 mM NaCl, 0.001% Tween-20, 5 mM DTT, 0.2% PEG 3350, and 2% DMSO. Compounds were tested in 6-point dose response and concentrations were adjusted between 5 nM to 50 μM depending on potency. The contact time varied between 30 and 60 sec and the dissociation time between 30 and 600 sec. Data was fit to a Kinetic 1:1 Binding Model in Biacore Evaluation software.

### Ternary Complex SPR method

Ternary complex SPR experiments were performed as previously described (*ChemMedChem*. 2020;15(1):17-25). A Series S C1 chip coupled to neutravidin (500-2000 RU) was used in a Biacore 8k (GE Health Sciences). The running buffer was 50 mM HEPES pH 7.5, 150 mM NaCl, 0.2% (w/v) PEG-3350, 0.5 mM TCEP, 0.01% Tween 20, 5 nM biotin, and 2% (v/v) DMSO. ERα (305-R548) with an N-terminal His-avi tag was immobilized on the sensorchip at a level of approximately 300-1000 RU. All channels were blocked by injecting 100 μg/mL amine-PEG-biotin (Thermo Fisher). Biotin was also used in the buffer to ensure no non-specific binding of the ligase proteins to the sensorchip through the biotinylated-tags. CIDEs were injected at 1 µM to saturation followed by ligase in 3-fold dose response at a top concentration of 100 nM for XIAP BIR2 (52-236) (C202A, C213G) and VHL (55-213) in complex with Elongin B (1-118) and Elongin C (17-112), and 1 µM for XIAP BIR3 (241-356). CIDE was injected at 1 µM in between every ligase injection to ensure ERα was saturated with CIDE on the chip surface. Data was analyzed with a 1:1 affinity model in Biacore 8k Evaluation Software and KD, k_on_, k_off_ and half-life (t_1/2_) = ln(2)/ k_off_ is reported. In cases where ternary complexes were not observed, KDs are reported as > top ligase concentration tested with half-life (t_1/2_) = 0. Data was collected in triplicate. Figures were made in Graphpad Prism 9.

### Breast cancer cell ER**α** high content fluorescence imaging assay

MCF-7 breast cancer cells were seeded on day 1 at a density of 10,000 cells per well in 384 well poly-lysine coated tissue culture plate (Greiner # T-3101-4), in 50 µL/well RPMI (phenol red free), 10% FBS (Charcoal stripped), containing L-glutamine. On day 2, compounds were serially diluted in DMSO spanning the concentrations 100 µM to 0.2 nM, in a Labcyte Echo Qualified 384-well polypropylene plate (P-05525). DMSO and 5 µM Fulvestrant (control compound) were added to designated wells. Compounds and controls were dispensed into the cell culture plate, using a Labcyte Echo acoustic dispenser (the final dispensed volume of each control and each compound was 50 nL and the final DMSO concentration was 0.1% v/v). Cell plates were incubated at 37 °C, for 4 hours. Fixation and permeabilization were carried out using a Biotek EL406 plate washer and dispenser as follow. Cells were fixed by addition of 15 µL of 16% paraformaldehyde (Electron Microscopy Sciences #15710-S) directly to the 50 µL cell culture medium in each well using the peristaltic pump 5 µL cassette on a Biotek EL406 (final concentration of formaldehyde was 3.7% w/v). Samples were incubated 30 minutes. Well contents were aspirated and 50 µL/well of Phosphate Buffered Saline (PBS) containing 0.5% w/v bovine serum albumen, and 0.5% v/v Triton X-100 (Antibody Dilution Buffer) was added to each well. Samples were incubated for 30 minutes. Well contents were aspirated and washed 3 times with 100 µL/well of PBS. Immunofluorescence staining of estrogen receptor alpha (ESR1) was carried out using a Biotek EL406 plate washer and dispenser as follows. The well supernatant was aspirated from the wells and 25 µL/well of anti-ESR1 mAb (F10) (Santa Cruz sc-8002) diluted 1:1000 in Antibody Dilution Buffer was dispensed. Samples were incubated for 2 hours at room temperature. Samples were washed 4 times with 100 µL/well of PBS. 25 µL/well of secondary antibody solution (Alexafluor 488 conjugate anti-mouse IgG (LifeTechnologies #A21202) diluted 1:1000 and Hoechst 33342 1 µg/mL diluted in Antibody Dilution Buffer) were dispensed into each well. Samples were incubated for 2 hours at room temperature. Samples were washed 3 times with 100 µL/well of PBS using a Biotek EL406. Quantitative fluorescence imaging of ESR1 was carried out using a Cellomics Arrayscan V (Thermo). Fluorescence images of the samples (Channel 1: XF53 Hoechst (DNA stain); Channel 2: XF53 FITC (ESR1 stain)) were acquired using a Cellomics VTI Arrayscan using the Bioapplication “Compartmental Analysis” using the auto-exposure (based on DMSO control wells) setting “peak target percentile” set to 25% target saturation for both channels. Channel 1 (DNA stain) was used to define the nuclear region (Circ). Measurements of “Mean_CircAvgIntCh2”, which is the Alexafluor 488 fluorescence intensity (ESR1) within the nuclear region, was measured on a per cell basis and averaged over all the measured cells. Data analysis was carried out using Genedata Screener Software, with DMSO and 5 nM Fulvestrant treated samples being used to define the 0% and −100% changes in ESR1. The “Robust Fit” method was used to define the inflexion point of curve (DC_50_) and the plateau of the maximal effect (S_inf_).

### PBMC co-culture

Isolated human PBMCs (ATCC PCS-800-011) were activated with human monoclonal antibodies against human CD3 (OKT3, Life Technologies, 16-0037) coated onto a flask and 2 ug/ml CD28 (CD28.2, Life Technologies, 16-0289) for 3 days. Stimulated PBMCs were maintained in RPMI-1640 supplemented media with 100 ng/ml CD3 and 10 ng/ml recombinant human IL-2 (Biolegend, 589104).

### Luminex analysis

Cell culture supernatants were diluted in growth media and analyzed for cytokines using the MILLIPLEX MAP Human Cytokine/Chemokine Magnetic Bead Panel (Premixed 30 Plex) according to manufacturer’s instructions.

### Immunohistochemistry

The staining intensity of the population of tumor cells with epithelial morphology was evaluated qualitatively using the H-score method [74].The staining intensity of the population of tumor cells with epithelial morphology was evaluated qualitatively using the H-score method [59]. The staining intensity of individual cells was judged on a scale of 0 (staining absent), 1 (weak), 2 (moderate) or 3 (strong), and the percent (0 to 100) of cells with each intensity was recorded. The overall H-score (0-300) is the sum of the four products of the staining intensity and the percent of cells at each intensity.

Four micron thick formalin-fixed, paraffin-embedded (FFPE) tissue sections were deparaffinized, rehydrated and processed on a Ventana Discovery XT autostainer using CC1 standard antigen retrieval. Anti-estrogen receptor clone SP1 was used as provided (“ready to use”; RTU) by the manufacturer (1 ug/mL) with primary antibody incubation at 37 C for 32 minutes, followed by anti-rabbit OmniMap HRP for 16 minutes with DAB chromogen detection.

### Proteomic sample preparation

MCF-7 cells were harvested with 8 M urea, collected and centrifuged to clear away cellular debris and recover protein in lysate. Protein concentration was calculated using a Bradford Assay. Lysates were then reduced with 4.5 mM dithiothretiol (DTT) at 65°C for 30 mins followed by 20 min alkylation of 11 mM iodoacetamide (IAA) at room temperature in the dark. Urea was diluted to working 2 M concentration with 20 mM HEPES (pH 8.0) and tryptic (Promega) digestion performed at an enzyme to substrate ratio of 1:25 overnight with incubation at 37°C. Tryptic peptides were acidified with trifluoroacetic acid before desalting using a C-18 Sep-pak (Waters) and dried. Samples were resuspended with 100 µL of 100 mM HEPES pH 8.0 followed by full vial labeling of TMT10plex reagent + TMT-131C (Thermo) for 1.5 hrs per the manufacturer’s protocol. TMT labeling was quenched with the addition of 8 µL of 5% hydroxylamine, combined and desalted with C-18 Sep-pak (Waters). Approximately 400 ug of eluents were subsequently dried. High pH reversed phase liquid chromatography was used to fractionate the 400 ug of TMT labeled peptides using an Agilent 1200 series HPLC system. Peptides were resuspended in 0.1% TFA/water and injected onto a Zorbax 300 Extend C-18 analytical column (2.1 x 150 mm and 3.5 um particle size). A linear gradient was applied of 15-60% solvent B (50 mM ABMIC in 90% MeCN) with Solvent A consisting of 50 mM AMBIC in 5% MeCN. Fractions were collected in 45 second intervals for a total of 63 mins across the gradient. A total of 96 fractions were obtained and subsequently every 12th fraction was combined for a final set of 24 distinct groups. Samples were then lyophilized, desalted via a STAGEtip, and injected for LC-MS/MS analysis.

### Mass spectrometric analysis of global proteins

Data was collected on Orbitrap Fusion Tribrid Mass Spectrometers (Thermo Fisher Scientific). Peptides were injected onto a New Objective PicoFrit Acquity® BEH130Å C18 column (1.7 µM, 100 µM × 250 mm) in Solvent A (98% water, 2%acetonitrile, 0.1% formic acid) via a Nanospray Flex Ion-Source (Thermo Scientific) at a voltage of 1.9 kV with a flow rate of 0.7 µL/min in Solvent A (98% water/2%acetonitrile/0.1% formic acid), and separation at a flow rate of 0.5 µL/min with a linear gradient of 2 to 35% solvent B (98% acetonitrile/2% water/0.1% formic acid) over 158 mins. Full MS scans were collected at 120,000 resolution in the orbitrap from 350 to 1600 m/z, with an automatic gain control (AGC) target of 2 × 10e5, and a maximum injection time of 50 ms. MS2 ions were selected with a 0.7 Da isolation width, AGC of 5 × 10e3, and a maximum injection time of 100 ms using a top speed data dependent mode. Fragmentation with CID energy of 35 and analyzed in ion trap. MS3 spectra were detected in the orbitrap by isolating 8 MS2 ions using the synchronized precursor selec-tion (SPS) mode and fragmentation via higher collision dissociation energy (HCD) of 55. The associated AGC target of 2 × 10e5, a maximum injection time of 150 ms, isolation width of 1.4 Da, and a resolution of 50,000 at 200 m/z made up the remaining parameters.

### Mass spectrometric data identification and quantification

MS/MS spectra were then searched using Mascot (v.2.4.1) with a UniProt DB (2017_08), appended with commonly found contaminating proteins as well as concatenated decoy sequences. The search parameters used allowed for 2 missed trypsin cleavages, precursor ion tolerance of 50 ppm, and a fragment ion tolerance of 0.8 Da. Searches also allowed for variable modifications of methionine oxidation (+15.9949 Da), and static modifications of cysteine (+57.0215) for carbamidomethylation and TMT tags on lysine and peptide N termini (+229.1629). Spectral peptide mapping (PSMs) were filtered with a 2% false discovery rate (FDR) on the peptide level and subsequently at 2% on the protein level. TMT reporter ions were quantified with an in-house software package known as Mojave46 by calculating the highest peak within 20 ppm of the theoretical reporter mass window whilst correcting for any isotope purities. Further filtering of quantified PSMs using total TMT reporter ion intensities that were greater than 50,000 and an isolation specificity of greater than 0.7 was performed. Then summarization to their matched proteins finalized the list of quantifiable PSMs. Protein abundance ratios among different treatment groups to DMSO were computed (as the percentage of sum total TMT signal for the protein across all treatment groups over DMSO), using R (3.2.2). P-values were also calculated based on student t-test with two-tailed comparison assuming equal variance.

### Pharmacokinetic studies

Plasma samples were analyzed for GNE-5472 at Genentech. LCMS/MS methios was as follows: Shimadzu Nexera (Columbia, MD, USA) coupled to a QTRAP® 5500 mass spectrometer (AB Sciex, Foster City, CA, USA) equipped with a turbo-electrospray interface in positive ionization mode. The aqueous mobile phase was water with 0.1% formic acid (A) and the organic mobile phase was acetonitrile with 0.1% formic acid (B). The gradient was started with 15% B, and then increased to 90% B in 0.3 minutes, and maintained at 90 % B for another 0.3 minutes, decreased to 15% B within 0.01Lminutes, and maintained at 15% B for another 0.2 minutes. The flow rate was 1.2 mL/min and the cycle time (injection to injection including instrument delays) was approximately 0.8Lminutes. A volume of 0.5 µL of the final extract was injected onto the analytical Kinetex XB-C18 100A column (30L×L2.1Lmm, 2.6 μm).

GNE-5472 quantitation was carried out using the multiple reaction monitoring (MRM) transition m/z 609.942 → 260.1, for the internal standard (Loperamide) (MRM) transition m/z 477.145 → 266.200.The optimized instrument conditions included a source temperature of 600 °C, a curtain gas pressure of 50 psi, a nebulizing gas (GS1) pressure of 50 psi, a heating gas (GS2) pressure of 50 psi, and a collision energy (CE) of 39 V for G03061334, 35 V Loperamide. IonSpray needle voltage was set at 5500 V. LC-MS/MS data were acquired and processed using Analyst software (v1.6.2). The calibration curve for quantitation of GNE-5472 was constructed by plotting the GNE-5472 to internal standard, Loperamide, peak area ratios with a weighted 1/x quadratic regression.The lower limit of quantiation (LLOQ) was 0.00289 µM.

### In vivo treatment in rat breast cancer model

All animal experiments were approved and performed with in accordance with Van Andel Institute IACUC approved protocols. Tumor bearing *NF1^PS-8^* CD Sprague Dawley female rats between 6-14 weeks old were used. For treatment studies, tumor growth was evaluated twice weekly using digital calipers and tumor volume calculated by length x width x depth. When tumor volume reached approximately 500 mm^3^ rats were randomized into treatment groups and dosed for 3 weeks or until rats reached euthanasia criteria. Respective doses were empty polymeric nanoparticle (PNP) vehicle, 7.5 mg/kg G03055472 PNP, or 7.5 mg/kg G03055472 PNP + 10 mg/kg rodent-like Enbrel (mTNFRII-IgG2a). Vehicle PNP and G03055472 PNP were administered once daily via intraperitoneal (i.p.) injection. Rodent-like Enbrel was administered as a 0.5 ml bolus 3x weekly, subcutaneously (s.c.), 4 hours prior to G0355472 PNP. Body weights were collected daily during the duration of dosing and both vehicle PNP and G03055472 PNP doses were based individual body weights. Animals were euthanized when tumor size exceeded 5000 mm^3^ and tissues were harvested for analysis.

