## Supplementary material for "Engineering ERα degraders with pleiotropic ubiquitin ligase ligands maximizes therapeutic efficacy by co-opting distinct effector ligases": Table 1

|  | **Average DC50 [M]** | **Avg DC50 [M] error** | **Average Dmax (%)** | **Avg Dmax (%) error** | **ligand/degrader class** |
| --- | --- | --- | --- | --- | --- |
| endoxifen • | 3.7E-11 |  | -69 |  |  |
| endoxifen+linker.1 | 1.4E-09 |  | -68 |  |  |
| endoxifen+linker.2 | 2.3E-09 |  | -71 |  |  |
| endoxifen+linker.3 | 6.4E-09 |  | -79 |  | ERa ligands |
| fulvestrant | 4.2E-11 | 1.1E-11 | -103 | 0 |  |
| fulvestrant | 1.6E-10 | 3.5E-10 | -102 | 2 |  |
| GDC-810 | 9.6E-10 | 6.6E-10 | -96 | 2 |  |
| GDC-927 | 4.4E-11 | 4.3E-11 | -97 | 2 |  |
| GDC-9545 | 3.2E-11 | 1.8E-11 | -101 | 2 | SERDs |
| pomalidomide | 1.0E-06 | 0.0 | 0 | 0 |  |
| GNE-7388 | 5.3E-10 | 6.9E-11 | -69 | 3 |  |
| GNE-8506 | 7.3E-10 | 5.5E-11 | -93 | 1 |  |
| GNE-8093 | 8.8E-12 | 1.5E-12 | -57 | 1 |  |
| GNE-2681 | 2.9E-12 | 1.0E-12 | -92 | 1 |  |
| GNE-3340 | 6.8E-12 | 6.6E-12 | -61 | 1 | CRBN |
| RG-7112 | 1.0E-06 | 0.0 | 0 | 0 |  |
| GNE-3144 | 1.0E-06 | 0.0 | 0 | 0 |  |
| GNE-3183 | 3.8E-08 | 3.9E-08 | -60 | 32 |  |
| GNE-3185 | 1.7E-08 | 7.9E-09 | -47 | 17 | MDM2 |
| XB2m54 analog | 1.0E-06 | 0.0 | 0 | 0 |  |
| GNE-5583 | 4.5E-09 |  | -78 |  |  |
| GNE-0079 | 1.6E-09 |  | -103 |  |  |
| GNE-0080 | 5.0E-09 |  | -105 |  |  |
| GNE-0081 | 6.4E-09 |  | -93 |  |  |
| GNE-0993 | 3.7E-09 |  | -102 |  |  |
| GNE-0994 | 1.9E-08 |  | -90 |  |  |
| GNE-1567 • | 8.0E-10 |  | -102 |  |  |
| GNE-4182 | 2.4E-09 |  | -100 |  |  |
| GNE-6734 | 1.0E-09 |  | -101 |  |  |
| GNE-6735 | 6.9E-10 |  | -102 |  |  |
| GNE-6736 | 2.8E-10 |  | -100 |  |  |
| GNE-6746 • | 7.2E-10 |  | -100 |  |  |
| GNE-7530 | 1.8E-09 | 9.5E-10 | -102 | 4 |  |
| GNE-7648 | 8.5E-10 |  | -101 |  |  |
| GNE-0356 | 3.6E-07 | 2.3E-07 | -92 | 17 |  |
| GNE-4081 | 1.4E-09 | 6.9E-10 | -69 | 2 |  |
| GNE-4082 | 2.5E-09 | 6.0E-10 | -80 | 2 |  |
| GNE-1513 • | 4.4E-10 |  | -105 |  |  |
| GNE-9535 • * | 9.1E-10 | 1.1E-09 | -103 | 4 |  |
| GNE-9536 • x | 1.3E-09 | 1.0E-09 | -95 | 4 |  |
| GNE-9537 • + | 1.9E-09 | 1.4E-09 | -82 | 1 |  |
| GNE-8946 | 3.8E-07 | 1.3E-07 | -100 | 0 | XIAP |
| GDC-0152 analog.1 | 1.0E-06 | 0.0 | 0 | 0 |  |
| GDC-0152 analog.2 | 1.0E-06 | 0.0 | 0 | 0 |  |
| GDC-0152 analog.3 | 1.0E-06 | 0.0 | 0 | 0 |  |
| GDC-0152 analog.4 | 1.0E-06 | 0.0 | 0 | 0 |  |
| GNE-4863 • | 2.5E-09 | 2.2E-10 | -94 | 3 |  |
| GNE-9297 • | 6.8E-09 | 2.6E-09 | -104 | 2 |  |
| GNE-5472 • | 2.1E-10 | 3.1E-10 | -105 | 1 | pan-IAP |
| GNE-5792 • | 4.2E-10 | 3.0E-11 | -106 | 1 |  |
| GNE-7387 • | 1.1E-09 | 3.1E-10 | -105 | 1 |  |
| GNE-8443 | 1.6E-09 | 5.7E-10 | -71 | 3 |  |
| GNE-8444 • | 2.4E-10 | 4.9E-11 | -76 | 4 |  |
| GNE-8445 | 1.1E-09 | 2.4E-10 | -84 | 1 |  |
| GNE-8446 • | 4.1E-10 | 4.3E-11 | -68 | 4 |  |
| GNE-1574 | 1.8E-10 | 2.0E-11 | -107 | 1 |  |
| GNE-2003 | 4.9E-10 | 3.9E-11 | -74 | 1 | VHL |

• : evaluated in ternary complex SPR studies (Table 2)

* x + : series of XIAP/ERa-CIDEs; same endoxifen/linker, different XIAP ligand affinities
