## Supplementary material for "Engineering ERα degraders with pleiotropic ubiquitin ligase ligands maximizes therapeutic efficacy by co-opting distinct effector ligases": Table 2

| **ERa ligand/CIDE** | **IF IC50** | **IF Dmax** | **SPR assay ligase** | **t 1/2 (s) average** | **t 1/2 (s) STDEV** | **KD (nM) average** | **KD (nM) STDEV** | **ka (1/Ms) average** | **kd (1/s) average** | **comments** |
| --- | --- | --- | --- | --- | --- | --- | --- | --- | --- | --- |
| GNE-1513 | 4.40E-10 | ***-105*** | XIAP BIR2 | ***482.0*** | 137.0 | ***2.6*** | 1.1 | 6.32E+05 | 1.52E-03 |  |
| GNE-9536 x | 1.30E-09 | ***-95*** | XIAP BIR2 | ***39.8*** | 7.9 | ***21.3*** | 8.2 | 9.13E+05 | 1.80E-02 |  |
| GNE-9537 + | 1.90E-09 | ***-82*** | XIAP BIR2 | ***11.9*** | 4.0 | ***30.7*** | 13.3 | 2.29E+06 | 6.64E-02 |  |
| GNE-6746 | 7.20E-10 | ***-100*** | XIAP BIR2 | ***224.0*** | 70.9 | ***4.0*** | 1.2 | 8.35E+05 | 3.35E-03 |  |
| GNE-9535 * | 9.10E-10 | ***-103*** | XIAP BIR2 | ***76.5*** | 30.5 | ***15.0*** | 7.5 | 7.57E+05 | 1.04E-02 |  |
| GNE-1567 | 8.00E-10 | ***-102*** | XIAP BIR2 | ***290.4*** | 83.2 | ***5.2*** | 2.0 | 5.48E+05 | 2.54E-03 |  |
| endoxifen | 3.70E-11 | ***-69*** | XIAP BIR2 | ***0*** |  | ***> 100*** |  |  |  | no complex detected |
| GNE-4863 | 2.50E-09 | ***-94*** | XIAP BIR3 | ***634.7*** | 171.6 | ***16.7*** | 5.0 | 7.04E+04 | 1.16E-03 |  |
| GNE-5472 | 2.10E-10 | ***-105*** | XIAP BIR3 | ***1055.0*** | 413.6 | ***12.6*** | 3.4 | 5.78E+04 | 7.40E-04 |  |
| GNE-9297 | 6.80E-09 | ***-104*** | XIAP BIR3 | ***111.1*** | 26.1 | ***134.0*** | 29.3 | 4.84E+04 | 6.47E-03 |  |
| endoxifen | 3.70E-11 | ***-69*** | XIAP BIR3 | ***0*** |  | ***> 1000*** |  |  |  | no complex detected |
| GNE-5792 | 4.20E-10 | ***-106*** | VHL | ***87.4*** | 16.3 | ***21.6*** | 4.3 | 3.76E+05 | 8.11E-03 |  |
| GNE-7387 | 1.10E-09 | ***-105*** | VHL | ***163.7*** | 16.2 | ***13.6*** | 2.3 | 3.16E+05 | 4.27E-03 |  |
| GNE-8446 | 4.10E-10 | ***-68*** | VHL | ***0*** |  | ***> 100*** |  |  |  | no complex detected |
| GNE-8444 | 2.40E-10 | ***-76*** | VHL | ***0*** |  | ***> 100*** |  |  |  | no complex detected |
| endoxifen | 3.70E-11 | ***-69*** | VHL | ***0*** |  | ***> 100*** |  |  |  | no complex detected |
|  |  |  |  | ***Fig. S2C correlation*** |  | ***Figure 2C correlation*** |  |  |  |  |

* x + : series of XIAP/ERa-CIDEs; same endoxifen/linker, different XIAP ligand affinities
