## Supplementary material for "Engineering ERα degraders with pleiotropic ubiquitin ligase ligands maximizes therapeutic efficacy by co-opting distinct effector ligases": Synthetic procedures

**General Methods.**

All commercial solvents and reagents were used without additional purification unless indicated otherwise. 1H NMR spectra were measured on Bruker Avance III 300, 400, or 500 MHz spectrometers. Chemical shifts (in ppm) were referenced to tetramethylsilane as an internal standard (δ = 0 ppm). Reaction progress was monitored by either a Shimadzu LCMS/UV system with an LC-30 AD solvent pump, Sil-30 AC autosampler, 2020 MS, SPDM30A UV detector, and CTO-20A column oven, using 2−98% acetonitrile/0.1% formic acid (or 0.01% ammonia) over 2.5 min or a Waters Acquity LCMS system using 2−98% acetonitrile/0.1% formic acid (or 0.1% ammonia) over 2 min. Flash column chromatography purifications were performed using a Teledyne Isco Combiflash Rf and Silicycle HP columns. Reverse-phase purification was done on a Phenomenex Gemini-NX C18 (30 × 100 mm, 5 μm) with a gradient of 5−95% acetonitrile/water (with 0.1% NH_4_OH or 0.1% formic acid) at 60 mL/min over 10 min. Preparative SFC separations were carried out on a PIC Solutions instrument. High-resolution mass spectrometry (HRMS) of final compounds was obtained on a Thermo UHPLC/QE with a Thermo-Q Exactive mass spectrometry detector using ESI ionization, following elution on an Acquity BEH C18 stationary phase (2.1 mm × 50 mm; 1.7 μm particle size) using a gradient of water/acetonitrile (3−97% over 7 min with 0.1% formic acid in both phases). Unless stated otherwise, analytical purity was >95% as determined by LCMS using UV 254 nm detection. Fulvestrant (CAS# 129453-61-8), endoxifen (CAS# 112093-28-4), brilanestrant (GDC-0810, CAS# 1365888-06-7), GDC-0927 (CAS# 1642297-01-5) and giredestrant (GDC-9545, CAS# 1953133-47-5) are commercially available.

**Synthesis of XIAP BIR2 ligand (XB2m54 analog)**

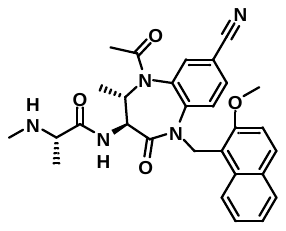

Synthesis of XIAP BIR2 ligand XB2m54 analog was reported here (compound **33**): Kester, R. F. et al Optimization of Benzodiazepinones as Selective Inhibitors of the X-Linked Inhibitor of Apoptosis Protein (XIAP) Second Baculovirus IAP Repeat (BIR2) Domain. *J. Med. Chem.* **56**, 7788-7803 (2013).

**Synthesis of degrader GNE-1513**

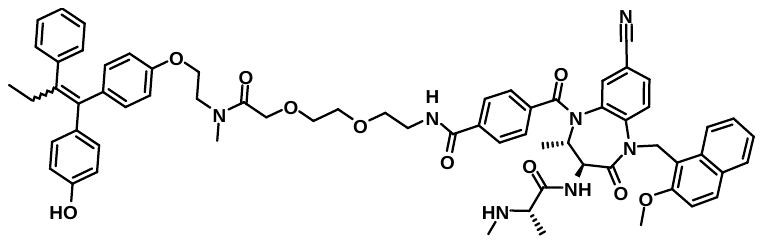

Synthesis of degrader GNE-1513 was reported here (Compound P1): WO2017201449A1

**Synthesis of degrader GNE-1567**

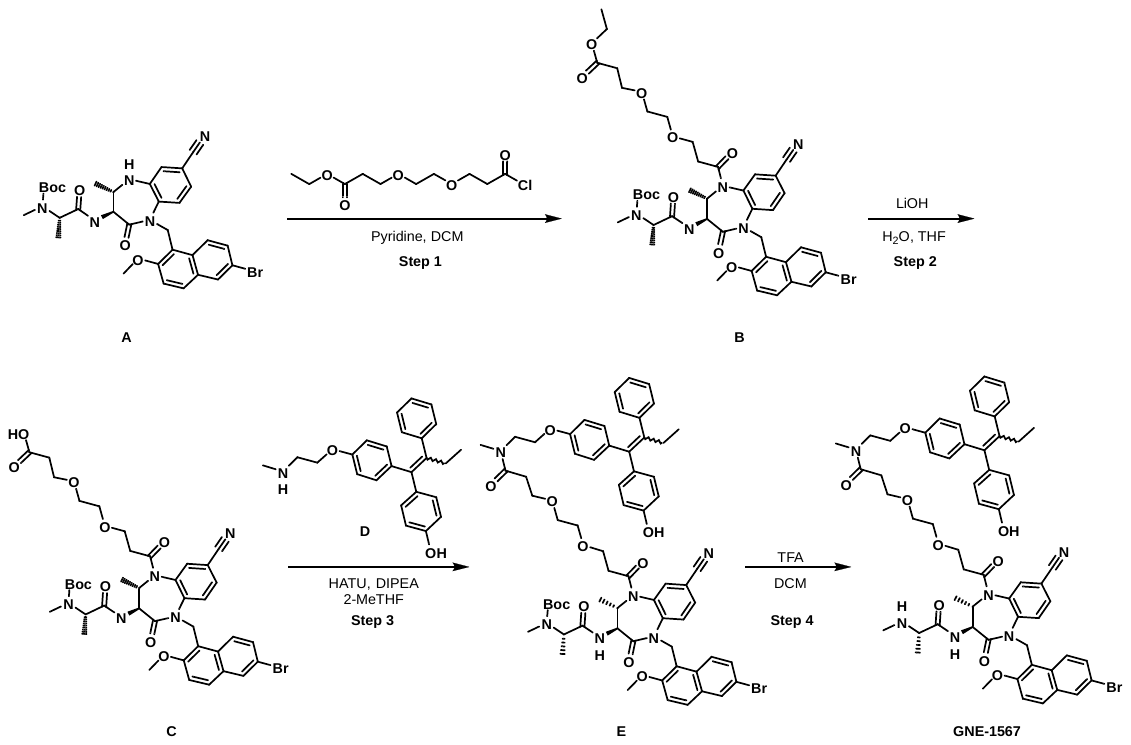

**Step 1: ethyl 3-(2-(3-((2R,3R)-5-((6-bromo-2-methoxynaphthalen-1-yl)methyl)-3-((R)-2-((tert-butoxycarbonyl)(methyl)amino)propanamido)-8-cyano-2-methyl-4-oxo-2,3,4,5-tetrahydro-1H-benzo[b][1,4]diazepin-1-yl)-3-oxopropoxy)ethoxy)propanoate (B)**

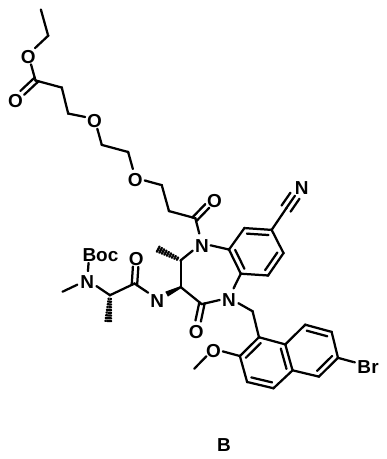

To a solution of tert-butyl N-[(1S)-2-[[(3S,4S)-1-[(6-bromo-2-methoxy-1-naphthyl)methyl]-7-cyano-4-methyl-2-oxo-4,5-dihydro-3H-1,5-benzodiazepin-3-yl]amino]-1-methyl-2-oxo-ethyl]-N-methyl-carbamate (**A**) (100 mg, 0.154 mmol, 1 eq.) at 0 ^o^C in DCM (1.5 mL) was added pyridine (60.8 mg, 0.768 mmol, 5 eq.) followed by ethyl 3-[2-(3-chloro-3-oxo-propoxy)ethoxy]propanoate (97.3 mg, 0.385 mmol, 2.5 eq.) dropwise. The reaction was stirred at 0 ºC for 1h, then allowed to warm to room temperature and stirred for an additional 15 h, then quenched by the addition of saturated NaHCO_3_ and extracted with DCM. The organic layers were combined, dried with sodium sulfate, and concentrated. The crude residue was purified by flash chromatography (3:1 iPrOAc:MeOH in heptanes) to afford the title compound (60.0 mg, 45%) of a yellow oil.

**Step 2: 3-(2-(3-((2R,3R)-5-((6-bromo-2-methoxynaphthalen-1-yl)methyl)-3-((R)-2-((tert-butoxycarbonyl)(methyl)amino)propanamido)-8-cyano-2-methyl-4-oxo-2,3,4,5-tetrahydro-1H-benzo[b][1,4]diazepin-1-yl)-3-oxopropoxy)ethoxy)propanoic acid (C)**

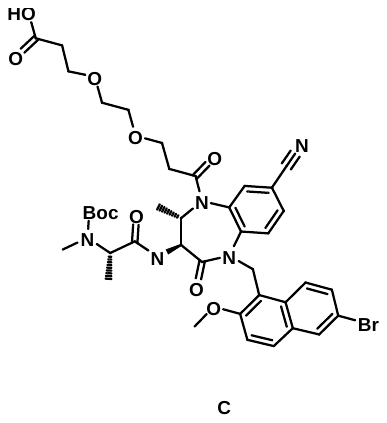

To a solution of ethyl 3-(2-(3-((2R,3R)-5-((6-bromo-2-methoxynaphthalen-1-yl)methyl)-3-((R)-2-((tert-butoxycarbonyl)(methyl)amino)propanamido)-8-cyano-2-methyl-4-oxo-2,3,4,5-tetrahydro-1H-benzo[b][1,4]diazepin-1-yl)-3-oxopropoxy)ethoxy)propanoate (**B**) (60.0 mg, 0.0692 mmol, 1 eq.) in THF (0.2 mL) and water (0.1 mL) was added LiOH (6.63 mg, 0.277 mmol, 4 eq.). The reaction was stirred at room temperature for 5 h. The solution was acidified to pH = 1 with 2N HCl. The solution was extracted with DCM 3x. The organic layers were combined, dried with Na_2_SO_4_ and concentrated. The crude product was carried over to the next step.

**Step 3: tert-butyl ((R)-1-(((3R,4R)-1-((6-bromo-2-methoxynaphthalen-1-yl)methyl)-7-cyano-5-(3-(2-(3-((2-(4-(1-(4-hydroxyphenyl)-2-phenylbut-1-en-1-yl)phenoxy)ethyl)(methyl)amino)-3-oxopropoxy)ethoxy)propanoyl)-4-methyl-2-oxo-2,3,4,5-tetrahydro-1H-benzo[b][1,4]diazepin-3-yl)amino)-1-oxopropan-2-yl)(methyl)carbamate (E)**

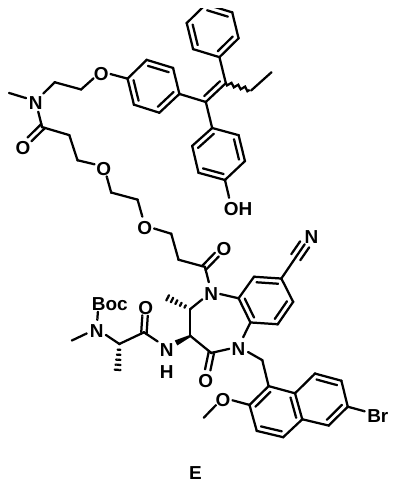

To a crude solution of 3-[2-[3-[(3S,4S)-1-[(6-bromo-2-methoxy-1-naphthyl)methyl]-3-[[(2S)-2-[tert-butoxycarbonyl(methyl)amino]propanoyl]amino]-7-cyano-4-methyl-2-oxo-3,4-dihydro-1,5-benzodiazepin-5-yl]-3-oxo-propoxy]ethoxy]propanoic acid (**C**) (39.0 mg, 0.0465 mmol, 1 eq.) in 2-MeTHF (1.4 mL) was added DIPEA (17.9 mg, 0.138 mmol, 2 eq.) and HATU (29.5 mg, 0.0761 mmol, 1.1 eq.) followed by 4-(1-(4-(2-(methylamino)ethoxy)phenyl)-2-phenylbut-1-en-1-yl)phenol (**D**) (18.2 mg, 0.0488 mmol, 1.05 eq.). The mixture was stirred at room temperature for 22h. Water was added and the solution was extracted 3x with EtOAc. The organic layers were combined then dried with sodium sulfate and concentrated to afford the crude title compound.

**Step 4: 3-(2-(3-((2R,3R)-5-((6-bromo-2-methoxynaphthalen-1-yl)methyl)-8-cyano-2-methyl-3-((R)-2-(methylamino)propanamido)-4-oxo-2,3,4,5-tetrahydro-1H-benzo[b][1,4]diazepin-1-yl)-3-oxopropoxy)ethoxy)-N-(2-(4-(1-(4-hydroxyphenyl)-2-phenylbut-1-en-1-yl)phenoxy)ethyl)-N-methylpropanamide (GNE-1567)**

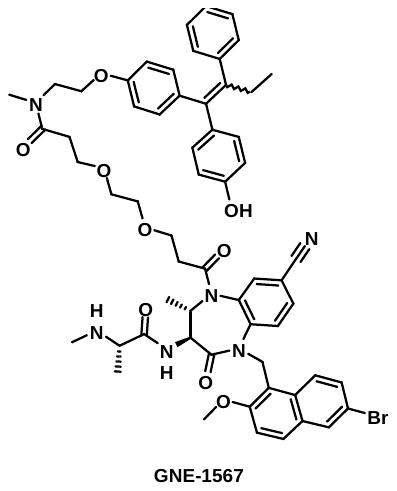

The crude tert-butyl ((R)-1-(((3R,4R)-1-((6-bromo-2-methoxynaphthalen-1-yl)methyl)-7-cyano-5-(3-(2-(3-((2-(4-(1-(4-hydroxyphenyl)-2-phenylbut-1-en-1-yl)phenoxy)ethyl)(methyl)amino)-3-oxopropoxy)ethoxy)propanoyl)-4-methyl-2-oxo-2,3,4,5-tetrahydro-1H-benzo[b][1,4]diazepin-3-yl)amino)-1-oxopropan-2-yl)(methyl)carbamate (**E**) was dissolved in DCM (1.5 mL). TFA (0.12 mL) was added and the reaction was stirred at room temperature for 2h. The solution was concentrated under vacuum, diluted with saturated NaHCO_3_ and extracted 2x with DCM. The organic layers were combined, dried with sodium sulfated and concentrated. Purified by reverse-phase HPLC to afford the title compound (14.7 mg, 19% over 3-step) as a mixture of E/Z isomers. MS (ESI): [M+H]^+^ = 1096. 1H NMR (400 MHz, DMSO) δ 9.40 (s, 0.5H), 9.15 (s, 0.5H), 8.50 (d, J = 8.9 Hz, 1H), 8.11 – 8.03 (m, 2H), 8.01 (ddt, J = 8.1, 6.0, 1.8 Hz, 1H), 7.94 (d, J = 9.1 Hz, 1H), 7.85 – 7.78 (m, 2H), 7.40 (dd, J = 9.1, 2.1 Hz, 1H), 7.37 – 7.28 (m, 1H), 7.22 – 7.12 (m, 2H), 7.12 – 7.03 (m, 4H), 7.00 – 6.94 (m, 1H), 6.90 (d, J = 9.0 Hz, 1H), 6.75 (d, J = 8.7 Hz, 1H), 6.69 (dd, J = 8.7, 3.4 Hz, 1H), 6.61 – 6.51 (m, 2H), 6.39 (d, J = 8.7 Hz, 1H), 5.93 (d, J = 14.9 Hz, 1H), 5.35 (d, J = 14.9 Hz, 1H), 4.81 (dt, J = 12.3, 6.2 Hz, 1H), 4.15 – 4.01 (m, 2H), 3.98 – 3.86 (m, 1H), 3.85 – 3.81 (m, 3H), 3.68 – 3.49 (m, 2H), 3.45 – 3.38 (m, 1H), 3.36 (s, 2H), 3.29 – 3.19 (m, 2H), 3.12 – 2.94 (m, 3H), 2.86 (br, 1H), 2.60 (dd, J = 10.7, 6.8 Hz, 1H), 2.39 (dd, J = 13.6, 7.6 Hz, 3H), 2.21 (s, 3H), 1.94 (s, 1H), 1.54 – 1.46 (m, 1H), 1.11 (d, J = 6.9 Hz, 3H), 0.90 (dd, J = 6.2, 2.1 Hz, 3H), 0.88 – 0.78 (m, 3H).

**Synthesis of XIAP BIR3 ligand (GNE-7463)**

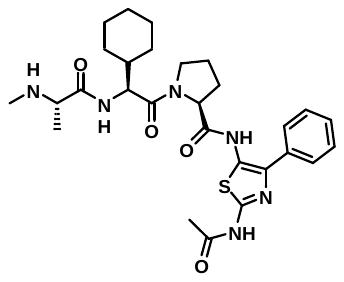

Synthesis of XIAP BIR3 ligand GNE-7463 was reported here (Example 144): US2010256115A1

**Synthesis of degrader GNE-5472**

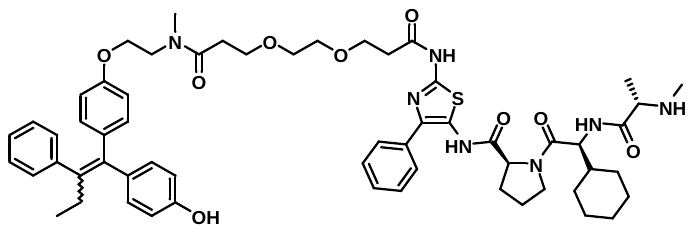

Synthesis of degrader GNE-5472 was reported here (Compound EC1): WO2020086858A1

**Synthesis of degrader GNE-5792**

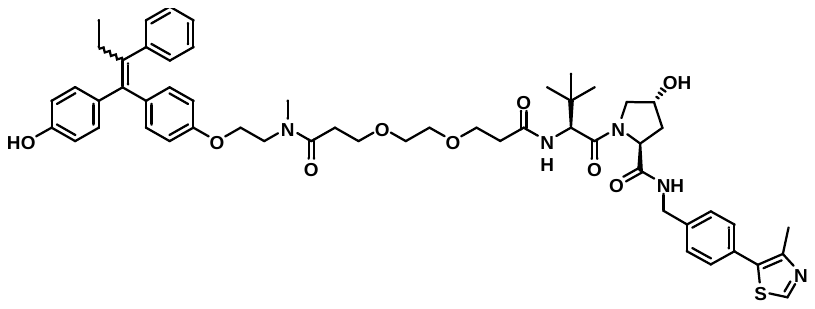

Synthesis of degrader GNE-5792 was reported here: Dragovich, P. S. et al. Antibody-mediated delivery of chimeric protein degraders which target estrogen receptor alpha (ERα). *Bioorganic Med. Chem. Lett.* **30**, 126907 (2020).

**Synthesis of degrader GNE-7387**

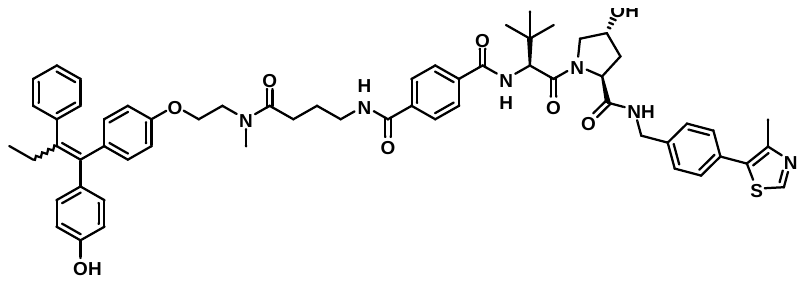

Synthesis of degrader GNE-7387 was reported here (Compound EC4): WO2020086858A1

**Synthesis of degrader GNE-9535**

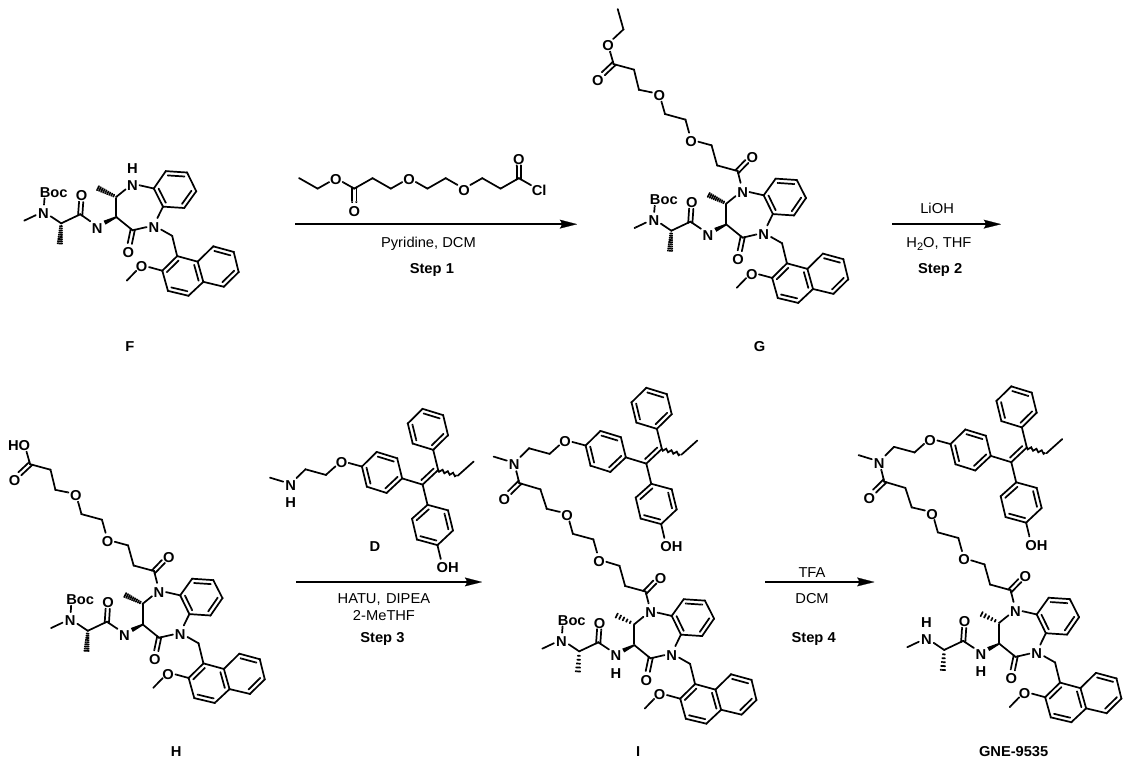

**Step 1: ethyl 3-(2-(3-((2R,3R)-3-((R)-2-((tert-butoxycarbonyl)(methyl)amino)propanamido)-5-((2-methoxynaphthalen-1-yl)methyl)-2-methyl-4-oxo-2,3,4,5-tetrahydro-1H-benzo[b][1,4]diazepin-1-yl)-3-oxopropoxy)ethoxy)propanoate (G)**

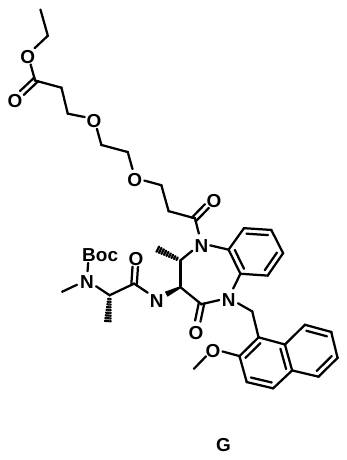

To a solution of tert-butyl ((R)-1-(((3R,4R)-1-((2-methoxynaphthalen-1-yl)methyl)-4-methyl-2-oxo-2,3,4,5-tetrahydro-1H-benzo[b][1,4]diazepin-3-yl)amino)-1-oxopropan-2-yl)(methyl)carbamate (**F**) (50.0 mg, 0.0915 mmol, 1 eq.) at 0 ^o^C in DCM (0.9 mL) was added pyridine (36.2 mg, 0.457 mmol, 5 eq.) followed by ethyl 3-[2-(3-chloro-3-oxo-propoxy)ethoxy]propanoate (57.8 mg, 0.229 mmol, 2.5 eq.) dropwise. The reaction was stirred at 0 ºC for 1h, then allowed to warm to room temperature and stirred for an additional 15 h, then quenched by the addition of saturated NaHCO_3_ and extracted with DCM. The organic layers were combined, dried with sodium sulfate, and concentrated to afford the desired product (65.0 mg) as a crude oil. The crude product was carried over to the next step.

**Step 2: 3-(2-(3-((2R,3R)-3-((R)-2-((tert-butoxycarbonyl)(methyl)amino)propanamido)-5-((2-methoxynaphthalen-1-yl)methyl)-2-methyl-4-oxo-2,3,4,5-tetrahydro-1H-benzo[b][1,4]diazepin-1-yl)-3-oxopropoxy)ethoxy)propanoic acid (H)**

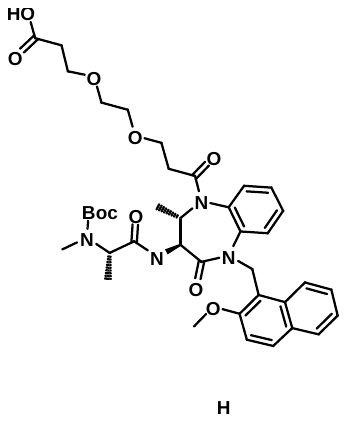

To a solution of ethyl 3-(2-(3-((2R,3R)-3-((R)-2-((tert-butoxycarbonyl)(methyl)amino)propanamido)-5-((2-methoxynaphthalen-1-yl)methyl)-2-methyl-4-oxo-2,3,4,5-tetrahydro-1H-benzo[b][1,4]diazepin-1-yl)-3-oxopropoxy)ethoxy)propanoate (**G**) (65.0 mg, 0.0852 mmol, 1 eq.) in THF (0.2 mL) and water (0.1 mL) was added LiOH (11.1 mg, 0.464 mmol, 5.5 eq.). The reaction was stirred at room temperature for 5 h. The solution was acidified to pH = 1 with 2N HCl. The solution was extracted with DCM 3x. The organic layers were combined, dried with Na_2_SO_4_ and concentrated. The crude product was carried over to the next step.

**Step 3: tert-butyl ((R)-1-(((2R,3R)-1-(3-(2-(3-((2-(4-(1-(4-hydroxyphenyl)-2-phenylbut-1-en-1-yl)phenoxy)ethyl)(methyl)amino)-3-oxopropoxy)ethoxy)propanoyl)-5-((2-methoxynaphthalen-1-yl)methyl)-2-methyl-4-oxo-2,3,4,5-tetrahydro-1H-benzo[b][1,4]diazepin-3-yl)amino)-1-oxopropan-2-yl)(methyl)carbamate (I)**

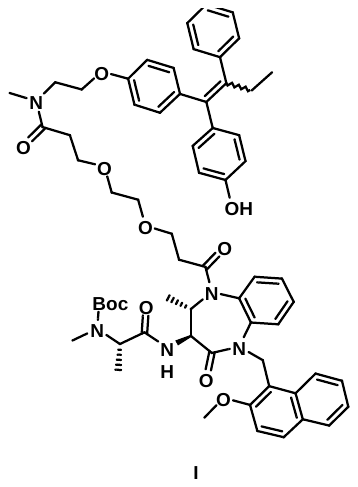

To a crude solution of 3-(2-(3-((2R,3R)-3-((R)-2-((tert-butoxycarbonyl)(methyl)amino)propanamido)-5-((2-methoxynaphthalen-1-yl)methyl)-2-methyl-4-oxo-2,3,4,5-tetrahydro-1H-benzo[b][1,4]diazepin-1-yl)-3-oxopropoxy)ethoxy)propanoic acid (**H**) (39.0 mg, 0.0531 mmol, 1 eq.) in 2-MeTHF (0.9 mL) was added DIPEA (22.0 mg, 0.170 mmol, 2 eq.) and HATU (36.4 mg, 0.0937 mmol, 1.1 eq.) followed by 4-(1-(4-(2-(methylamino)ethoxy)phenyl)-2-phenylbut-1-en-1-yl)phenol (**D**) (20.8 mg, 0.0557 mmol, 1.05 eq.). The mixture was stirred at room temperature for 22h. Water was added and the solution was extracted 3x with EtOAc. The organic layers were combined then dried with sodium sulfate and concentrated to afford the crude title compound.

**Step 4:** **N-(2-(4-(1-(4-hydroxyphenyl)-2-phenylbut-1-en-1-yl)phenoxy)ethyl)-3-(2-(3-((2R,3R)-5-((2-methoxynaphthalen-1-yl)methyl)-2-methyl-3-((R)-2-(methylamino)propanamido)-4-oxo-2,3,4,5-tetrahydro-1H-benzo[b][1,4]diazepin-1-yl)-3-oxopropoxy)ethoxy)-N-methylpropanamide (GNE-9535)**

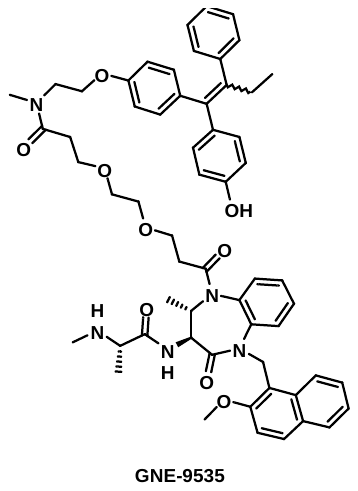

The crude tert-butyl ((R)-1-(((2R,3R)-1-(3-(2-(3-((2-(4-(1-(4-hydroxyphenyl)-2-phenylbut-1-en-1-yl)phenoxy)ethyl)(methyl)amino)-3-oxopropoxy)ethoxy)propanoyl)-5-((2-methoxynaphthalen-1-yl)methyl)-2-methyl-4-oxo-2,3,4,5-tetrahydro-1H-benzo[b][1,4]diazepin-3-yl)amino)-1-oxopropan-2-yl)(methyl)carbamate (I) was dissolved in DCM (0.5 mL). TFA (0.14 mL) was added and the reaction was stirred at room temperature for 2h. The solution was concentrated under vacuum, diluted with saturated NaHCO_3_ and extracted 2x with DCM. The organic layers were combined, dried with sodium sulfated and concentrated. Purified by reverse-phase HPLC to afford the title compound (25.9 mg, 28% over 4-step) as a mixture of E/Z isomers. MS (ESI): [M+H]^+^ = 990.6. 1H NMR (400 MHz, DMSO) δ 9.40 (s, 0.5H), 9.21 (d, J = 8.8 Hz, 1H), 9.15 (s, 0.5H), 8.86 – 8.60 (m, 3H), 8.06 – 7.99 (m, 1H), 7.87 (d, J = 8.2 Hz, 1H), 7.81 – 7.72 (m, 2H), 7.55 – 7.45 (m, 1H), 7.36 – 7.29 (m, 2H), 7.29 – 7.19 (m, 2H), 7.19 – 7.12 (m, 3H), 7.12 – 7.03 (m, 4H), 7.00 – 6.94 (m, 1H), 6.94 – 6.88 (m, 1H), 6.79 – 6.72 (m, 1H), 6.72 – 6.66 (m, 1H), 6.63 – 6.52 (m, 2H), 6.43 – 6.35 (m, 1H), 5.93 (d, J = 14.6 Hz, 1H), 5.33 (d, J = 14.7 Hz, 1H), 4.83 (dq, J = 12.2, 6.1 Hz, 1H), 4.14 – 4.01 (m, 2H), 3.99 – 3.76 (m, 5H), 3.74 – 3.50 (m, 3H), 3.21 – 3.11 (m, 2H), 3.09 – 2.92 (m, 3H), 2.84 (d, J = 29.9 Hz, 2H), 2.70 – 2.52 (m, 2H), 2.44 – 2.34 (m, 2H), 1.47 (d, J = 6.9 Hz, 3H), 1.37 – 1.21 (m, 1H), 0.92 (dd, J = 6.3, 1.7 Hz, 3H), 0.89 – 0.78 (m, 3H).

**Synthesis of degrader GNE-9536**

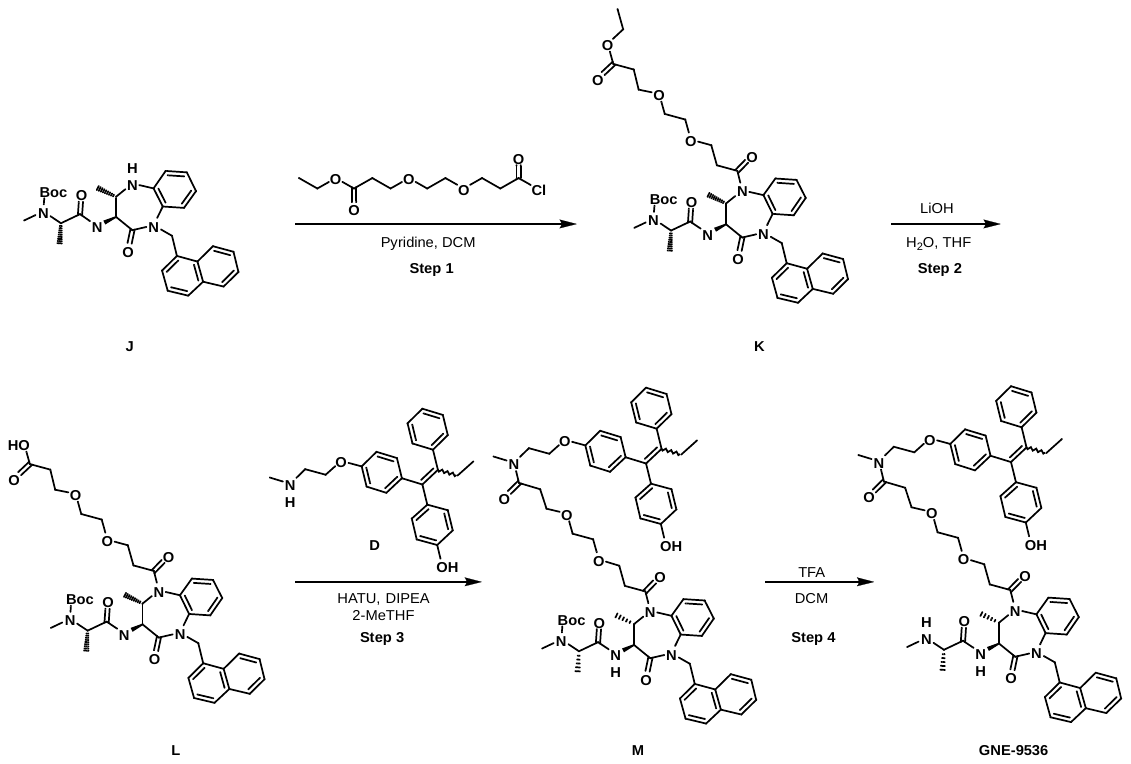

**Step 1: ethyl 3-(2-(3-((2R,3R)-3-((R)-2-((tert-butoxycarbonyl)(methyl)amino)propanamido)-2-methyl-5-(naphthalen-1-ylmethyl)-4-oxo-2,3,4,5-tetrahydro-1H-benzo[b][1,4]diazepin-1-yl)-3-oxopropoxy)ethoxy)propanoate (K)**

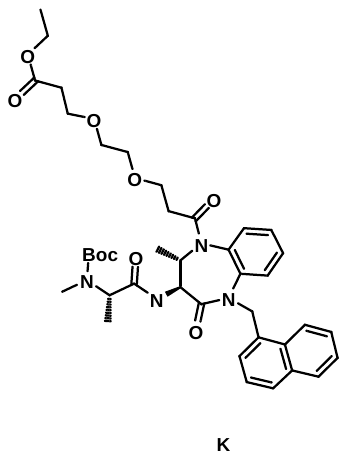

To a solution of tert-butyl methyl((R)-1-(((3R,4R)-4-methyl-1-(naphthalen-1-ylmethyl)-2-oxo-2,3,4,5-tetrahydro-1H-benzo[b][1,4]diazepin-3-yl)amino)-1-oxopropan-2-yl)carbamate (**J**) (50.0 mg, 0.0968 mmol, 1 eq.) at 0 ^o^C in DCM (1.0 mL) was added pyridine (38.3 mg, 0.484 mmol, 5 eq.) followed by ethyl 3-[2-(3-chloro-3-oxo-propoxy)ethoxy]propanoate (61.2 mg, 0.242 mmol, 2.5 eq.) dropwise. The reaction was stirred at 0 ºC for 1h, then allowed to warm to room temperature and stirred for an additional 15 h, then quenched by the addition of saturated NaHCO_3_ and extracted with DCM. The organic layers were combined, dried with sodium sulfate, and concentrated. The crude residue was purified by flash chromatography (3:1 iPrOAc:MeOH in heptanes) to afford the title compound (60.0 mg, 85%) of a yellow oil.

**Step 2: 3-(2-(3-((2R,3R)-3-((R)-2-((tert-butoxycarbonyl)(methyl)amino)propanamido)-2-methyl-5-(naphthalen-1-ylmethyl)-4-oxo-2,3,4,5-tetrahydro-1H-benzo[b][1,4]diazepin-1-yl)-3-oxopropoxy)ethoxy)propanoic acid (L)**

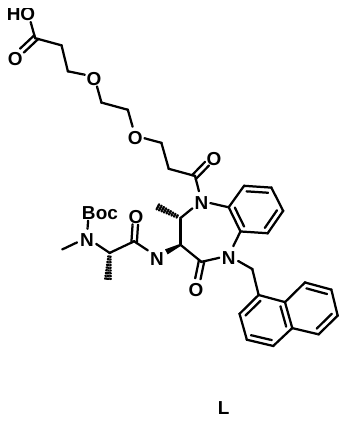

To a solution of ethyl 3-(2-(3-((2R,3R)-3-((R)-2-((tert-butoxycarbonyl)(methyl)amino)propanamido)-2-methyl-5-(naphthalen-1-ylmethyl)-4-oxo-2,3,4,5-tetrahydro-1H-benzo[b][1,4]diazepin-1-yl)-3-oxopropoxy)ethoxy)propanoate (**K**) (65.0 mg, 0.0887 mmol, 1 eq.) in THF (0.2 mL) and water (0.1 mL) was added LiOH (11.4 mg, 0.475 mmol, 5.4 eq.). The reaction was stirred at room temperature for 5 h. The solution was acidified to pH = 1 with 2N HCl. The solution was extracted with DCM 3x. The organic layers were combined, dried with Na_2_SO_4_ and concentrated. The crude product was carried over to the next step.

**Step 3: tert-butyl ((R)-1-(((2R,3R)-1-(3-(2-(3-((2-(4-(1-(4-hydroxyphenyl)-2-phenylbut-1-en-1-yl)phenoxy)ethyl)(methyl)amino)-3-oxopropoxy)ethoxy)propanoyl)-2-methyl-5-(naphthalen-1-ylmethyl)-4-oxo-2,3,4,5-tetrahydro-1H-benzo[b][1,4]diazepin-3-yl)amino)-1-oxopropan-2-yl)(methyl)carbamate (M)**

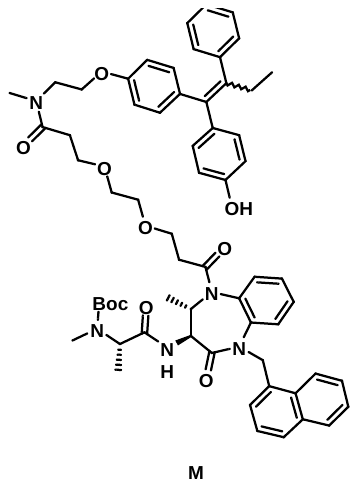

To a crude solution of 3-(2-(3-((2R,3R)-3-((R)-2-((tert-butoxycarbonyl)(methyl)amino)propanamido)-2-methyl-5-(naphthalen-1-ylmethyl)-4-oxo-2,3,4,5-tetrahydro-1H-benzo[b][1,4]diazepin-1-yl)-3-oxopropoxy)ethoxy)propanoic acid (**L**) (39.0 mg, 0.0553 mmol, 1 eq.) in 2-MeTHF (1.8 mL) was added DIPEA (22.9 mg, 0.177 mmol, 2 eq.) and HATU (37.9 mg, 0.0976 mmol, 1.1 eq.) followed by 4-(1-(4-(2-(methylamino)ethoxy)phenyl)-2-phenylbut-1-en-1-yl)phenol (**D**) (21.7 mg, 0.0581 mmol, 1.05 eq.). The mixture was stirred at room temperature for 22h. Water was added and the solution was extracted 3x with EtOAc. The organic layers were combined then dried with sodium sulfate and concentrated to afford the crude title compound.

**Step 4:** **N-(2-(4-(1-(4-hydroxyphenyl)-2-phenylbut-1-en-1-yl)phenoxy)ethyl)-N-methyl-3-(2-(3-((2R,3R)-2-methyl-3-((R)-2-(methylamino)propanamido)-5-(naphthalen-1-ylmethyl)-4-oxo-2,3,4,5-tetrahydro-1H-benzo[b][1,4]diazepin-1-yl)-3-oxopropoxy)ethoxy)propanamide (GNE-9536)**

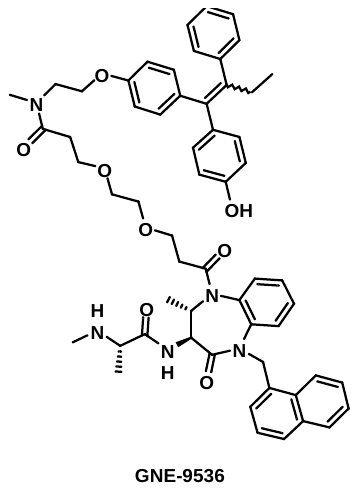

The crude tert-butyl ((R)-1-(((2R,3R)-1-(3-(2-(3-((2-(4-(1-(4-hydroxyphenyl)-2-phenylbut-1-en-1-yl)phenoxy)ethyl)(methyl)amino)-3-oxopropoxy)ethoxy)propanoyl)-2-methyl-5-(naphthalen-1-ylmethyl)-4-oxo-2,3,4,5-tetrahydro-1H-benzo[b][1,4]diazepin-3-yl)amino)-1-oxopropan-2-yl)(methyl)carbamate (**M**) was dissolved in DCM (0.6 mL). TFA (0.15 mL) was added and the reaction was stirred at room temperature for 2h. The solution was concentrated under vacuum, diluted with saturated NaHCO_3_ and extracted 2x with DCM. The organic layers were combined, dried with sodium sulfated and concentrated. Purified by reverse-phase HPLC to afford the title compound (19.5 mg, 23% over 3-step) as a mixture of E/Z isomers. MS (ESI): [M+H]^+^ = 960.5. 1H NMR (400 MHz, DMSO) δ 9.40 (s, 0.5H), 9.21 (d, J = 8.8 Hz, 1H), 9.15 (s, 0.5H), 8.85 – 8.63 (m, 3H), 8.06 – 7.99 (m, 1H), 7.87 (d, J = 8.2 Hz, 1H), 7.81 – 7.72 (m, 2H), 7.55 – 7.45 (m, 1H), 7.36 – 7.29 (m, 2H), 7.29 – 7.19 (m, 2H), 7.19 – 7.12 (m, 3H), 7.12 – 7.03 (m, 4H), 7.00 – 6.94 (m, 1H), 6.94 – 6.88 (m, 1H), 6.79 – 6.72 (m, 1H), 6.72 – 6.66 (m, 1H), 6.63 – 6.52 (m, 2H), 6.43 – 6.35 (m, 1H), 5.93 (d, J = 14.6 Hz, 1H), 5.33 (d, J = 14.7 Hz, 1H), 4.83 (dq, J = 12.2, 6.1 Hz, 1H), 4.14 – 4.01 (m, 2H), 3.99 – 3.76 (m, 5H), 3.74 – 3.50 (m, 3H), 3.21 – 3.11 (m, 2H), 3.09 – 2.92 (m, 3H), 2.84 (d, J = 29.9 Hz, 2H), 2.70 – 2.52 (m, 2H), 2.44 – 2.34 (m, 2H), 1.47 (d, J = 6.9 Hz, 3H), 1.37 – 1.21 (m, 1H), 0.92 (dd, J = 6.3, 1.7 Hz, 3H), 0.89 – 0.78 (m, 3H).

**Synthesis of degrader GNE-9537**

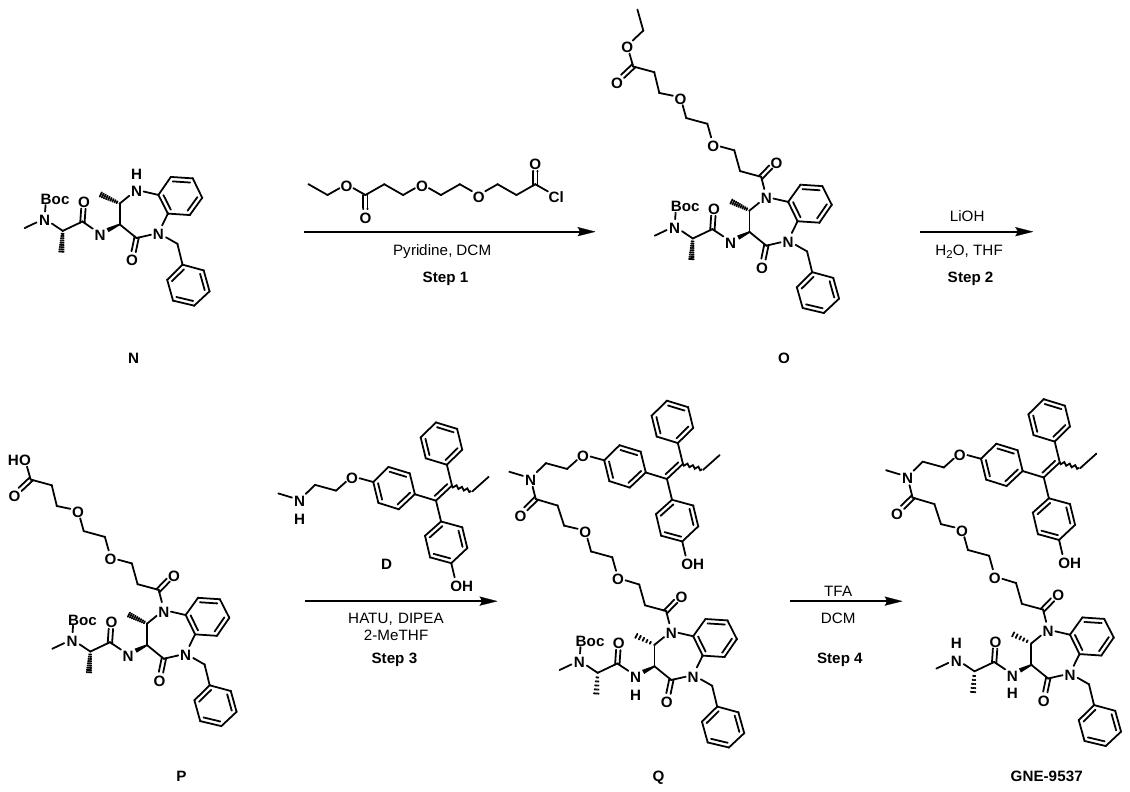

**Step 1: ethyl 3-(2-(3-((2R,3R)-5-benzyl-3-((R)-2-((tert-butoxycarbonyl)(methyl)amino)propanamido)-2-methyl-4-oxo-2,3,4,5-tetrahydro-1H-benzo[b][1,4]diazepin-1-yl)-3-oxopropoxy)ethoxy)propanoate (O)**

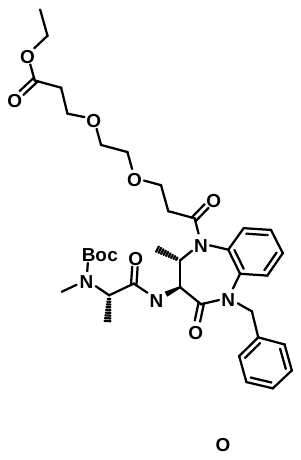

To a solution of tert-butyl ((R)-1-(((3R,4R)-1-benzyl-4-methyl-2-oxo-2,3,4,5-tetrahydro-1H-benzo[b][1,4]diazepin-3-yl)amino)-1-oxopropan-2-yl)(methyl)carbamate (**N**) (50.0 mg, 0.107 mmol, 1 eq.) at 0 ^o^C in DCM (1.1 mL) was added pyridine (42.4 mg, 0.536 mmol, 5 eq.) followed by ethyl 3-[2-(3-chloro-3-oxo-propoxy)ethoxy]propanoate (67.8 mg, 0.268 mmol, 2.5 eq.) dropwise. The reaction was stirred at 0 ºC for 1h, then allowed to warm to room temperature and stirred for an additional 15 h, then quenched by the addition of saturated NaHCO_3_ and extracted with DCM. The organic layers were combined, dried with sodium sulfate, and concentrated. The crude residue was purified by flash chromatography (3:1 iPrOAc:MeOH in heptanes) to afford the title compound (65.0 mg, 89%) of a yellow oil.

**Step 2: 3-(2-(3-((2R,3R)-5-benzyl-3-((R)-2-((tert-butoxycarbonyl)(methyl)amino)propanamido)-2-methyl-4-oxo-2,3,4,5-tetrahydro-1H-benzo[b][1,4]diazepin-1-yl)-3-oxopropoxy)ethoxy)propanoic acid (P)**

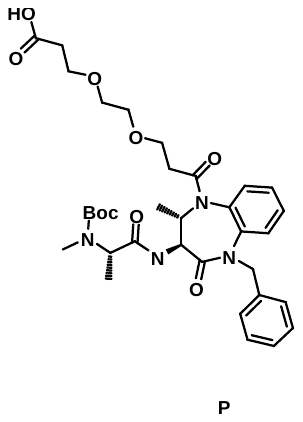

To a solution of ethyl 3-(2-(3-((2R,3R)-5-benzyl-3-((R)-2-((tert-butoxycarbonyl)(methyl)amino)propanamido)-2-methyl-4-oxo-2,3,4,5-tetrahydro-1H-benzo[b][1,4]diazepin-1-yl)-3-oxopropoxy)ethoxy)propanoate (**O**) (65.0 mg, 0.0952 mmol, 1 eq.) in THF (0.2 mL) and water (0.1 mL) was added LiOH (11.8 mg, 0.494 mmol, 5.2 eq.). The reaction was stirred at room temperature for 5 h. The solution was acidified to pH = 1 with 2N HCl. The solution was extracted with DCM 3x. The organic layers were combined, dried with Na_2_SO_4_ and concentrated. The crude product was carried over to the next step.

**Step 3: tert-butyl ((R)-1-(((3R,4R)-1-benzyl-5-(3-(2-(3-((2-(4-(1-(4-hydroxyphenyl)-2-phenylbut-1-en-1-yl)phenoxy)ethyl)(methyl)amino)-3-oxopropoxy)ethoxy)propanoyl)-4-methyl-2-oxo-2,3,4,5-tetrahydro-1H-benzo[b][1,4]diazepin-3-yl)amino)-1-oxopropan-2-yl)(methyl)carbamate (Q)**

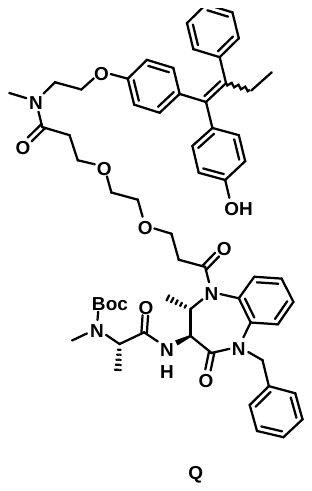

To a crude solution of 3-(2-(3-((2R,3R)-5-benzyl-3-((R)-2-((tert-butoxycarbonyl)(methyl)amino)propanamido)-2-methyl-4-oxo-2,3,4,5-tetrahydro-1H-benzo[b][1,4]diazepin-1-yl)-3-oxopropoxy)ethoxy)propanoic acid (**P**) (39.0 mg, 0.0596 mmol, 1 eq.) in 2-MeTHF (1.9 mL) was added DIPEA (24.6 mg, 0.190 mmol, 2 eq.) and HATU (40.6 mg, 0.105 mmol, 1.1 eq.) followed by 4-(1-(4-(2-(methylamino)ethoxy)phenyl)-2-phenylbut-1-en-1-yl)phenol (**D**) (23.4 mg, 0.0625 mmol, 1.05 eq.). The mixture was stirred at room temperature for 22h. Water was added and the solution was extracted 3x with EtOAc. The organic layers were combined then dried with sodium sulfate and concentrated to afford the crude title compound.

**Step 4:** **3-(2-(3-((2R,3R)-5-benzyl-2-methyl-3-((R)-2-(methylamino)propanamido)-4-oxo-2,3,4,5-tetrahydro-1H-benzo[b][1,4]diazepin-1-yl)-3-oxopropoxy)ethoxy)-N-(2-(4-(1-(4-hydroxyphenyl)-2-phenylbut-1-en-1-yl)phenoxy)ethyl)-N-methylpropanamide (GNE-9537)**

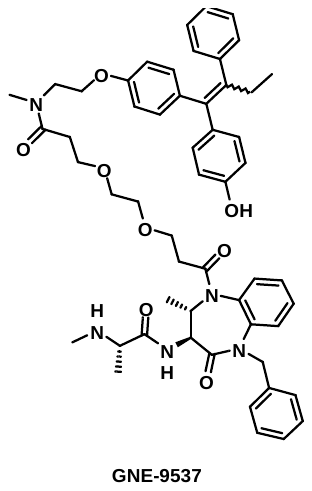

The crude tert-butyl ((R)-1-(((3R,4R)-1-benzyl-5-(3-(2-(3-((2-(4-(1-(4-hydroxyphenyl)-2-phenylbut-1-en-1-yl)phenoxy)ethyl)(methyl)amino)-3-oxopropoxy)ethoxy)propanoyl)-4-methyl-2-oxo-2,3,4,5-tetrahydro-1H-benzo[b][1,4]diazepin-3-yl)amino)-1-oxopropan-2-yl)(methyl)carbamate (**Q**) was dissolved in DCM (0.6 mL). TFA (0.16 mL) was added and the reaction was stirred at room temperature for 2h. The solution was concentrated under vacuum, diluted with saturated NaHCO_3_ and extracted 2x with DCM. The organic layers were combined, dried with sodium sulfated and concentrated. Purified by reverse-phase HPLC to afford the title compound (42 mg, 49% over 3-step) as a mixture of E/Z isomers. MS (ESI): [M+H]^+^ = 910.5. 1H NMR (400 MHz, DMSO) δ 9.40 (s, 0.5H), 9.15 – 9.08 (m, 1.5H), 8.76 – 8.71 (m, 3H), 7.75 (d, J = 8.2 Hz, 1H), 7.58 – 7.53 (m, 1H), 7.41 – 7.32 (m, 2H), 7.24 – 7.04 (m, 10H), 7.00 – 6.87 (m, 2H), 6.78 – 6.67 (m, 2H), 6.62 – 6.52 (m, 2H), 6.43 – 6.36 (m, 1H), 5.40 (d, J = 15.0 Hz, 1H), 4.98 (dd, J = 11.7, 6.2 Hz, 1H), 4.74 (d, J = 14.6 Hz, 1H), 4.15 – 4.01 (m, 2H), 3.97 – 3.79 (m, 2H), 3.73 – 3.54 (m, 2H), 3.35 – 3.28 (m, 2H), 3.12 (s, 1H), 3.04 (s, 1H), 2.95 (s, 1H), 2.87 (s, 1H), 2.80 (s, 1H), 2.73 – 2.61 (m, 1H), 2.42 – 2.30 (m, 2H), 2.03 – 1.87 (m, 1H), 1.42 (d, J = 6.9 Hz, 3H), 1.02 (d, J = 6.2 Hz, 3H), 0.88 – 0.79 (m, 4H).

**Synthesis of degrader GNE-6746**

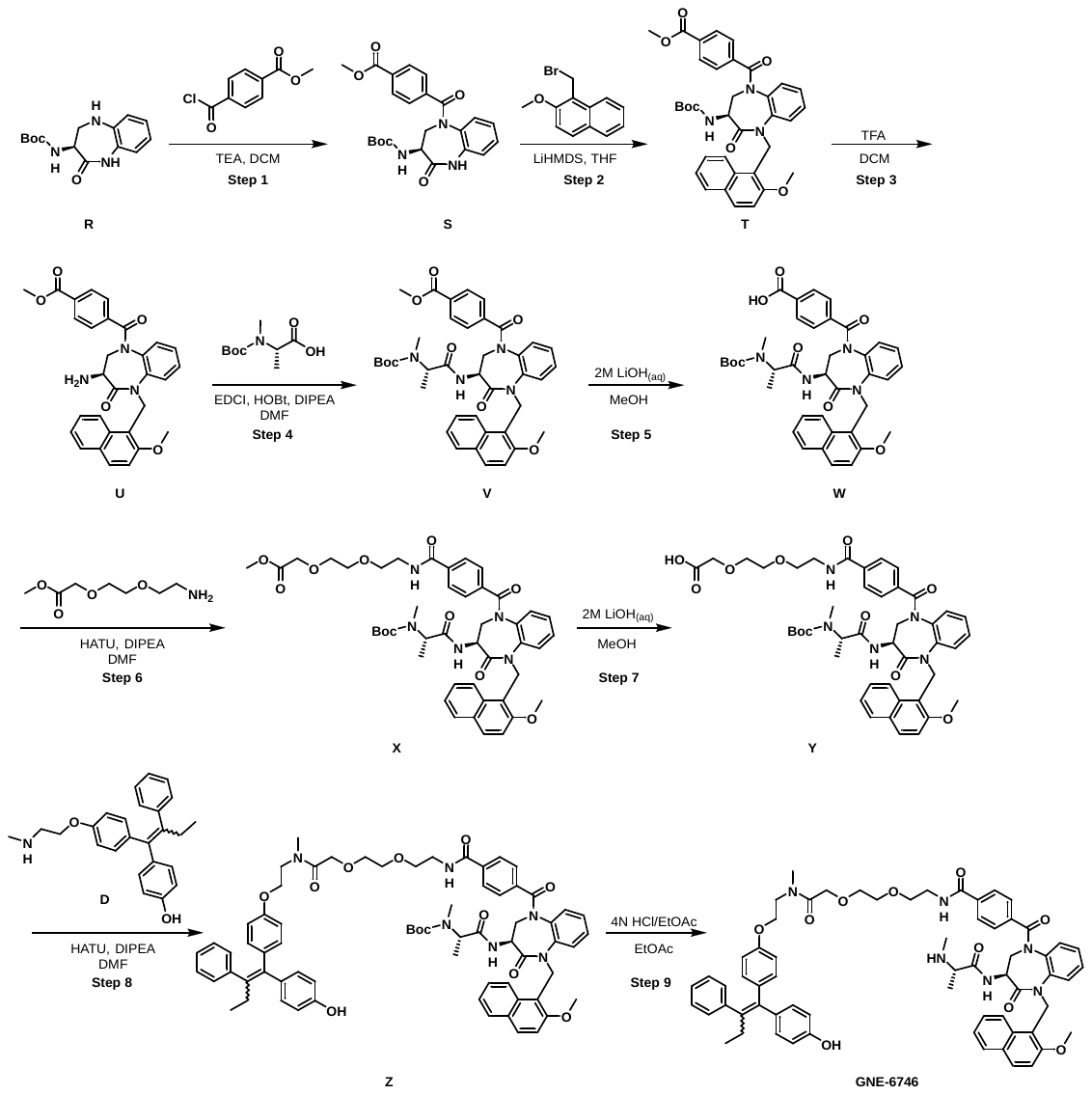

**Step 1: methyl (R)-4-(3-((tert-butoxycarbonyl)amino)-4-oxo-2,3,4,5-tetrahydro-1H-benzo[b][1,4]diazepine-1-carbonyl)benzoate (S)**

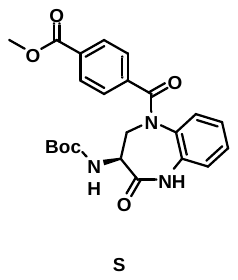

To a solution of tert-butyl (R)-(2-oxo-2,3,4,5-tetrahydro-1H-benzo[b][1,4]diazepin-3-yl)carbamate (**R**) (4.00 g, 14.4 mmol, 1 eq.) and methyl 4-chlorocarbonylbenzoate (3.15 g, 15.9 mmol, 1.1 eq.) in DCM (30 mL) was added TEA (4.38 g, 43.3 mmol, 3 eq) at 0 ^o^C .The reaction was stirred at room temperature for 1h. The reaction was quenched with water (10 mL), 1M HCl was added into the mixture until pH = 6 and the solution was extracted with DCM (100 mL), washed with water (30 mL*2) and dried over Na_2_SO_4_. The crude mixture purified by flash column chromatography eluting 20% EtOAc in petroleum ether to afford the title product (2.5g, 39% yield) as white solid. MS (ESI): [M+Na]^+^ = 462.1.

**Step 2: methyl (R)-4-(3-((tert-butoxycarbonyl)amino)-5-((2-methoxynaphthalen-1-yl)methyl)-4-oxo-2,3,4,5-tetrahydro-1H-benzo[b][1,4]diazepine-1-carbonyl)benzoate (T)**

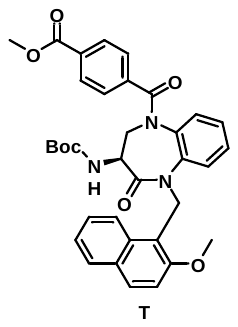

To a solution of methyl (R)-4-(3-((tert-butoxycarbonyl)amino)-4-oxo-2,3,4,5-tetrahydro-1H-benzo[b][1,4]diazepine-1-carbonyl)benzoate (**S**) (2.50 g, 5.69 mmol, 1 eq.) in THF (12 mL) was added 1M LiHMDS (8.54 mL, 8.54 mmol, 1.5 eq.) in THF at -78 ^o^C under N_2_. The mixture was stirred at -78 ^o^C for 30 min. Then, a solution of 1-(bromomethyl)-2-methoxy-naphthalene (1.57 g, 6.26 mmol, 1.1 eq.) in THF (3 mL) was added dropwise into the reaction mixture. The reaction was warm to room temperature and stirred for 16 h. The reaction was quenched with a saturated NH_4_Cl, extracted with EtOAc (100mL), washed with brine (30mL*2) and dried over Na_2_SO_4_. The crude residue was purified by flash column chromatography eluting 20% EtOAc in petroleum ether to afford the title product (2.5 g, 72% yield) as white solid. MS (ESI): [M+Na]^+^ = 632.1.

**Step 3: methyl (R)-4-(3-amino-5-((2-methoxynaphthalen-1-yl)methyl)-4-oxo-2,3,4,5-tetrahydro-1H-benzo[b][1,4]diazepine-1-carbonyl)benzoate (U)**

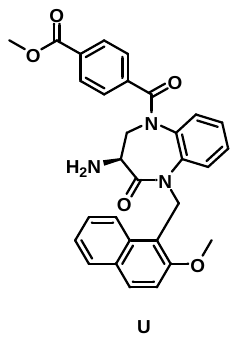

To a solution of methyl (R)-4-(3-((tert-butoxycarbonyl)amino)-5-((2-methoxynaphthalen-1-yl)methyl)-4-oxo-2,3,4,5-tetrahydro-1H-benzo[b][1,4]diazepine-1-carbonyl)benzoate (**T**) (1.5 g, 2.46 mmol, 1 eq.) in DCM (12 mL) was added TFA (4.0 mL) at 0 ^o^C. The reaction was allowed to warm up to room temperature and was stirred for 1h. The mixture was concentrated under vacuum to afford the title product (1.3 g) as yellow solid. The crude product was used in the next step without further purification. MS (ESI): [M+H]+ = 510.2.

**Step 4: methyl 4-((R)-3-((R)-2-((tert-butoxycarbonyl)(methyl)amino)propanamido)-5-((2-methoxynaphthalen-1-yl)methyl)-4-oxo-2,3,4,5-tetrahydro-1H-benzo[b][1,4]diazepine-1-carbonyl)benzoate (V)**

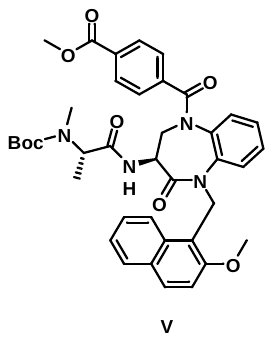

To a solution of methyl (R)-4-(3-amino-5-((2-methoxynaphthalen-1-yl)methyl)-4-oxo-2,3,4,5-tetrahydro-1H-benzo[b][1,4]diazepine-1-carbonyl)benzoate (**U**) (2.10 g, 4.12 mmol, 1 eq.) and (2S)-2-[tert-butoxycarbonyl(methyl)amino]propanoic acid (1.09 g, 5.36 mmol, 1.3 eq.) in DMF (15mL), was added DIPEA (1.60 g, 12.4 mmol, 3 eq.), HOBt (835 mg, 6.18 mmol, 1.5 eq.) and EDCI (1.19 g, 6.18 mmol, 1.5 eq.) at 0 ^o^C. The reaction was stirred at room temperature for 16 h. The mixture was added dropwise into water (100 mL) and the solid was collected by filtration. The crude product was purified by flash column chromatography eluting 20% EtOAc in petroleum ether to afford the title product (2.3 g, 80% yield) as white solid. MS (ESI): [M+Na]^+^ = 717.1.

**Step 5: 4-((R)-3-((R)-2-((tert-butoxycarbonyl)(methyl)amino)propanamido)-5-((2-methoxynaphthalen-1-yl)methyl)-4-oxo-2,3,4,5-tetrahydro-1H-benzo[b][1,4]diazepine-1-carbonyl)benzoic acid (W)**

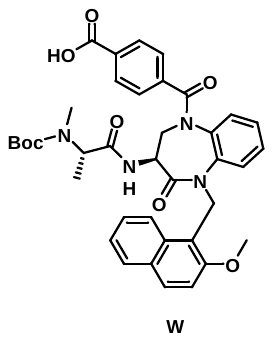

To a solution of methyl 4-((R)-3-((R)-2-((tert-butoxycarbonyl)(methyl)amino)propanamido)-5-((2-methoxynaphthalen-1-yl)methyl)-4-oxo-2,3,4,5-tetrahydro-1H-benzo[b][1,4]diazepine-1-carbonyl)benzoate (**V**) (400 mg, 0.580 mmol, 1 eq.) in MeOH (8 mL) was added 2M LiOH (0.86 mL, 1.73 mmol, 3 eq.). The reaction was stirred at 0 ^o^C for 16h. The reaction was concentrated under vacuum, then 2M HCl was added dropwise into the mixture until pH=6. The mixture was extracted with EtOAc (60 mL), washed with water (20mL*2), dried over Na_2_SO_4_ and concentrated under vacuum to afford the title product (390 mg) as white solid. The crude product was used in the next step without further purification. MS (ESI): [M+Na]+ = 703.4.

**Step 6: methyl 2-(2-(2-(4-((R)-3-((R)-2-((tert-butoxycarbonyl)(methyl)amino)propanamido)-5-((2-methoxynaphthalen-1-yl)methyl)-4-oxo-2,3,4,5-tetrahydro-1H-benzo[b][1,4]diazepine-1-carbonyl)benzamido)ethoxy)ethoxy)acetate (X)**

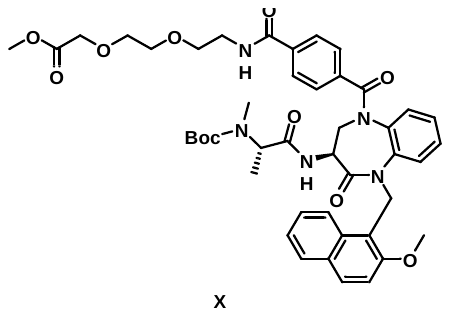

To a solution of 4-((R)-3-((R)-2-((tert-butoxycarbonyl)(methyl)amino)propanamido)-5-((2-methoxynaphthalen-1-yl)methyl)-4-oxo-2,3,4,5-tetrahydro-1H-benzo[b][1,4]diazepine-1-carbonyl)benzoic acid (**W**) (130 mg, 0.190 mmol, 1 eq.) in DMF (1.5mL) was added DIPEA (74.0 mg, 0.570 mmol, 3 eq.) and HATU (109 mg, 0.290 mmol, 1.5 eq.). The reaction was stirred at 20 ^o^C for 1h. The reaction was quenched with water (10mL), extracted with EtOAc (60 mL), washed with brine (20mL*4) and dried over Na_2_SO_4_. The crude residue was purified by flash column chromatography eluting 100% EtOAc in petroleum ether to afford the title product (110 mg, 68% yield) as yellow solid. MS (ESI): [M+H]+ = 840.5.

**Step 7: 2-(2-(2-(4-((R)-3-((R)-2-((tert-butoxycarbonyl)(methyl)amino)propanamido)-5-((2-methoxynaphthalen-1-yl)methyl)-4-oxo-2,3,4,5-tetrahydro-1H-benzo[b][1,4]diazepine-1-carbonyl)benzamido)ethoxy)ethoxy)acetic acid (Y)**

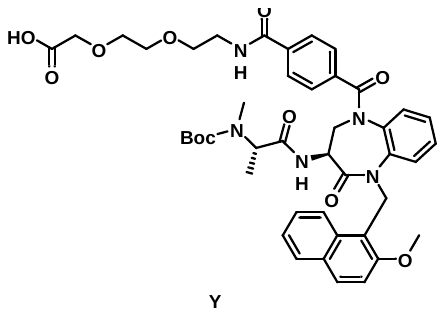

To a solution of methyl 2-(2-(2-(4-((R)-3-((R)-2-((tert-butoxycarbonyl)(methyl)amino)propanamido)-5-((2-methoxynaphthalen-1-yl)methyl)-4-oxo-2,3,4,5-tetrahydro-1H-benzo[b][1,4]diazepine-1-carbonyl)benzamido)ethoxy)ethoxy)acetate (**X**) (100 mg, 0.120 mmol, 1 eq.) in MeOH (3 mL) was added 2M LiOH (0.3 mL, 0.600 mmol, 5 eq.). The reaction was stirred at 20 ^o^C for 16h. The mixture was concentrated under vacuum and 2M HCl was added into the mixture until pH=6. The mixture was extracted with EtOAc (60 mL), washed with water (20 mL*2), dried over Na_2_SO_4_ and concentrated under vacuum to afford the title product (100 mg) as white solid. The crude product was used in the next step without further purification. MS (ESI): [M+Na]+ = 848.5.

**Step 8: tert-butyl ((R)-1-(((R)-5-(4-((2-(2-(2-((2-(4-(1-(4-hydroxyphenyl)-2-phenylbut-1-en-1-yl)phenoxy)ethyl)(methyl)amino)-2-oxoethoxy)ethoxy)ethyl)carbamoyl)benzoyl)-1-((2-methoxynaphthalen-1-yl)methyl)-2-oxo-2,3,4,5-tetrahydro-1H-benzo[b][1,4]diazepin-3-yl)amino)-1-oxopropan-2-yl)(methyl)carbamate (Z)**

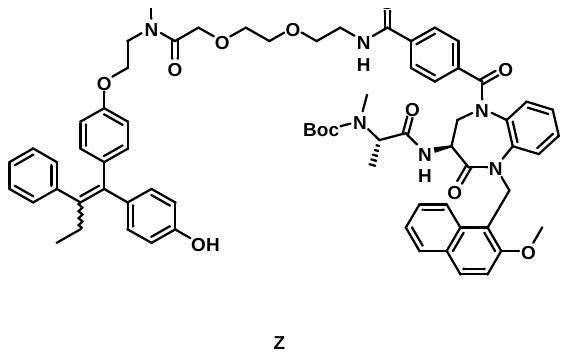

To a solution of 2-(2-(2-(4-((R)-3-((R)-2-((tert-butoxycarbonyl)(methyl)amino)propanamido)-5-((2-methoxynaphthalen-1-yl)methyl)-4-oxo-2,3,4,5-tetrahydro-1H-benzo[b][1,4]diazepine-1-carbonyl)benzamido)ethoxy)ethoxy)acetic acid (**Y**) (90.0 mg, 0.110 mmol, 1 eq.), 4-(1-(4-(2-(methylamino)ethoxy)phenyl)-2-phenylbut-1-en-1-yl)phenol (**D**) (48.8 mg, 0.130mmol, 1.2 eq.) and HATU (62.2 mg, 0.160 mmol, 1.5 eq.) in DMF (2 mL) was added DIPEA (42.3 mg, 0.330 mmol, 3 eq.) at 0 ^o^C. The reaction was stirred at room temperature for 1h. The reaction was work up by adding water (15 mL) into the mixture. The solid was collected by filtration and the crude product was purified by prep-HPLC (ACN/0.225% FA in water) to afford the title product (80 mg, 62% yield) as white solid. MS (ESI): [M+H]+ = 1182.2.

**Step 9: N-(2-(2-(2-((2-(4-(1-(4-hydroxyphenyl)-2-phenylbut-1-en-1-yl)phenoxy)ethyl)(methyl)amino)-2-oxoethoxy)ethoxy)ethyl)-4-((R)-5-((2-methoxynaphthalen-1-yl)methyl)-3-((R)-2-(methylamino)propanamido)-4-oxo-2,3,4,5-tetrahydro-1H-benzo[b][1,4]diazepine-1-carbonyl)benzamide (GNE-6746)**

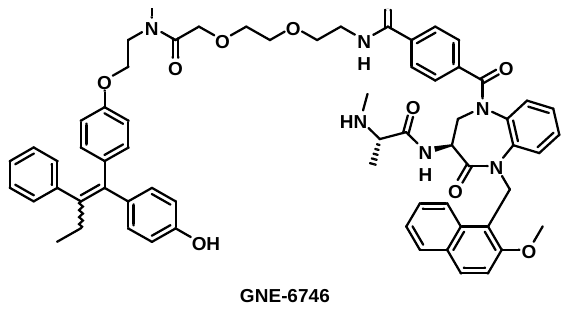

To a solution of tert-butyl ((R)-1-(((R)-5-(4-((2-(2-(2-((2-(4-(1-(4-hydroxyphenyl)-2-phenylbut-1-en-1-yl)phenoxy)ethyl)(methyl)amino)-2-oxoethoxy)ethoxy)ethyl)carbamoyl)benzoyl)-1-((2-methoxynaphthalen-1-yl)methyl)-2-oxo-2,3,4,5-tetrahydro-1H-benzo[b][1,4]diazepin-3-yl)amino)-1-oxopropan-2-yl)(methyl)carbamate (**Z**) (70.0 mg, 0.0600 mmol, 1 eq.) in EtOAc (2 mL) was added 4M HCl in EtOAc (3.mL) at 0 ^o^C. The reaction was stirred at room temperature for 1h. The reaction mixture was concentrated under vacuum and purified by prep-HPLC (ACN/0.225% FA in water) to afford the title product (19.5 mg, 29% yield) as white solid. MS (ESI): [M+H]+ = 1081.6. 1H NMR (400 MHz, Methanol-d4) δ 8.51 (brs, 1 H), 8.12 (d, J = 8.8 Hz, 1 H), 7.85- 7.87 (m, 2 H), 7.55 - 7.60 (m, 1 H), 7.24 - 7.42 (m, 4 H), 6.97 - 7.17 (m, 10 H), 6.86 - 6.90 (m, 1 H), 6.71 - 6.81 (m, 3 H), 6.63 (d, J = 8.4 Hz, 1 H), 6.50-6.53 (m, 1 H), 6.37-6.43 (m, 2 H), 6.23 (d, J = 15 Hz, 1 H), 5.82 (d, J = 7.9 Hz, 2 H), 5.54 (d, J = 15 Hz, 1 H), 4.74- 4.76 (m, 1 H), 4.35 - 4.45 (m, 2 H), 4.27 (d, J = 19 Hz, 1 H), 4.11 - 4.18 (m, 2 H), 3.94 - 3.98 (m, 3 H), 3.85 - 3.91 (m, 1 H), 3.54 - 3.82 (m, 10 H), 3.48-3.49 (m, 2 H), 3.06 (s, 1 H), 2.90 - 2.99 (m, 2 H), 2.60 (d, J = 4.0 Hz, 3 H), 2.42 - 2.51 (m, 2 H), 1.56 (d, J = 5.3 Hz, 3 H), 0.84 - 0.92 (m, 3 H).

**Synthesis of degrader GNE-9297**

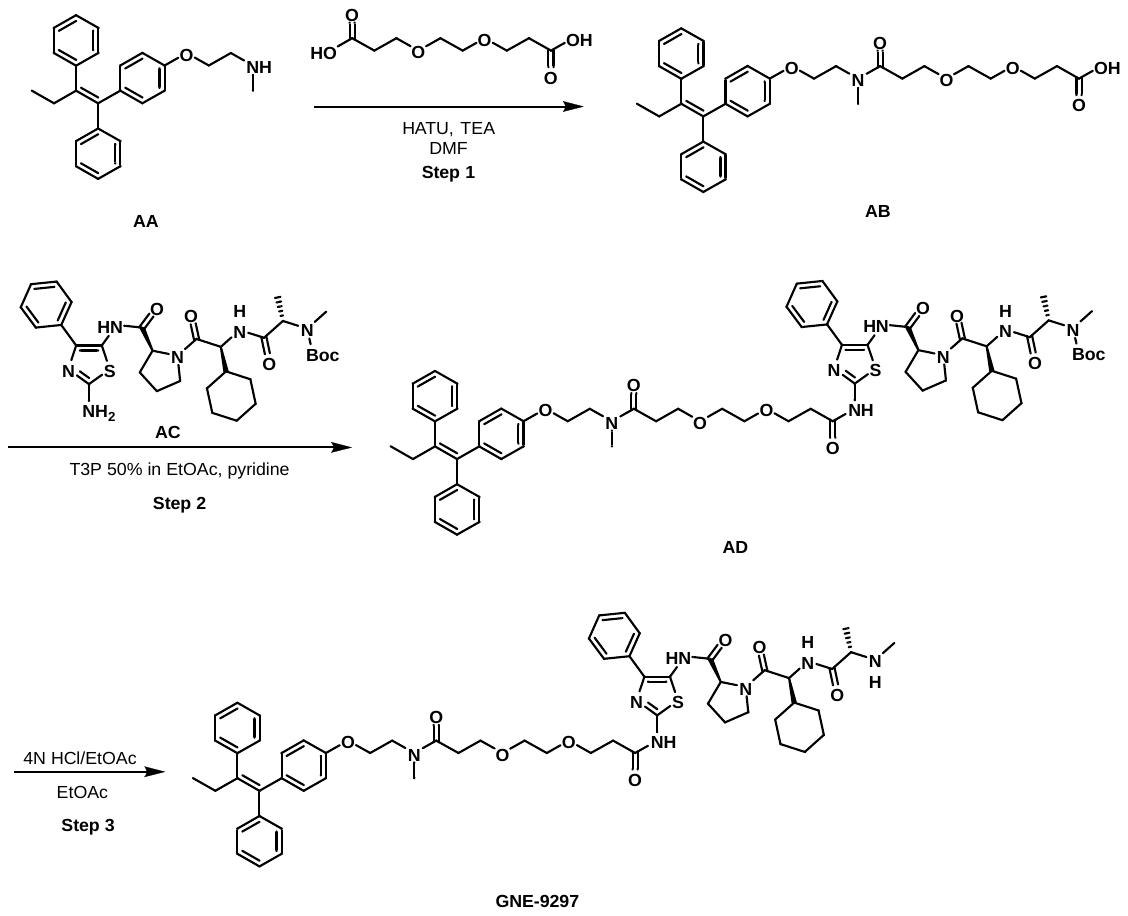

**Step 1:** **(Z)-3-(2-(3-((2-(4-(1,2-diphenylbut-1-en-1-yl)phenoxy)ethyl)(methyl)amino)-3-oxopropoxy)ethoxy)propanoic acid (AB)**

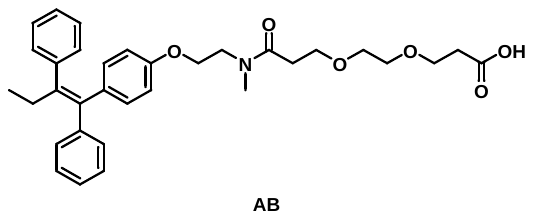

A mixture of 3-[2-(2-carboxyethoxy)ethoxy]propanoic acid (184 mg, 0.890 mmol, 1.1 eq.) and TEA (247 mg, 2.44 mmol, 3 eq.) in DMF (10 mL) was added HATU (340 mg, 0.890 mmol, 1.1 eq) and 2-[4-[(Z)-1,2-diphenylbut-1-enyl]phenoxy]-N-methyl-ethanamine hydrochloride (320 mg, 0.810 mmol, 1 eq.) at 20 ^o^C. The mixture was stirred for1 h. The reaction was quenched by addition of water (20 mL) and extracted with EtOAc (30 mL*3). The organic layer was washed with brine (30 mL) and concentrated. The residue was purified byf lash column chromatography (0-8% MeOH/DCM) to afford the title compound (210 mg, 47% yield) as a brown solid. MS (ESI): [M+H]+ = 546.3.

**Step 2: tert-butyl ((R)-1-(((R)-1-cyclohexyl-2-((R)-2-((2-(3-(2-(3-((2-(4-((Z)-1,2-diphenylbut-1-en-1-yl)phenoxy)ethyl)(methyl)amino)-3-oxopropoxy)ethoxy)propanamido)-4-phenylthiazol-5-yl)carbamoyl)pyrrolidin-1-yl)-2-oxoethyl)amino)-1-oxopropan-2-yl)(methyl)carbamate (AD)**

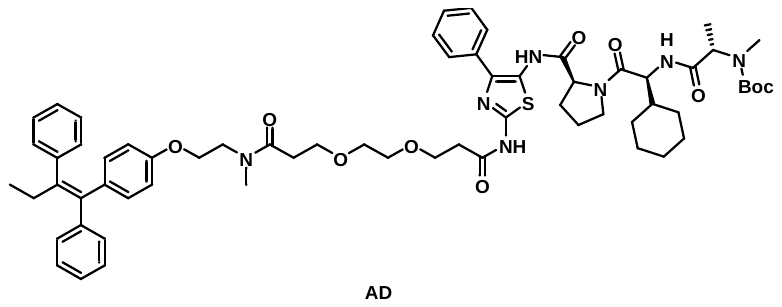

To a solution of (Z)-3-(2-(3-((2-(4-(1,2-diphenylbut-1-en-1-yl)phenoxy)ethyl)(methyl)amino)-3-oxopropoxy)ethoxy)propanoic acid (**AB**) (150 mg, 0.275 mmol, 1 eq.) and tert-butyl ((R)-1-(((R)-2-((R)-2-((2-amino-4-phenylthiazol-5-yl)carbamoyl)pyrrolidin-1-yl)-1-cyclohexyl-2-oxoethyl)amino)-1-oxopropan-2-yl)(methyl)carbamate (**AC**) (186 mg, 0.303 mmol, 1.1 eq) in pyridine (16 mL) was added T3P 50% in EtOAc (0.3 mL, 0.481 mmol, 1.75 eq.) at 20 ^o^C . The mixture was stirred for 12 h. The reaction was quenched by adding water (10 mL), extracted with EtOAc (30 mL*2) and washed with brine (20 mL). The organic layer was concentrated to give crude product, which was purified by prep-TLC (8% MeOH/DCM) to afford the title compound (120 mg, 38% yield) as a light yellow oil. MS (ESI): [M+H]+ = 1140.7.

**Step 3: (R)-1-((R)-2-cyclohexyl-2-((R)-2-(methylamino)propanamido)acetyl)-N-(2-(3-(2-(3-((2-(4-((Z)-1,2-diphenylbut-1-en-1-yl)phenoxy)ethyl)(methyl)amino)-3-oxopropoxy)ethoxy)propanamido)-4-phenylthiazol-5-yl)pyrrolidine-2-carboxamide (GNE-9297)**

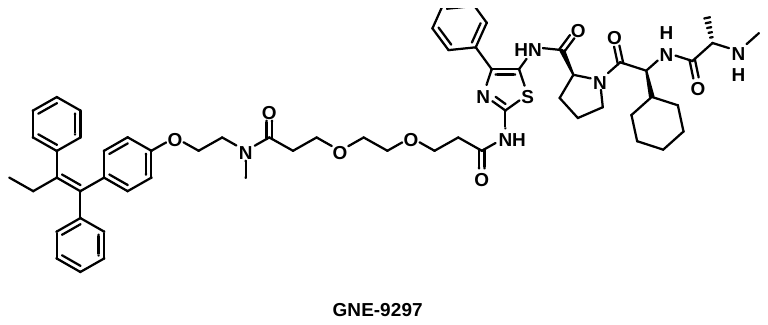

To a solution of tert-butyl ((R)-1-(((R)-1-cyclohexyl-2-((R)-2-((2-(3-(2-(3-((2-(4-((Z)-1,2-diphenylbut-1-en-1-yl)phenoxy)ethyl)(methyl)amino)-3-oxopropoxy)ethoxy)propanamido)-4-phenylthiazol-5-yl)carbamoyl)pyrrolidin-1-yl)-2-oxoethyl)amino)-1-oxopropan-2-yl)(methyl)carbamate (**AD**) (120 mg, 0.110 mmol, 1 eq.) in EtOAc (5 mL) was added 4M HCl in EtOAc (2 mL) at 20 ^o^C. The mixture was stirred for 1 h. The mixture was concentrated to give crude product, which was purified by prep-HPLC (35-65% ACN/0.225% FA in water) to afford the title compound (40 mg, 34% yield) as an off white solid. MS (ESI): [M+H]+ = 1040.7. 1H NMR (400 MHz, Methanol-d4) δ 8.47 (br s, 1H), 7.80 - 7.64 (m, 2H), 7.38 (t, J = 7.6 Hz, 2H), 7.35 - 7.22 (m, 4H), 7.20 - 7.11 (m, 4H), 7.10 - 7.04 (m, 3H), 6.78 - 6.68 (m, 2H), 6.54 - 6.46 (m, 2H), 4.61 - 4.46 (m, 2H), 3.99 - 3.84 (m, 3H), 3.79 - 3.57 (m, 8H), 3.57 - 3.46 (m, 4H), 2.98 (s, 1.5H), 2.86 (s, 1.5H), 2.70 - 2.59 (m, 3H), 2.58 - 2.50 (m, 4H), 2.41 (q, J = 7.6 Hz, 2H), 2.22 - 1.88 (m, 4H), 1.86 - 1.55 (m, 6H), 1.41 (d, J = 6.8 Hz, 3H), 1.30 - 0.96 (m, 5H), 0.87 (t, J = 7.6 Hz, 3H).

**Synthesis of degrader GNE-4863**

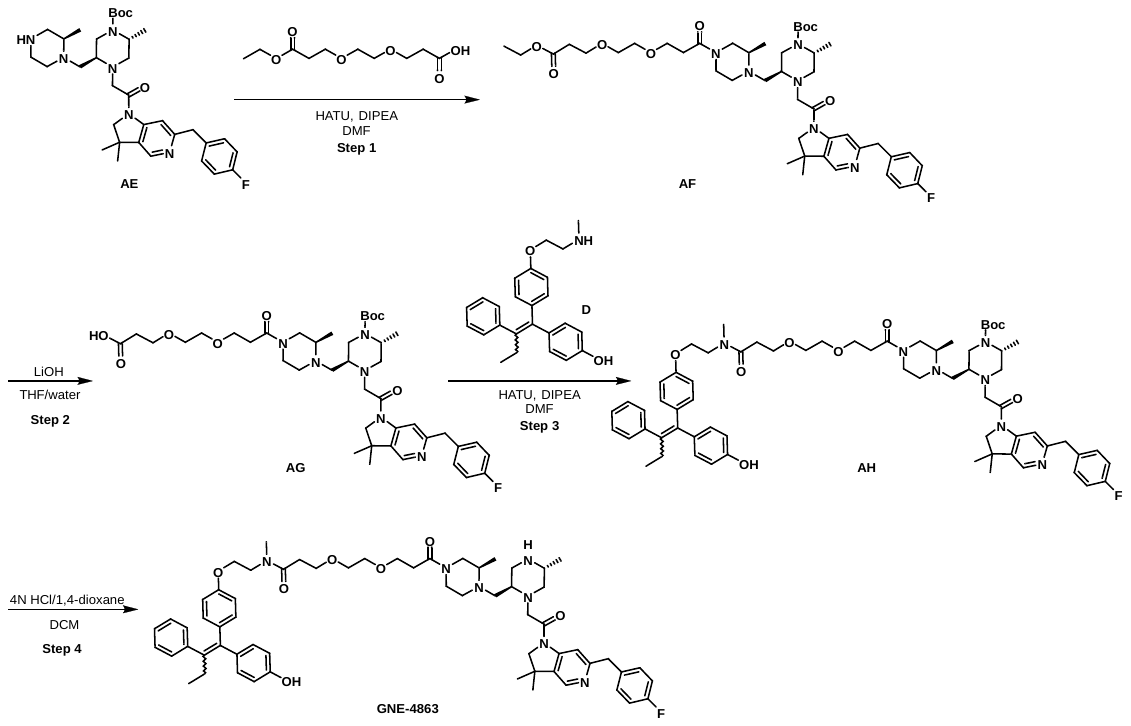

**Step 1: tert-butyl (2R,5S)-5-(((R)-4-(3-(2-(3-ethoxy-3-oxopropoxy)ethoxy)propanoyl)-2-methylpiperazin-1-yl)methyl)-4-(2-(6-(4-fluorobenzyl)-3,3-dimethyl-2,3-dihydro-1H-pyrrolo[3,2-c]pyridin-1-yl)-2-oxoethyl)-2-methylpiperazine-1-carboxylate (AF)**

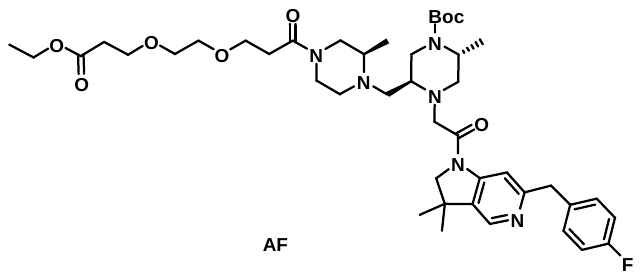

A solution of tert-butyl (2R,5S)-4-(2-(6-(4-fluorobenzyl)-3,3-dimethyl-2,3-dihydro-1H-pyrrolo[3,2-c]pyridin-1-yl)-2-oxoethyl)-2-methyl-5-(((R)-2-methylpiperazin-1-yl)methyl)piperazine-1-carboxylate (**AE**) (200 mg, 0.330 mmol, 1 eq.), 3-(2-(3-ethoxy-3-oxopropoxy)ethoxy)propanoic acid (92.4 mg, 0.390 mmol, 1.2 eq.) and DIEA (127 mg, 0.990 mmol, 3 eq.) in DMF (3 mL) was stirred at 25 ^o^C for 1 min. HATU (125 mg, 0.330 mmol, 1 eq.) was added and the mixture was stirred at 25 ^o^C for 1 h. The reaction mixture was purified by reverse phase flash chromatography ( 0-100% ACN/H_2_O) to afford the title compound (189 mg, 70% yield) as a white solid. MS (ESI): [M+H]+ = 825.6.

**Step 2: 3-(2-(3-((R)-4-(((2S,5R)-4-(tert-butoxycarbonyl)-1-(2-(6-(4-fluorobenzyl)-3,3-dimethyl-2,3-dihydro-1H-pyrrolo[3,2-c]pyridin-1-yl)-2-oxoethyl)-5-methylpiperazin-2-yl)methyl)-3-methylpiperazin-1-yl)-3-oxopropoxy)ethoxy)propanoic acid (AG)**

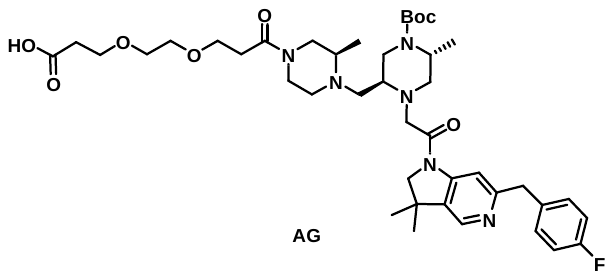

A solution of tert-butyl (2R,5S)-5-(((R)-4-(3-(2-(3-ethoxy-3-oxopropoxy)ethoxy)propanoyl)-2-methylpiperazin-1-yl)methyl)-4-(2-(6-(4-fluorobenzyl)-3,3-dimethyl-2,3-dihydro-1H-pyrrolo[3,2-c]pyridin-1-yl)-2-oxoethyl)-2-methylpiperazine-1-carboxylate (**AF**) (130 mg, 0.160 mmol, 1 eq.) in THF (4mL) was stirred at 25 ^o^C for 1 min. Then LiOH (10.4 mg, 0.470 mmol, 3 eq.) in water (2 mL) was added and the mixture was stirred at 25 ^o^C for 1 h. The solvent was concentrated under vacuum and the residue was purified by reverse phase flash chromatography (0-100% ACN/ 0.5% NH_4_HCO_3_ in water) to afford the title compound (109 mg, 87% yield) as a white solid. MS (ESI): [M+H]+ = 797.6.

**Step 3: tert-butyl (2R,5S)-4-(2-(6-(4-fluorobenzyl)-3,3-dimethyl-2,3-dihydro-1H-pyrrolo[3,2-c]pyridin-1-yl)-2-oxoethyl)-5-(((R)-4-(3-(2-(3-((2-(4-(1-(4-hydroxyphenyl)-2-phenylbut-1-en-1-yl)phenoxy)ethyl)(methyl)amino)-3-oxopropoxy)ethoxy)propanoyl)-2-methylpiperazin-1-yl)methyl)-2-methylpiperazine-1-carboxylate (AH)**

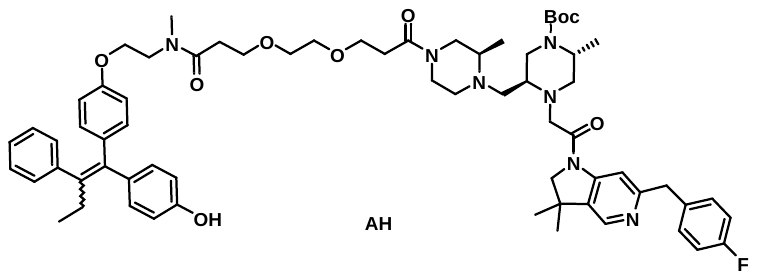

A solution of 3-(2-(3-((R)-4-(((2S,5R)-4-(tert-butoxycarbonyl)-1-(2-(6-(4-fluorobenzyl)-3,3-dimethyl-2,3-dihydro-1H-pyrrolo[3,2-c]pyridin-1-yl)-2-oxoethyl)-5-methylpiperazin-2-yl)methyl)-3-methylpiperazin-1-yl)-3-oxopropoxy)ethoxy)propanoic acid (**AG**) (109 mg, 0.140 mmol, 1 eq.), 4-(1-(4-(2-(methylamino)ethoxy)phenyl)-2-phenylbut-1-en-1-yl)phenol (**D**) (51.1 mg, 0.140 mmol, 1 eq.) and DIEA (52.9 mg, 0.410 mmol, 3 eq.) in DMF (2 mL) was stirred at 25 ^o^C for 5 min. Then HATU (57.2 mg, 0.150 mmol, 1.1 eq) was added and the mixture was stirred at 25 ^o^C for 1 h. The residue was purified by reverse phase flash chromatography (0-100% ACN/ 0.5% NH_4_HCO_3_ in water) to afford the title compound (130 mg, 83% yield) as a white solid. MS (ESI): [M+H]+ = 1152.8.

**Step 4: 3-(2-(3-((R)-4-(((2R,5R)-1-(2-(6-(4-fluorobenzyl)-3,3-dimethyl-2,3-dihydro-1H-pyrrolo[3,2-c]pyridin-1-yl)-2-oxoethyl)-5-methylpiperazin-2-yl)methyl)-3-methylpiperazin-1-yl)-3-oxopropoxy)ethoxy)-N-(2-(4-(1-(4-hydroxyphenyl)-2-phenylbut-1-en-1-yl)phenoxy)ethyl)-N-methylpropanamide (GNE-4863)**

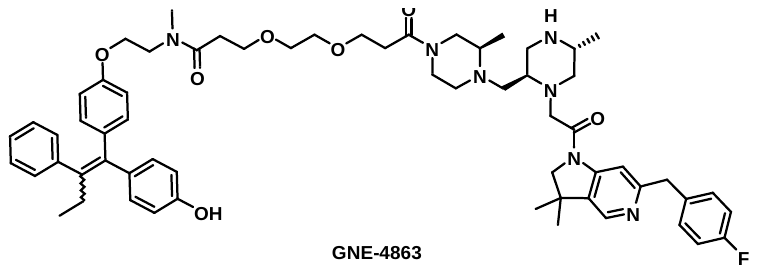

A solution of tert-butyl (2R,5S)-4-(2-(6-(4-fluorobenzyl)-3,3-dimethyl-2,3-dihydro-1H-pyrrolo[3,2-c]pyridin-1-yl)-2-oxoethyl)-5-(((R)-4-(3-(2-(3-((2-(4-(1-(4-hydroxyphenyl)-2-phenylbut-1-en-1-yl)phenoxy)ethyl)(methyl)amino)-3-oxopropoxy)ethoxy)propanoyl)-2-methylpiperazin-1-yl)methyl)-2-methylpiperazine-1-carboxylate (**AH**) (130 mg, 0.110 mmol, 1 eq.) in DCM (2 mL) was stirred at 25 ^o^C for 1 min. Then 4N HCl/1,4-dioxane (2 mL) was added and the mixture was stirred at 25 ^o^C for 30 min. The solvent was concentrated under vacuum. The residue was purified by reverse phase flash chromatography (5-100% ACN/ 0.5% NH_4_HCO_3_ in water) to afford the title compound (101 mg, 85% yield) as a white solid. MS (ESI): [M+H]+ = 1052.5. 1H NMR (400 MHz, DMSO-d6) δ 9.52 - 9.14 (br m, 1H), 8.28 (s, 1H), 7.28 - 7.26 (m, 2H), 7.17 - 7.12 (m, 2H), 7.11 - 7.07 (m, 6H), 6.96 - 6.91 (m, 2H), 6.75 - 6.67 (m, 2H), 6.58 - 6.54 (m, 2H), 6.38 (d, J = 8 Hz, 2H), 4.14 - 3.86 (m, 5H), 3.84 - 3.69 (m, 3H), 3.67 - 3.52 (m, 8H), 3.45 - 3.36 (m, 4H), 3.06 - 2.87 (m, 4H), 2.80 - 2.67 (m, 5H), 2.66 - 2.52 (m, 4H), 2.43 - 2.32 (m, 4H), 2.28 - 2.12 (m, 2H), 1.97 - 1.72 (m, 3H), 1.30 (d, J = 4.0 Hz, 6H), 0.97 - 0.88 (m, 3H), 0.84 - 0.79 (m, 6H).

**Synthesis of degrader GNE-8444 and GNE-8446**

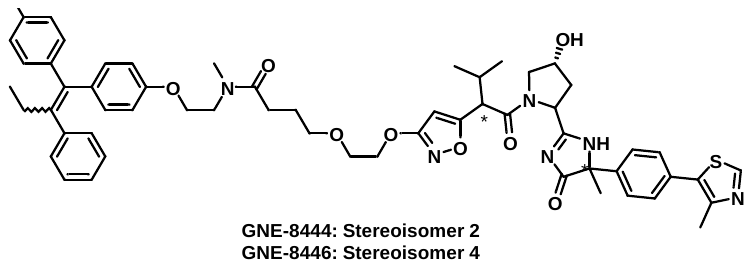

Synthesis of degrader GNE-8444 and GNE-8446 was reported here (Ex. 1003.1-4): WO2019084026A1

**Synthesis of ligand GNE-4402**

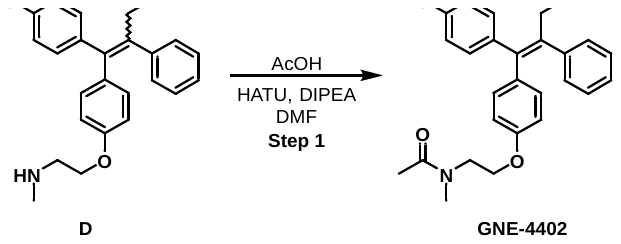

**Step 1: (Z)-N-(2-(4-(1-(4-hydroxyphenyl)-2-phenylbut-1-en-1-yl)phenoxy)ethyl)-N-methylacetamide**

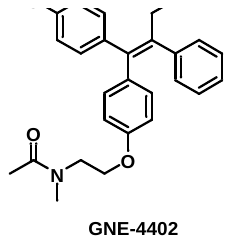

To a solution of acetic acid (7.24 µL, 7.590 mg, 0.126 mmol, 1 eq.) in 2-MeTHF (1.3 mL) was added HATU (53.9 mg, 0.139 mmol, 1.1 eq.) and DIPEA (32.7 mg, 0.253 mmol, 2 eq.). The mixture was stirred at room temperature for 30 min, then a solution of 4-(1-(4-(2-(methylamino)ethoxy)phenyl)-2-phenylbut-1-en-1-yl)phenol (**D**) (49.6 mg, 0.133 mmol, 1.05 eq.) in 2-MeTHF (0.5 mL) was added. The mixture was stirred at room temperature for 22h. Water was added and the solution was extracted 3x with iPrOAc. The organic layers were combined then dried with sodium sulfate and concentrated in vacuum. The crude product was purified by chiral reverse-phase chromatography to afford the Z isomer (18 mg, 34% yield). MS (ESI): [M+H]+ = 416.2. NMR showed a 1:1 mixture of 2 amide rotamers: 1H NMR (400 MHz, DMSO) δ 9.39 (s, 1H), 7.21 – 7.14 (m, 2H), 7.14 – 7.05 (m, 3H), 6.97 (d, J = 8.3 Hz, 2H), 6.81 – 6.66 (m, 4H), 6.63 – 6.54 (m, 2H), 3.98 (t, J = 5.3 Hz, 1H), 3.90 (t, J = 5.8 Hz, 1H), 3.58 (t, J = 5.3 Hz, 1H), 3.52 (t, J = 5.8 Hz, 1H), 2.98 (s, 1.5H), 2.80 (s, 1.5H), 2.53 (s, 4H), 2.45 – 2.36 (m, 2H), 1.98 (s, 1.5H), 1.94 (s, 1.5H), 0.84 (t, J = 7.4 Hz, 3H).

**LSMC characterization of compounds in figure S1B:**

| **Compound ID** | **Structure** | **MW** | **MS (ESI): [M+H]+** |
| --- | --- | --- | --- |
| RG-7112_analog | 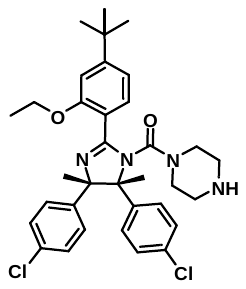 | 607.6 | 607.1 |
| GNE-5481 | 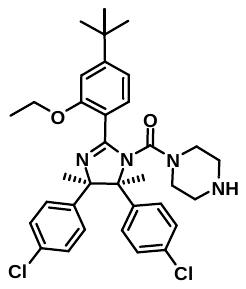 | 607.6 | 607.4 |
| GNE-3144 | 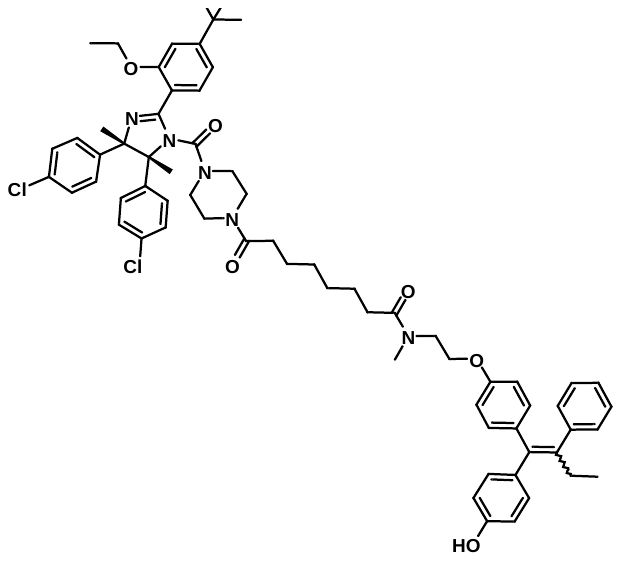 | 1119.3 | 1121.5 |
| GNE-3183 | 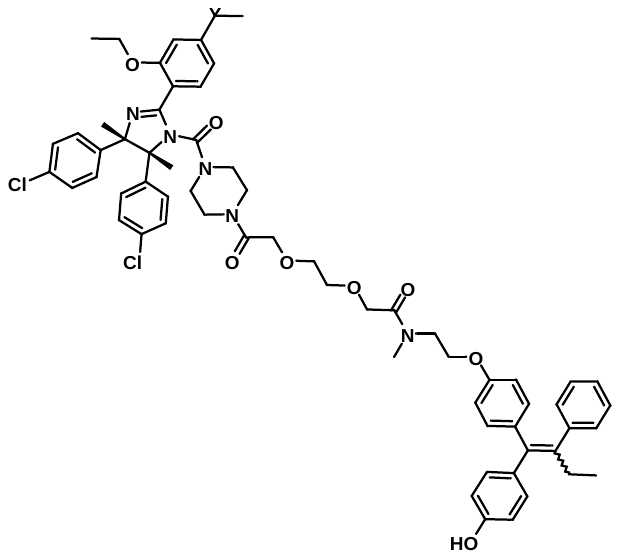 | 1123.2 | 1124.4 |
| GNE-3185 | 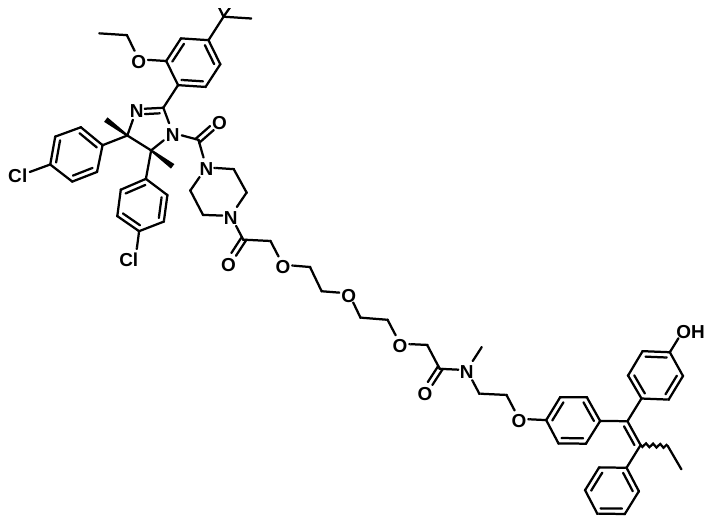 | 1167.3 | 1167.5 |
| GNE-0429 | 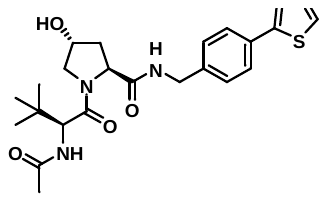 | 472.6 | 473.1 |
| GNE-7388 | 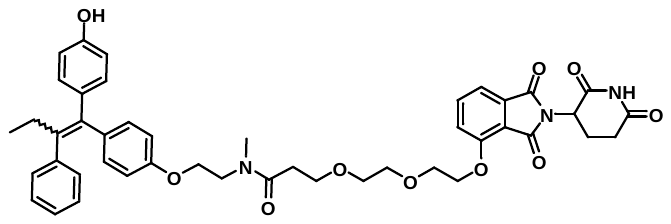 | 789.9 | 790.5 |

**LSMC characterization of compounds in figure 1C/table 1:**

| **Compound ID** | **Structure** | **MW** | **MS (ESI): [M+H]+** |
| --- | --- | --- | --- |
| endoxifen+linker.1 | 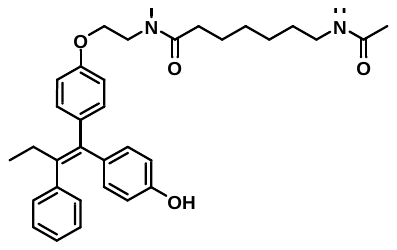 | 542.7 | 543.3 |
| endoxifen+linker.2 | 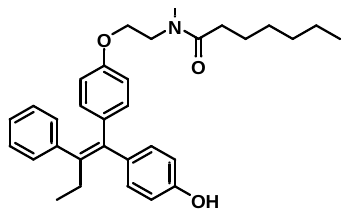 | 485.7 | 486.3 |
| endoxifen+linker.3 | 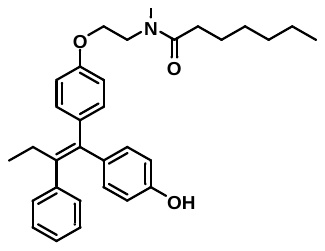 | 485.7 | 486.3 |
| GNE-8506 | 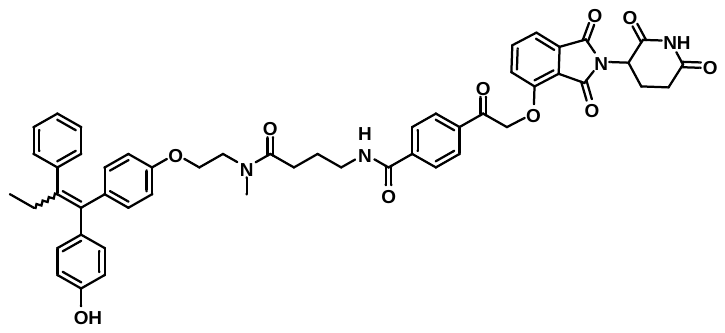 | 876.9 | 877.6 |
| GNE-8093 | 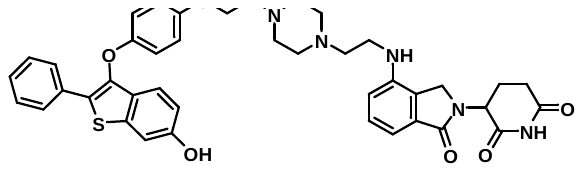 | 731.9 | 732.2 |
| GNE-2681 | 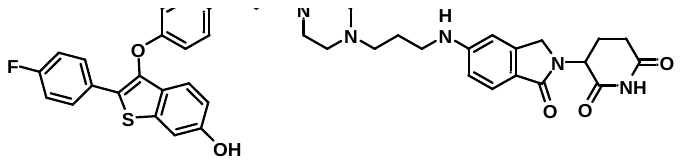 | 763.9 | 764.3 |
| GNE-3340 | 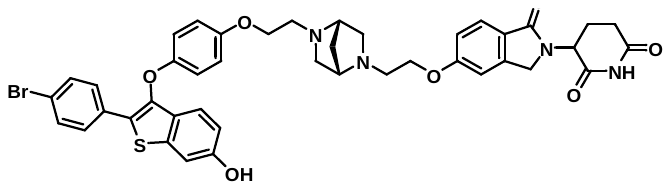 | 823.8 | 823.1 |
| XB2m54 analog | 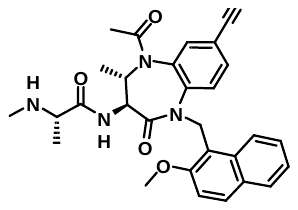 | 513.6 | 536.1 (M+Na) |
| GNE-5583 | 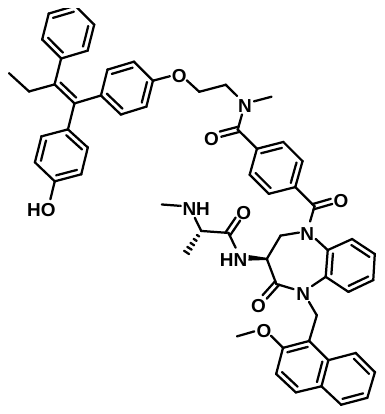 | 936.1 | 936.6 |
| GNE-0079 | 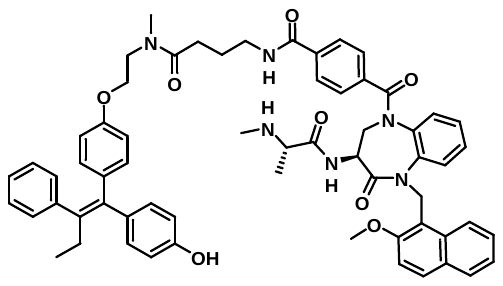 | 1021.2 | 1021.7 |
| GNE-0080 | 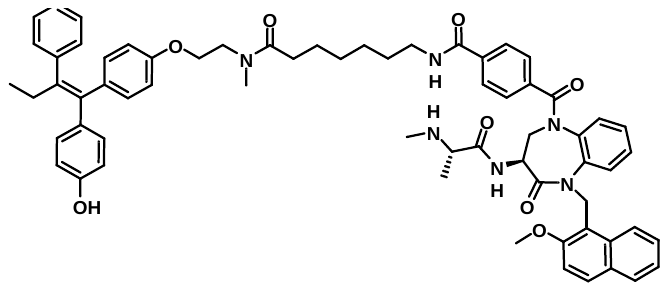 | 1063.3 | 1064.3 |
| GNE-0081 | 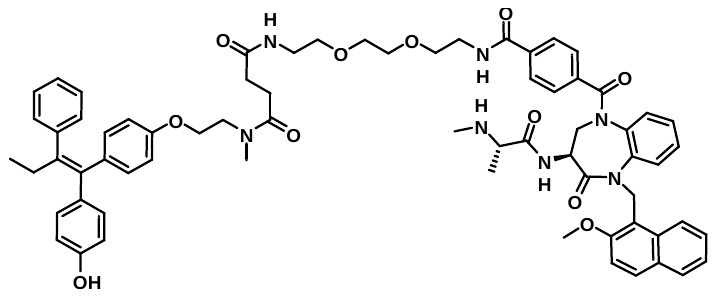 | 1166.4 | 584.1 ([M+H]/2) |
| GNE-0993 | 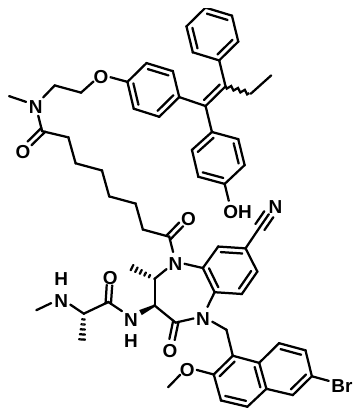 | 1062.1 | 1061.5 |
| GNE-0994 | 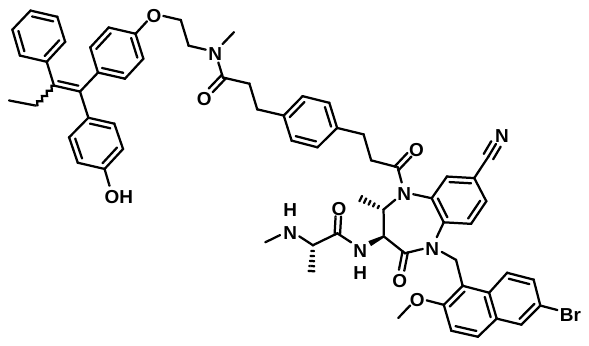 | 1110.2 | 1109.5 |
| GNE-4182 | 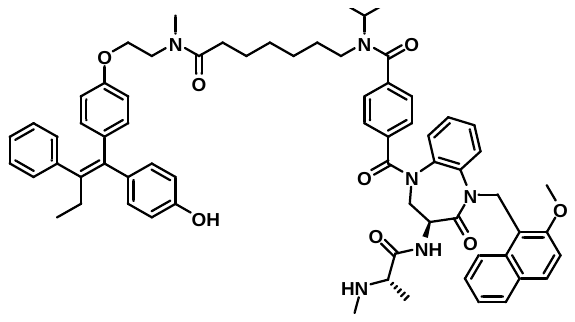 | 1105.4 | 1106.2 |
| GNE-6734 | 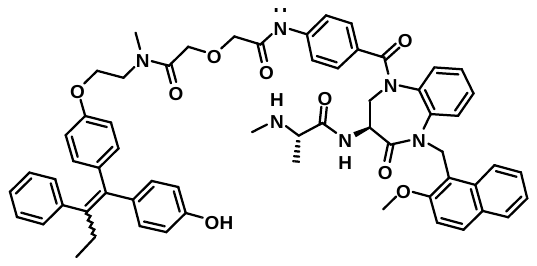 | 1023.2 | 1024.3 |
| GNE-6735 | 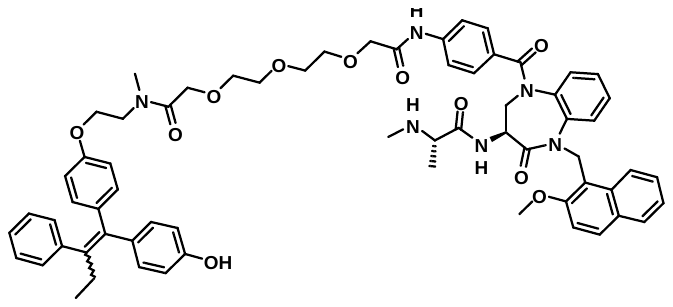 | 1111.3 | 1112.3 |
| GNE-6736 | 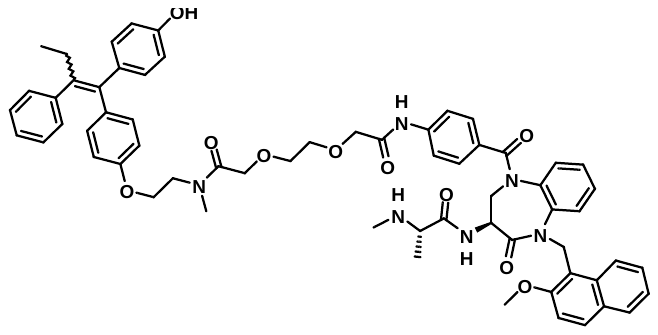 | 1067.3 | 1068.3 |
| GNE-7530 | 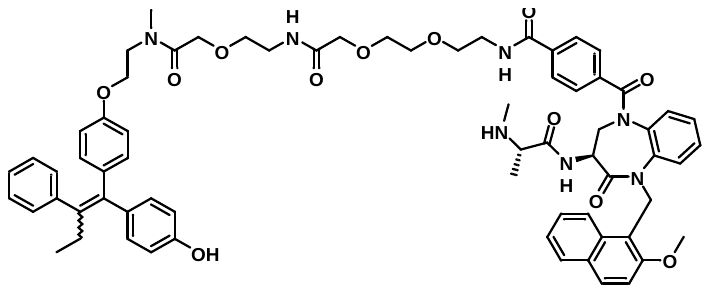 | 1182.3 | 1182.9 |
| GNE-7648 | 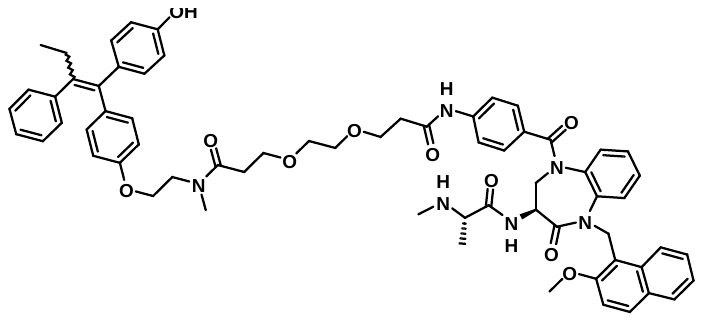 | 1095.3 | 1096.3 |
| GNE-0356 | 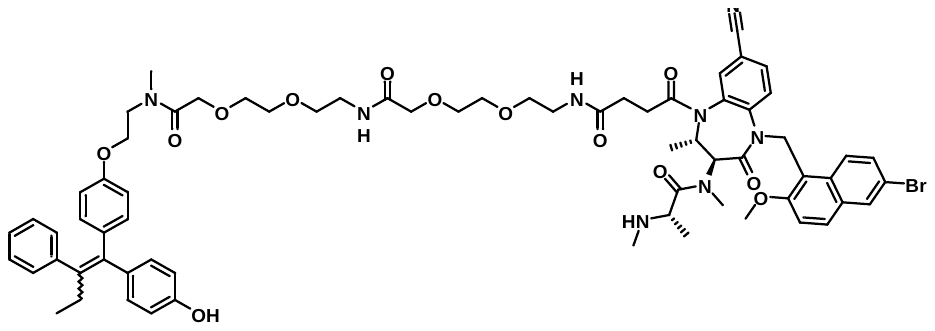 | 1310.4 | 1309.5 |
| GNE-4081 | 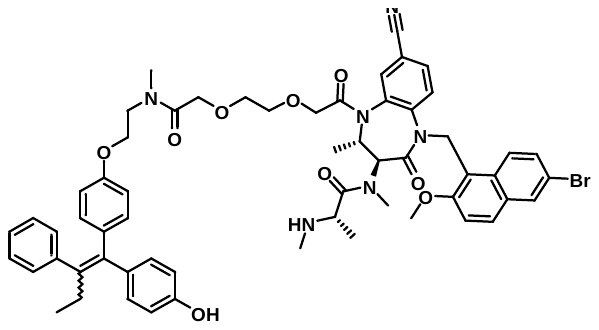 | 1080.1 | 1080.4 |
| GNE-4082 | 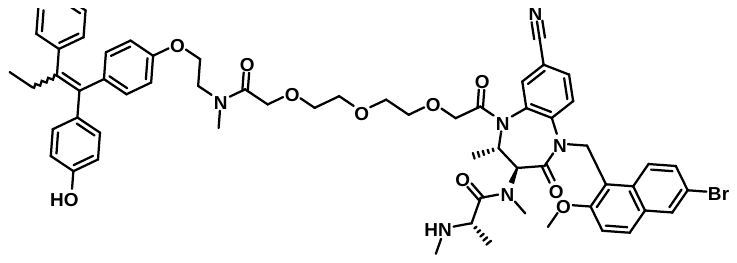 | 1124.1 | 1125.4 |
| GNE-8946 | 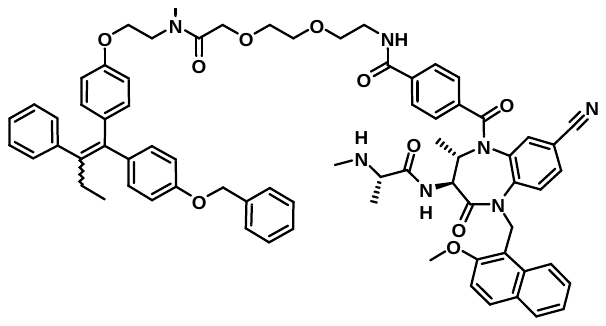 | 1210.4 | 1211.4 |
| GDC-0152 analog.1 | 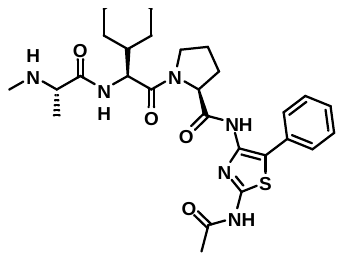 | 554.7 | 555.3 |
| GDC-0152 analog.2 | 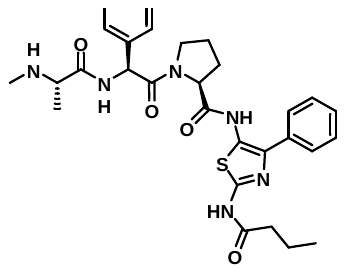 | 576.7 | 577.3 |
| GDC-0152 analog.3 | 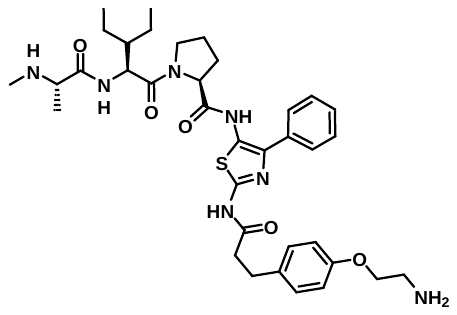 | 703.9 | 704.3 |
| GDC-0152 analog.4 | 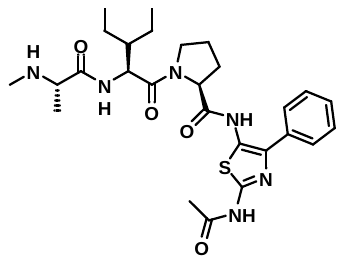 | 554.7 | 555.3 |
| GNE-8443 | 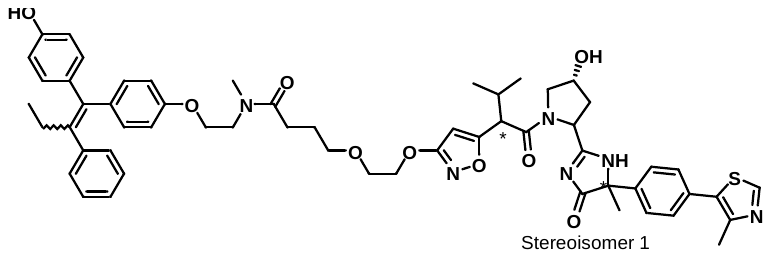 | 1009.2 | 1009.4 |
| GNE-8445 | 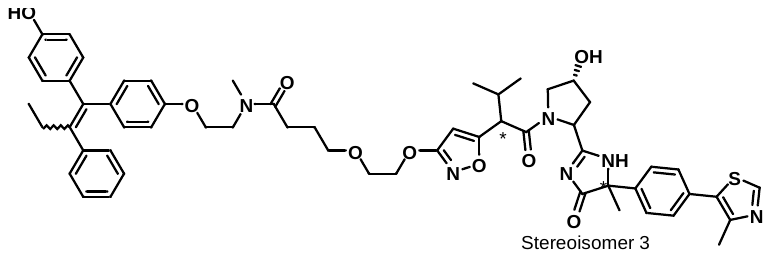 | 1009.2 | 1009.4 |
| GNE-1574 | 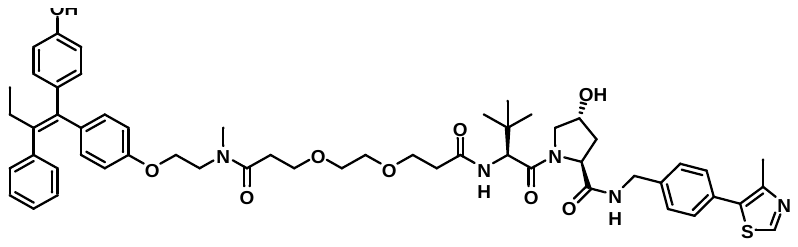 | 974.2 | 974.6 |
| GNE-2003 | 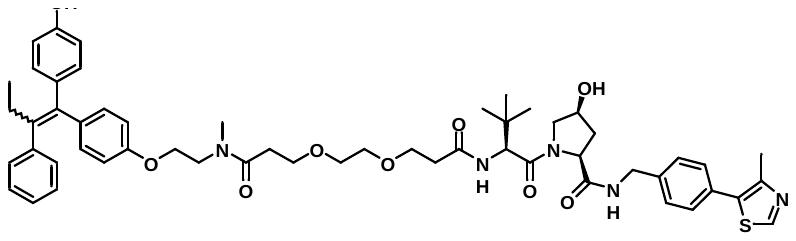 | 974.2 | 996.5 (M+Na) |
